# RNA polymerase II transcription with partially assembled TFIID complexes

**DOI:** 10.1101/2023.11.27.567046

**Authors:** Vincent Hisler, Paul Bardot, Dylane Detilleux, Matthieu Stierle, Emmanuel Garcia Sanchez, Claire Richard, Lynda Hadj Arab, Cynthia Ehrhard, Bastien Morlet, Yavor Hadzhiev, Matthieu Jung, Stéphanie Le Gras, Luc Négroni, Ferenc Müller, László Tora, Stéphane D. Vincent

**Affiliations:** Université de Strasbourg, IGBMC UMR 7104-UMR-S 1258, F-67400 Illkirch, France; CNRS, UMR 7104, F-67400 Illkirch, France; Inserm, UMR-S 1258, F-67400 Illkirch, France; IGBMC, Institut de Génétique et de Biologie Moléculaire et Cellulaire, F-67400 Illkirch, France; Proteomics platform; GenomEast; Institute of Cancer and Genomic Sciences, College of Medical and Dental Sciences, University of Birmingham, B152TT, Birmingham, UK

**Keywords:** TFIID, TATA binding protein (TBP), TBP-associated factors (TAFs), TAF7, TAF10, RNA polymerase II, nascent transcription, mass spectrometry, mouse, embryonic stem cells (ESCs), assembly

## Abstract

The recognition of core promoter sequences by the general transcription factor TFIID is the first step in the process of RNA polymerase II (Pol II) transcription initiation. Metazoan holo-TFIID is composed of the TATA binding protein (TBP) and of 13 TBP associated factors (TAFs). Inducible *Taf7* knock out (KO) results in the formation of a Taf7-less TFIID complex, while *Taf10* KO leads to serious defects within the TFIID assembly pathway. Either TAF7 or TAF10 depletions correlate with the detected TAF occupancy changes at promoters, and with the distinct phenotype severities observed in mouse embryonic stem cells or mouse embryos. Surprisingly however, under either *Taf7* or *Taf10* deletion conditions, TBP is still associated to the chromatin, and no major changes are observed in nascent Pol II transcription. Thus, partially assembled TFIID complexes can sustain Pol II transcription initiation, but cannot replace holo-TFIID over several cell divisions and/or development.

## Introduction

RNA polymerase II (Pol II) is responsible for the transcription of all protein coding genes, as well as long non coding RNA and some short non coding RNA (reviewed in ^1^). Pol II transcription is finely regulated, allowing gene by gene variable levels of expression depending on the cellular context. As a result, dysfunction in Pol II transcription is associated with a wide range of pathologies such as cancer, metabolic or neural diseases. Pol II transcription is first regulated by the binding of specific transcription factors to enhancers. These transcription factors are able to recruit different classes of transcriptional co-activator complexes that modify the structure of the chromatin and lead to chromatin opening and/or creating anchor points for the recruitment of other complexes. The action of these co-activators is crucial to create a favorable context for transcription initiation. Pol II recruitment via the formation of the pre-initiation complex (PIC) on active promoters is the obligatory step for transcription initiation.

The PIC is composed of 6 general transcription factors (GTFs): TFIIA, TFIIB, TFIID, TFIIE, TFIIF and TFIIH, and Pol II (reviewed in ^1^). TFIID is the first GTF to bind to the promoter, initiating the nucleation of the PIC. In Metazoans, holo-TFIID is composed of the TATA-binding protein (TBP) and of 13 TBP-associated factors (TAFs), among which 6 are present in 2 copies ^11,12^ and is recruited to the promoter by multiple non-mutually exclusive mechanisms. The double bromodomain of TAF1 and the PHD domain of TAF3 allow TFIID interaction with epigenetic marks associated with open chromatin such as acetylated histones and H3K4me3, respectively ^2,3^. Interactions between some TAFs and different transcription factors have been reported, indicating that the TAFs could also facilitate enhancer/core promoter interactions ^4–7^. Importantly, TFIID interacts with DNA motifs such as the TATA-box via TBP ^8^ or the initiator element (Inr) via TAF1 and TAF2 ^9,10^.

Holo-TFIID is composed of three lobes, called A, B and C (Figure 1A) ^10,12^. Lobes A and B are characterized by the presence of TAFs sharing a conserved histone fold domain (HFD) that allows the formation of specific heterodimers. Lobe A is composed of 10 subunits: TAF5, TBP as well as the TAF4/TAF12, TAF6/TAF9, TAF11/13 and TAF3/TAF10 HF pairs. Lobe B contains 7 subunits: TAF5 and the TAF4/TAF12, TAF6/TAF9 and TAF8/TAF10 histone fold pairs ^10,12^. The HEAT domains of the 2 copies of TAF6, together with TAF1 and TAF8 C-terminal half connect these two lobes and, in association with TAF2, constitutes the lobe C ^10,12^.

**Figure 1:**
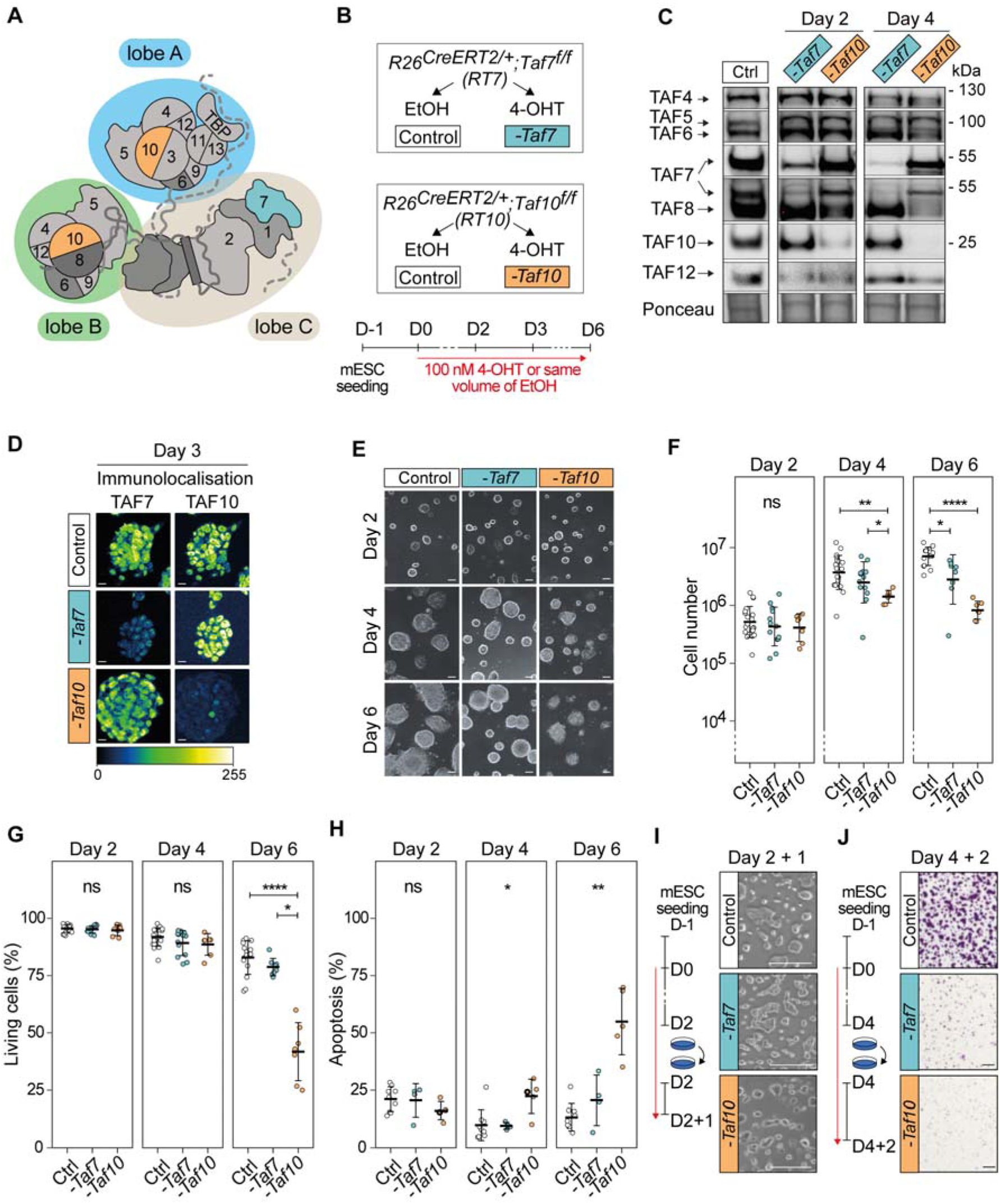
Phenotypic analysis of the conditional depletion of TAF7 or TAF10 in mESCs. **(A)** Schematic structure of TFIID. Lobe A is indicated in blue, lobe B in green and lobe C in beige. **(B)** Strategy of the induction of the deletion of *Taf7* (-*Taf7*) in *R26^CreERT2/+^;Taf7^f/f^*(*RT7*) and *Taf10* (-*Taf10*) in *R26^CreERT2/+^;Taf10^f/f^* (*RT10*) mESCs. Cells are plated at day (D) - 1, 100 nM 4-hydroxytamoxifen (4-OHT, depleted) or the same of volume of ethanol (control) were added at D0 and maintained until the day of the analysis. **(C)** Western blot analyses of TAF4, TAF5, TAF6, TAF7, TAF8, TAF10 and TAF12 proteins expression after *Taf7* (-*Taf7*) or *Taf10* (-*Taf10*) deletions after 2 and 4 days of 4-OHT treatment. As a control (Ctrl), *RT7* cells were treated 2 days with EtOH. The Ponceau staining is displayed at the bottom of the panel. **(D)** Immunolocalization of TAF7 and TAF10 in *RT7* and *RT10* cells treated for 3 days with 4-OHT. As a control, *RT10* cells were treated with EtOH for 3 days. Color scale (Green Fire Blue LUT scale) is indicated at the bottom. Scale bar; 15µm. **(E)** Monitoring of cell growth over time. *RT7* and *RT10* cells were treated over 6 days, and pictures were taken at D2, D4 and D6. Scale bar; 50 µm. **(F)** Log10 of the total number of cells at D2, D4 and D6 of treatment. **(G)** Percentage of living cells after 2, 4, and 6 days of treatment determined by Trypan blue staining. For (E, F), Ctrl: D2; N = 20, D4; n = 20, D6; n = 15, -*Taf7*: D2; n = 13, D4; n = 13, D6; n = 8, *-Taf10*: D2; n = 7, D4; n = 7, D6; n = 7 biological replicates. **(H)** Percentage of apoptotic cells after 2, 4 and 6 days of treatment determined by Annexin V and propidium iodide (PI) staining. For D2, D4 and D6, Ctrl; n = 9, -*Taf7*; n = 4, *-Taf10*; n = 5 biological replicates. The bars correspond to the mean ± standard deviation. Kruskal-Wallis test followed by Dunn post hoc test if significant: ns; not significant, * <0.05; **<0.01; *** <0.001, **** <0.0001. **(I)** Cell density after passage at D2 and 1 day of extra culture. **(J)** Cell density evaluated by Crystal violet staining after passage at D4 and 2 days of culture. Scale bar; 150 µm. For D, F and G, the control conditions correspond to *RT7* cells treated with EtOH.

Recently, we have described the sequential TFIID assembly pathway that involves co-translational assembly steps ^13^. The early assembly events coincide with the co-translational formation of protein pairs, mostly through the dimerization of HFD-containing subunits. The products of the first assembly steps in turn access the second level of the assembly pathway by combining with each other in few, structurally well-defined steps. Importantly, assembly in step 2 occurs post-translationally and leads to the buildup of larger sub-complexes such as the core-TFIID, the 8-TAF complex and a partially assembled lobe A ^13^. The core-TFIID contains two copies of TAF5, TAF4/TAF12 and TAF6/TAF9, and is the basic structure of the both lobes A and B ^10–12^. The addition of the 3-TAF submodule composed of TAF2, TAF8 and TAF10 to the core TFIID, results in the formation of the 8-TAF complex which contains a full B lobe ^14^. In the third step, the products of step 2 engage co-translationally with the nascent TAF1 polypeptide. The pre-assembled building blocks of TFIID are recruited by the distinct assembly domains of nascent TAF1 as the distinct interaction domains of TAF1 emerge from the ribosome channel ^13^. The first N-terminal anchor point interacts with TBP, which engages with nascent TAF1 by binding to its TAND domain ^15^. The second interaction anchor point, the TAF6 binding motifs (T6BMs), interact with two copies of TAF6 HEAT domains of pre-assembled lobe A and lobe B, effectively bringing together these two lobes. The third anchor point at the end of the assembly process, recruits TAF7, which is known to interact with the TAF1 central domain (DUF3591/RAPiD) ^16,17^. Upon completion of TAF1 protein synthesis, a fully assembled TFIID is released and readily translocated in the nucleus^13^. An interaction between TAF7 and TAF11/TAF13 has also been identified in the cytoplasm ^18^ and it was proposed that the biochemically-reconstituted TAF1/TAF7/TAF11/TAF13/TBP sub-complex, also called S-TAF, could integrate the 8-TAF complex to form the holo-TFIID^19^.

Interestingly, TFIID can have variable compositions in higher eukaryotes (reviewed in ^20^). First, non-canonical TFIID complexes have been described, as TAF2- and TAF10-lacking TFIID complexes have been reported from yeast, *Drosophila* and human cells ^4,21–23^. Second, some TAFs have paralogs ^20^: *Drosophila* testis specific TAF4 (Nht), TAF5 (Can), TAF6 (Mia), TAF8 (Sa) and TAF12 (Rye) ^24,25^, vertebrates TAF4B ^26^, TAF7L ^27^ and TAF9B ^28,29^, and primate-specific TAF1L ^30^. In the mouse, TAF4B, TAF9B and TAF7L are associated with cell-specific functions. These paralogues in certain cases are co-expressed with their relative paralogs ^31–33^ and may even have partial overlapping functions ^31^. As TAF4B and TAF9B are part of TFIID ^34,35^ and TAF4 and TAF9 are present in 2 copies in TFIID ^11^, this suggests the possible combination between these paralogs within TFIID.

TFIID is important *in vivo* as mutations in *Taf7*, *Taf8* and *Taf10* lead to peri-implantation lethality in the mouse ^36–38^. Interestingly, in both *Taf8^-/-^* and *Taf10^-/-^* blastocysts, inner cell masses fail to outgrowth *in vitro,* while mutant trophectoderm is not affected ^36,37^, indicating that TAF8 or TAF10 requirements depend on the cellular context. These observations were further supported by conditional deletion of *Taf7* or *Taf10* in different cell types. While TAF7 is important for early thymocytes proliferation and differentiation, it is not required for the final step of thymocyte differentiation ^38^. Similarly, conditional deletion of *Taf10* in keratinocytes, liver cells and embryonic mesoderm has different effects depending on the developmental stage and on the cell types ^35,39,40^. Conditional deletion of *Taf10* in the early mesoderm leads to the absence of developing limb buds and to a growth arrest in the trunk region around embryonic day (E) 10 ^35^. Remarkably, cyclic transcription involved in the elongation of the embryo (reviewed in ^41^) is still active in the absence of TAF10 protein at E9.5, prior to the growth arrest, suggesting that active Pol II transcription does not rely on primarily on TAF10 in this context ^35^. Depletion of TAF10 severely affects the assembly of TFIID ^35,37,40^, whereas TAF7 depletion does not ^38^.

Overall, TFIID composition and requirements are variable depending on the cellular context, however as the available data have been gathered from different systems it is not possible to draw comparative conclusions about the function of partial and/or holo-TFIID complexes in cellular homeostasis. In this study, we analyzed the phenotype of the loss of *Taf7* and *Taf10* in comparable conditions, in pluripotent mouse embryonic stem cells (mESCs). Both subunits requirements depend on the cellular context and in the assembly chain of TFIID, TAF10 is an early subunit whereas TAF7 is a later one. We monitored the consequences of TAF7 or TAF10 depletion on holo-TFIID assembly and chromatin distribution, as well as on Pol II transcription, in order to test whether partial TFIID complexes are able to sustain Pol II transcription. Our data show that loss of *Taf7* has a less severe phenotype than the loss of *Taf10*. Importantly, the decrease in the amount of holo-TFIID is very drastic after efficient induction of *Taf7* or *Taf10* deletion. While TAF7 depletion results in the assembly of a TAF7-lacking TFIID, TAF10 depletion strongly affects TFIID assembly, leading to the only presence of the core-TFIID. Moreover, in both situations, TBP is still present on the promoters at the chromatin level. Surprisingly however, under either *Taf7* or *Taf10* deletion conditions, nascent Pol II transcription is still active. Conditional deletion of *Taf7* or *Taf10* in the developing mesoderm confirmed the difference in the phenotypic severities with *Taf7* conditional mutants being less affected than *Taf10* conditional mutants but later on the development of all conditional embryos is blocked. Altogether, our data demonstrate that partial TFIID modules are able to sustain nascent Pol II transcription in a given cellular environment, but do not support active gene expression changes over several cell divisions and/or development.

## Results

### Deletion of distinct TFIID subunits causes different phenotypic severities in mESCs

In order to get insight into the specific molecular consequences of the deletion of *Taf7* or *Taf10*, we derived mESCs from blastocysts carrying the inducible ubiquitously expressed *R26^CreERT^*^2^ allele ^42^ associated with the *Taf7^f^* or *Taf10^f^*(*R26^CreERT2/+^;Taf7^f/f^* and *R26^CreERT2/+^;Taf10^f/f^*, hereafter called *RT7* and *RT10*, respectively). The CRE recombinase induced deletion of the floxed alleles of *Taf7 ^f/f^* (*-Taf7)* or *Taf10 ^f/f^* (*-Taf10*) was induced by addition of 4-hydroxy tamoxifen (4-OHT) in the culture medium at day 0 (Fig 1B). Efficiency of the deletion was monitored at days 2 and 4 by western blot analyses and depletion of TAF7 protein in *RT7* mESCs was observed as early as day 2 and almost complete at day 4 (Figure 1C). Similarly, depletion of TAF10 protein in *RT10* mESCs was observed as early as day 2 and no longer detectable at day 4 (Figure 1C). Depletion of the protein of interest does not affect the expression of TAF4, TAF5, TAF6 and TAF12 TFIID subunits, and in agreement with our earlier studies, depletion of TAF10 results in the destabilization of its HF partner; TAF8 ^35,43^ (Figure 1C). The depletion of TAF7 and TAF10 proteins is homogenous within the cellular population as shown by immunolocalization (Figure 1D, Figure S1A). Altogether, these data indicate that we have established an efficient cellular model to study the effects of the depletion of TAF7 or TAF10 proteins.

Next, we investigated the effect of the 4-OHT treatment on cell proliferation and viability at day 2, 4 and 6. In culture, *RT7* and *RT10* mutant cells form smaller colonies compared to control cells (Figure 1E), indicating that both TAF7 and TAF10 are required for mESC maintenance. These data are in agreement with the incapacity of *Taf7^-/-^* null and *Taf10^-/-^* null inner cell masses to grow *in vitro* ^37,38^.

We then quantified the evolution of the number of cells in each mESCs clones, in control and mutant conditions. After two days of 4-OHT addition there were no detectable differences in cell numbers between the control and both mutant cells (Figure 1F). Depletion of TAF7 resulted in a minor decrease in cell number only at day 6. In contrast, TAF10 depletion resulted in a significant reduction in cell number already at day 4 (Figure 1F). To find out whether this effect on cell growth is also related to a decrease in cell proliferation, we performed EdU incorporation assay to quantify cells in S-phase. We observed a reduction tendency in the percentage of S-phase cells in both mutants from day 2 (Figure S1B). Thus, TAF7 and TAF10 do not appear to be required for cell cycle progression. The decrease in cell growth might to be related solely to viability of the cells. To test this hypothesis, we then investigated the evolution of the cell viability (Figure 1G). No differences between the control and the *RT7* and *RT10* mutant cells were observed at day 2 and day 4. However, at day 6, while depletion of TAF7 did not have any significant effect, depletion of TAF10 reduced seriously the viability (about a 2-fold magnitude) in *RT10* mutant cells. Analysis of apoptosis by quantification of Annexin V signal showed that the percentage of apoptotic cells ranged from 10 to 25% from day 2 to day 6, in control and mutant *RT7* cells (Figure 1H) indicating that TAF7 depletion does not induce apoptosis. On the contrary, although at days 2 and 4 no significant increase in apoptosis could be observed in mutant *RT10* cells, at day 6, the apoptosis rate increased 2-fold in the absence of TAF10 when compared to control conditions (Figure 1H). Finally, we evaluated the ability of the *RT7* and *RT10* mutant cells to form colonies after replating. When the mutant cells were split on day 2 and analyzed one day later (D2+1), no significant difference was observed in colony size or number of living cells for the two mutants (Figure 1I, Figure S1C). In contrary, when the mutant cells were split at day 4 and analyzed 2 days after (D4+2), their capacity to form colonies was severely impaired (Figure 1J, Figure S1D), indicating that both TAF7 and TAF10 are required for mESCs maintenance from day 4 onward.

Altogether, these data indicate that TAF10 in growing mESCs has a much more important role than TAF7. TAF10 is absolutely required for the growth and survival of the mESCs, whereas TAF7 depletion does not, or only weakly, affect mESC growth and survival. Importantly, this difference is not the consequence of the expression of TAF7 paralogs as they are not expressed in controls nor in the mutant cells (Figure S1E-G).

### The severity of the TAF10 depletion phenotype is not caused by defects in SAGA assembly

TAF10 is a shared subunit between TFIID and the SAGA coactivator complex, whereas TAF7 is a TFIID specific subunit ^44^. SAGA assembly is defective in E9.5 *Taf10* conditional mutant embryos ^35^. Therefore, we hypothesized that the difference in phenotype severity between mutant *RT7* and *RT10* mESCs could be associated with a combined impairment of both TFIID and SAGA function in *RT10* cells. To test this hypothesis, we focused on SUPT7L, the HFD-containing interaction partner of TAF10 in SAGA ^45^. Importantly, *Supt7l* loss of function in mESCs disrupts SAGA assembly ^46^. Western blot analysis of SUPT7L expression in the *RT7* line indicated that TAF7 depletion does not affect SUPT7L expression as expected (Figure 2A), but surprisingly, TAF10 depletion resulted in the loss of SUPT7L expression (Figure 2A) indicating that TAF10 is required for the stability of its two HFD partners; TAF8 and SUPT7L. These results indicate that *Supt7l* and *Taf10* loss have similar effects on SAGA assembly. In order to be able to deplete TAF7 in a context where SAGA assembly is impaired, we engineered a *Supt7l* homozygous mutation in the *RT7* line (hereafter called *RT7;Supt7l^-/-^*) using the same strategy as in ^46^, (Figure 2B). Co-depletion of SUPT7L and TAF7 in mutant *RT7;Supt7l^-/-^* cells, was validated by western blot at day 2, 4 and 6 (Figure 2C). We confirmed by immunoprecipitation coupled with mass spectrometry (IP-MS) using antibodies against SUPT20H, a subunit of the SAGA core structure, that SAGA is not assembled in absence of TAF10 and/or SUPT7L (Figure 2D) ^35,46^.

**Figure 2:**
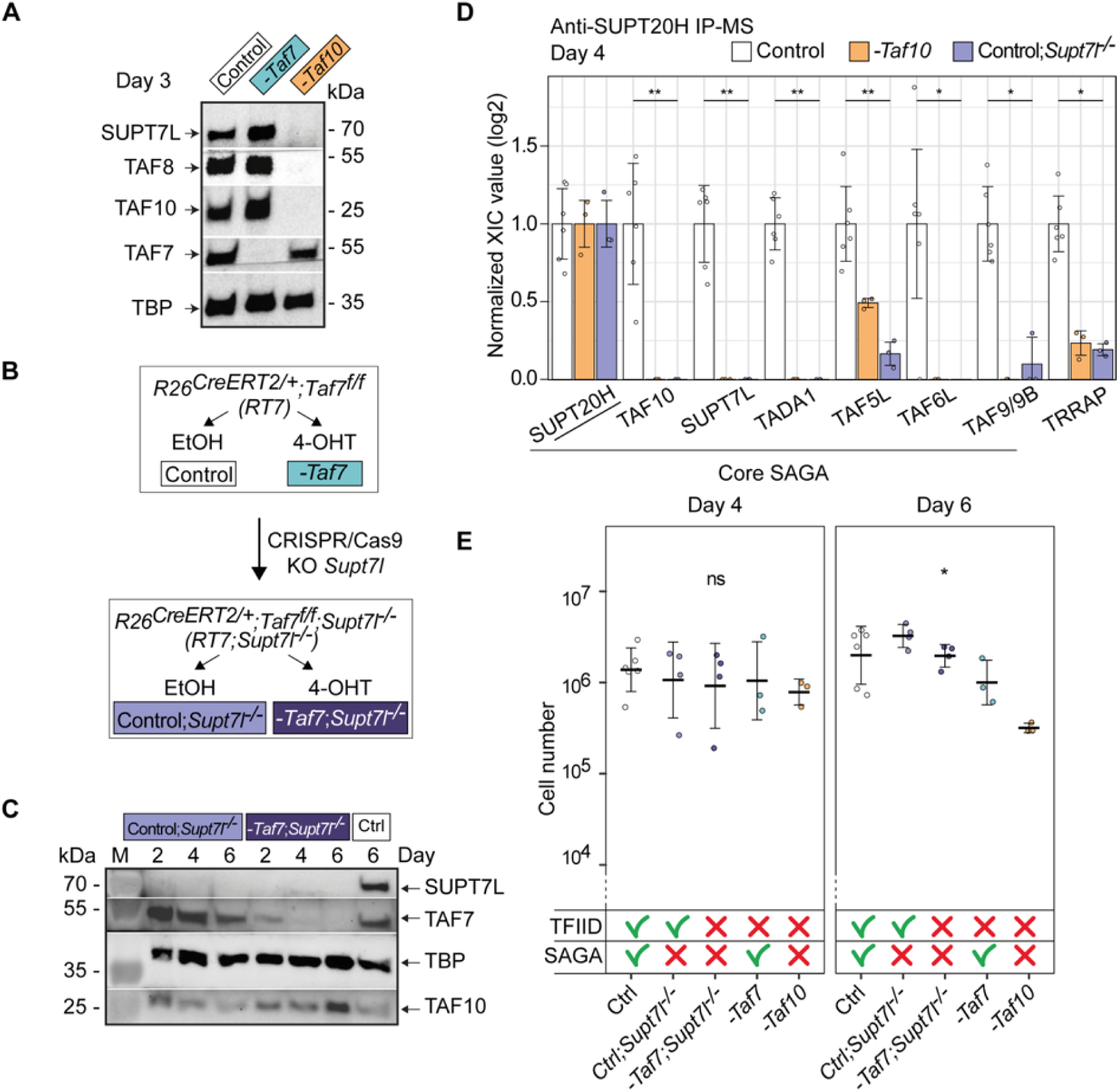
The more severe phenotype in TAF10 depleted mESCs is not due to the SAGA assembly defect. **(A)** Western blot analyses of SUPT7L, TAF8, TAF10, TAF7 and TBP expression after deletion of *Taf7* (-*Taf7*) or of *Taf10* (-*Taf10*). *RT7* and *RT10* cells were treated 3 days with 4-OHT and as control, *RT7* cells were treated 3 days with EtOH. **(B)** CRISPR/Cas9 strategy to generate the *R26^CreERT2/+^;Taf7^f/f^* ;*Supt7l^-/-^* (*RT7*;*Supt7l^-/-^*) mESCs. The second coding exon (exon 3) of *Supt7l* was deleted in *RT7* mESCs. Deletion of *Taf7* in the *Supt7l^-/-^* genetic background is induced by 4-OHT treatment. **(C)** Western blot analyses of SUPT7L, TAF7, TBP and TAF10 expression in *RT7;Supt7l^-/-^* cells after 2, 4 and 6 days of treatment with EtOH (Control; *Supt7l^-/-^*) or 4-OHT (*-Taf7*; *Supt7l^-/-^*). As a wild type control, *RT7* cells were treated 6 days with EtOH. M; molecular weight marker. **(D)** Anti-SUPT20H IP-MS analyses on nuclear enriched lysates from *RT10* and *RT7* cells treated 4 days with EtOH (Control, *RT7* and *RT10* data merged), *RT10* mESCs treated 4 days with 4-OHT (- *Taf10*) and *RT7*;*Supt7l^-/-^* cells treated 4 days with EtOH (Control;*Supt7l^-/-^*). For each of the proteins of interest, the XIC values of Control; *Supt7l^-/-^* and *-Taf10* lysates were normalized to those of *RT7* and *RT10* control cells treated with EtOH, respectively. Control; n = 2 biological replicates x 3 technical replicates, -*Taf10*; n = 1 x 3, Control;*Supt7l^-/-^*; n = 1 x 3. Means ± standard deviation are shown. Kruskal-Wallis test: * <0.05; **<0.01. **(E)** Total number of cells after 4 and 6 days of treatment. The control condition corresponds to the merging of *RT7* and *RT10* cells treated with EtOH. The impact on TFIID and SAGA assembly is indicated at the bottom. D4, D6: Ctrl; n = 6, Ctrl;*Supt7l^-/-^*; n = 4, *-Taf7;Supt7l^-/-^*; n = 4, *-Taf7;* n = 3, *- Taf10*; n = 3 biological replicates. Means ± standard deviation are shown. Kruskal-Wallis test followed by a Dunn post hoc test: ns; not significant, * <0.05.

*Supt7l* loss of function results only in a slight growth defect in similar culture conditions ^46^ and our different cellular measurements indicated that the growth and survival of non-treated *RT7;Supt7l^-/-^* mESCs are not affected (Figure 2E, Figure S2A, B). After induction of TAF7 depletion in the *Supt7l^-/-^* background, cell number, cell viability and cell death did not change significantly when compared to TAF7 depletion alone (Figure 2E, Figure S2A, B). Therefore, SAGA disruption does not aggravate the phenotype of TAF7 loss, strongly suggesting that the difference in the observed severity between mutant *RT7* and mutant *RT10* mESCs is not caused by the disruption of SAGA in mutant *RT10* cells, but by the different molecular consequences associated to the loss of TAF7, or TAF10 in TFIID.

### Depletion of TAF7 or TAF10 leads to the formation of distinct partial TFIID complexes

In order to analyze the molecular changes occurring in the TFIID complex following depletion of TAF7 or TAF10, we first performed gel filtration coupled to western blot analyses on *RT7* or *RT10* mESCs nuclear enriched whole cell lysates (NWCEs) collected at day 3, in control and mutant conditions. In control cells (*RT10* treated with vehicle), fractions B-C contained the TFIID complex, as suggested by the molecular weight and the co-localization of TBP and all the tested TAFs (Figure 3A, blue box). In mutant *RT7* extracts, the profile is very similar to the control lysate, except for the absence of TAF7, strongly suggesting that a TFIID complex without TAF7 is present in these cells (Figure 3B, green box). In mutant *RT10* cells however, very little high molecular weight TAF complex could be detected, Consequently, TAF4, TAF5 and TAF6 were relocated to lower molecular weight complexes, peaking at K-L fractions (Figure 3C, purple box), indicating that the integrity of the holo-TFIID complex was lost in these cells after TAF10 depletion.

**Figure 3:**
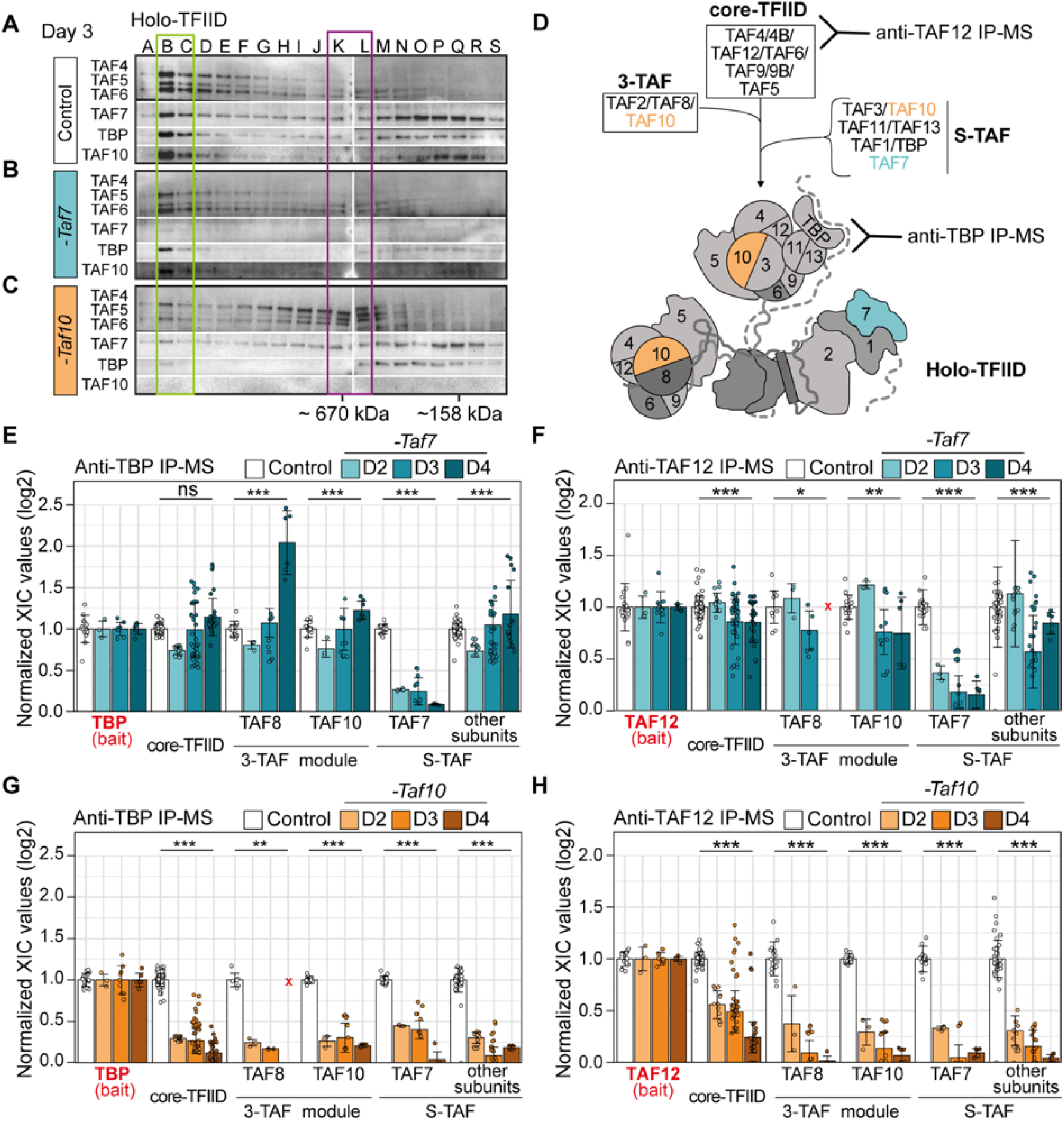
Depletion of TAF7 or TAF10 differentially affects TFIID assembly. **(A-C)** Gel filtration coupled to western blot analysis of TAF4, TAF5, TAF6, TAF7, TAF10 and TBP expression in control *RT10* mESCs treated with EtOH (Control) (A), *RT7* (*-Taf7*) and *RT10* (*- Taf10*) depleted mESCs treated with 4-OHT (B,C) for 3 days. The letters on the top correspond of the different fractions analyzed. Molecular weight positions are indicated at the bottom. Positions of the different complexes are indicated by boxes (n=1). **(D)** Assembly of the holo-TFIID complex representation. Anti-TBP antibodies will detect the presence of the holo-TFIID only while anti-TAF12 antibodies will detect all intermediates from core-TFIID to holo-TFIID. **(E-H)** Relative normalized XIC values of TFIID subunits from anti-TBP IP- MS (E,F) and anti-TAF12 IP-MS (G,H) from *RT7* (E,G) and *RT10* mESCs (F,H) at day 2, 3 and 4 of EtOH (Control) or 4-OHT (-*Taf7* and -*Taf10*) treatment. Subunits of the same submodules (see D) were merged together except for the bait proteins TBP (E,F) and TAF12 (G,H), and TAF7, TAF8 and TAF10. Note that TAF2 data were not taken into account because TAF2 was poorly detected in the controls in the 2 IP-MS. Core-TFIID corresponds to TAF4, TAF5, TAF6, TAF9/9B and TAF12, S-TAF corresponds to TAF1, TAF3, TAF11, TAF13 and TBP for. D2; n = 1 biological replicate x 3 technical replicates, D3; n = 3 biological replicates x 3 technical replicates, D4; n = 2 biological replicates x 3 technical replicates. Each dot corresponds to one measure of one subunit. Red crosses indicate proteins not detected in the control condition. Means ± standard deviation are shown, Kruskal-Wallis test: ns; not significant, * <0.05; **<0.01; *** <0.001.

To explore the composition of the assembled partial complexes after TAF7, or TAF10 depletion, we performed IP-MS using anti-TAF12 and anti-TBP antibodies on NWCEs after 2, 3 and 4 days of treatment. As TAF12 is part of the core-TFIID and TBP only part of the holo-TFIID complex, these 2 types of IPs allow to discriminate between the different partial or complete TFIID complexes (Figure 3D). Anti-TAF12 IP detects all the intermediate partial TFIID complexes assembling on the core-TFIID whereas anti-TBP IP only immunoprecipitates the holo-TFIID complex (Figure 3D). As TBP is also part of the SL1 and TFIIIB complexes, which are involved in Pol I and Pol III transcription initiation, respectively ^47^, we tested whether the lack of TAF7 or TAF10 would interfere with the interaction between TBP and SL1 subunits, TAF1A, TAF1B, TAF1C and TAF1D, or the BRF1 subunit of TFIIIB (Sup Figure 3A, B). Remarkably, as already observed in the *TCre;Taf10* mutant embryos ^35^, TAF10 depletion resulted in increased interaction between TBP and its non-TFIID partners, and mainly with the RNA pol III machinery (Figure S3B). These data indicate that TBP is relocated to these complexes when TAF10 is depleted, suggesting that TBP incorporation in TFIID is indeed impaired when TAF10 is absent in the cells. This relocation was not observed when TAF7 was depleted (Sup Figure 3A) suggesting that TBP incorporation into TFIID is not affected under these conditions.

The depletion of TAF7 induced the loss of TAF7 in the anti-TBP or -TAF12 IP-ed complexes, as expected, but did not induce significant changes in the IP-ed subunits of TFIID (Figure 3E, F, Sup Figure 3C, D). In both IP-MS, the signal detected for each of the TFIID subunits remained relatively stable on each day of the treatments analyzed. This indicates that the absence of TAF7 has a very limited effect on TFIID assembly, resulting in a TAF7-lacking TFIID complex, as suggested also by the gel filtration analysis (Fig 3A, B). In contrast, depletion of TAF10 caused a more severe impairment of TFIID assembly as shown by the important changes in the TFIID subunits detected in either anti-TBP (Figure 3G, Sup Figure 3E) or anti-TAF12 IPs (Figure 3H, Sup Figure 3F). First, there is a strong decrease in the interaction between TBP and all the other TFIID subunits (about 4-fold already at day 2) in mutant *RT10* extracts (Figure 3G, Sup Figure 3E). Second, anti-TAF12 IP-MS showed that when TAF10 is depleted, TAF12 maintains a stronger interaction with TAF5, TAF6, TAF9/9B and TAF4/4B core-TFIID subunits, but interactions with the other TAFs or TBP are severely decreased. In particular, the signals of the 3-TAF complex and S-TAF complex subunits decreased by about 4-fold at day 2 in the *RT10* mutant extracts compared to the control condition (Figure 3H). Signal of TAF5, TAF6, and TAF9/9B only decreased by 2-fold (Sup Figure 3F) and gradually decreases on the following days while TAF4/4B – TAF12 interaction was maintained up to day 3 (Sup Figure 3F). Altogether, these data indicate that the core-TFIID is the main remaining partial TFIID complex present in the mutant *RT10* mESCs, and confirm that TAF10 presence is important for the recruitment of other subunits on the core-TFIID to assemble the holo-TFIID.

Altogether, these data demonstrate that TFIID assembly is differentially affected by either the depletion in TAF7 or by that of TAF10. A TFIID-like complex lacking TAF7, hereafter called TAF7-less TFIID, persists in mutant *RT7* cells. However, in mutant *RT10* cells, holo-TFIID complex is not assembled, and a partial core-TFIID-like complex containing TAF4/4B, TAF5, TAF6; TAF9/9B, TAF12, but lacking TBP, remains.

### TBP is recruited to the chromatin in TAF7 and TAF10 depleted mESCs

As at day 3 and 4, holo-TFIID depletion is well established while there is no major phenotype in both TAF7 or TAF10 depleted mESCs at day3, we decided to focus our next analyses at that stage to avoid potential secondary effects. Moreover, as depletion of TAF7 or TAF10 result in distinct partially assembled TFIID complexes, either containing or lacking TBP respectively, we next investigated whether TBP would be recruited on promoters under these conditions. To this end, first we prepared and analyzed WCE, cytoplasmic-enriched (CE), nuclear-enriched (NE) and chromatin (ChrE) extracts from control mESCs at day 3 ^48^. As expected, western blot analysis indicated enrichment of H3 in chromatin extracts and of Tubulin in the cytoplasmic fraction in control WCE (Sup Figure 4A). TAF4, TAF5, TAF6 and TBP are more intense in the control ChrE, indicating a tight association of these TFIID subunits with the chromatin. After day 3 of 4-OHT treatment, while TAF7 is strongly depleted in the WCE, CE, NE and ChrE fractions, there is no obvious difference in the levels of TAF4, TAF5, TAF6, TAF8 and TAF10 between control and 4-OHT treated RT7 cells (Sup Figure 4B), further reinforcing our previous observation that a TAF7-less TFIID partial complex is present in the TAF7 depleted cells. Interestingly TBP levels appear not or only mildly decreased in the TAF7 depleted condition, suggesting that the loading of TBP on the chromatin might be not or only slightly affected under the TAF7-lacking condition. Remarkably, when TAF10 is depleted, as expected no more TAF8 is detected ^35,43^ in all the fractions, however no obvious differences could be observed in TAF4, TAF5, TAF6 and TAF7, as well as TBP levels (Sup Figure 4C), suggesting that TBP recruitment at the promoters may not be significantly affected in TAF10 depleted cells.

### TAF7-less TFIID, or partial core TAF complexes are recruited at the promoters in TAF7 or TAF10 depleted mESCs, respectively

In order to get more insight into the distribution of TBP on promoters in control, TAF7, or TAF10 depleted mESCs, we conducted CUT&RUN analyses at 3 days after 4-OHT treatment, or no treatment, using an anti-TBP antibody. Annotation analyses indicated that in all conditions TBP peaks were primarily located in promoter regions, as expected (Sup Figure 5A-D). Nevertheless, comparison of the accumulation of TBP at the positions of the TBP bound regions in the control conditions, showed that TBP is still present on most of the promoters in TAF7- (Figure 4A, Sup Figure 5E) or in TAF10-depleted cells (Figure 4F, Sup Figure 5J). Nevertheless, the mean profile of TBP indicated that in both conditions, there is a small reduction in the level of TBP present at the TBP bound regions (Figure 4B, G).

**Figure 4:**
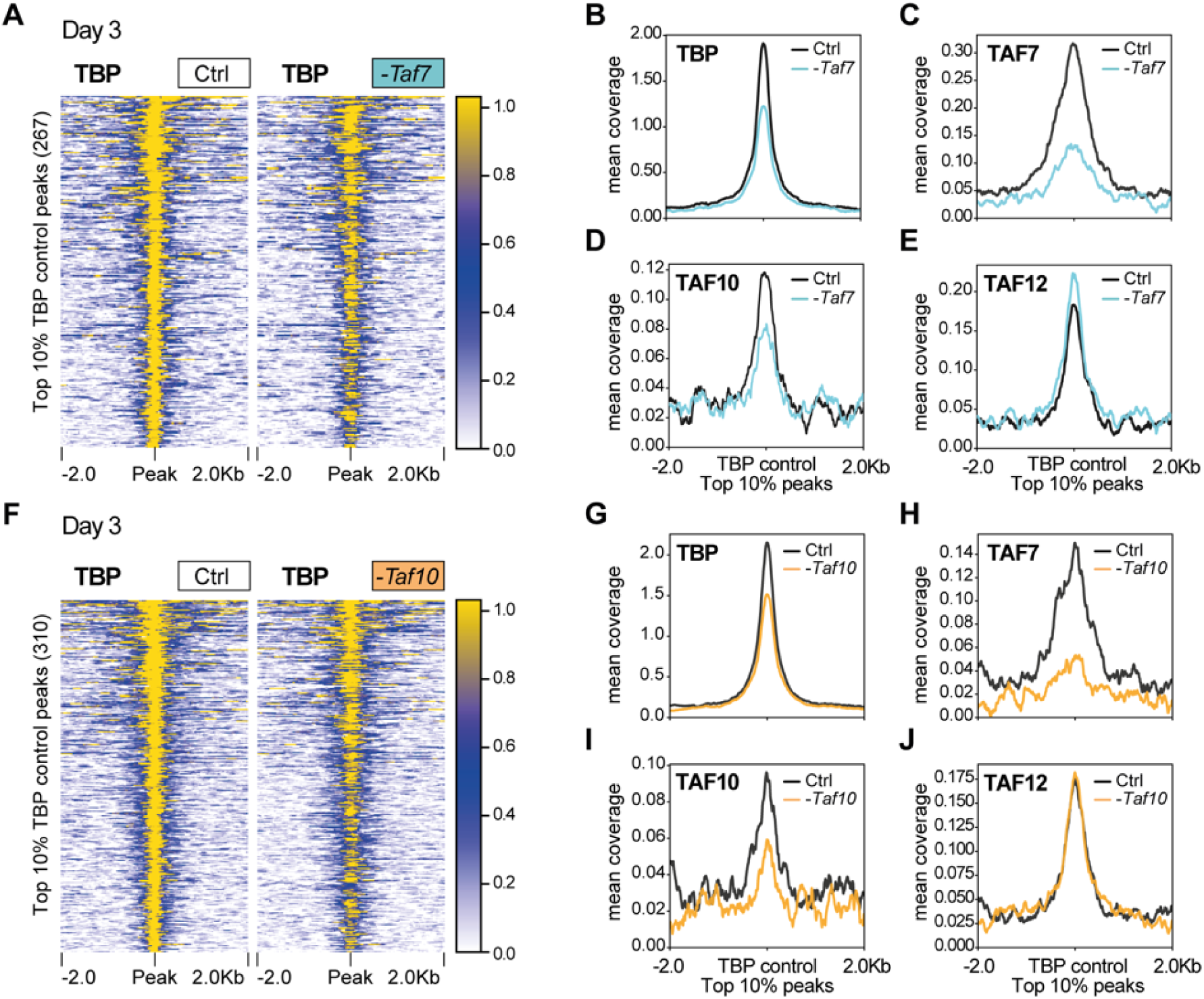
TBP is still recruited at promoters in *TAF7* or *TAF10*-depleted mESCs. **(A)** Heatmap showing the TBP distribution at the position of the top 10% (fold enrichment) TBP CUT&RUN peaks (± 2 kb) detected in the control condition, in *RT7* control (Ctrl) and TAF7-depleted (-*Taf7*) mESCs after 3 days of treatment. **(B-E)** Mean profile of TBP (B), TAF7 (C), TAF10 (D) and TAF12 (E) protein accumulation at the location of the TBP control peaks, in control (black line) and in TAF7 depleted (blue line) mESCs. **(F)** Heatmap showing the TBP distribution at the position of the top 10% TBP CUT&RUN peaks (± 2 kb) detected in the control condition, in *RT10* control (Ctrl) and TAF10-depleted (-*Taf10*) mESCs after 3 days of treatment. **(G-J)** Mean profile of TBP (G), TAF7 (H), TAF10 (I) and TAF12 (J) protein accumulation at the location of the TBP control peaks, in control (black line) and in TAF10 depleted (orange line) mESCs.

We also carried out anti-TAF7, -TAF10 and -TAF12 CUT&RUN experiments to measure the impact of TAF7 or TAF10 depletion on the accumulation of TAF7, TAF10 and TAF12 at the positions of TBP bound promoters detected above. As expected, we observed a strong depletion of TAF7 in 4-OHT treated *RT7* mESCs (Figure 4C, Sup Figure 5G), but no important decrease of either TAF10 or TAF12 was detected at these sites (Figure 4D, E, Sup Figure 5H, I), further indicating that under TAF7 depletion a TAF7-less TFIID is still binding to these promoters. Moreover, we observed a decrease of TAF10 in 4-OHT treated *RT10* mESCs at the TBP bound sites together with a strong decrease of TAF7 at these promoters (Figure 4I, H; Sup Figure 5M, L). Importantly, TAF12 accumulation was not affected in these TAF10-lacking cells (Figure 4J, Sup Figure 5N), suggesting that a core-TFIID-like complex containing TAF12, may bind to these sites possibly together with TBP.

### Global Pol II activity is maintained in TAF7 and TAF10 depleted mESCs

According to the text book view, holo-TFIID is the first GTF to recognize the promoters and to start the formation of the PIC. After depletion of TAF7 or TAF10, the amount of holo-TFIID drastically decreased in mESCs. We therefore wondered about the impact of the presence of the remaining partial TAF-containing complexes on nascent Pol II transcription. We first analyzed the incorporation of 5-ethynyl uridine (EU) after one hour of incubation in the culture medium. EU incorporation was assessed after 2 to 5 days of 4-OHT treatment (Figure 5A) and no obvious differences were observed between control and mutant *RT7,* or *RT10* cells (Figure 5A, B). To control the metabolic RNA labeling efficiency, we pre-incubated control cells with two different Pol II transcription inhibitors triptolide (transcription initiation inhibitor) or flavopiridol (transcription elongation inhibitor). As expected, EU incorporation was readily detected in non-treated cells, and abolished in the triptolide or flavopiridol treated cells (Figure 5C, D). To further verify that RNA Pol II is actively transcribing in our mutant cell lines, we investigated the phosphorylation status of the C-terminal domain of the Pol II subunit RPB1 in mutant *RT7* and *RT10* cells. Western blot analysis of RPB1 from extracts obtained at day 3 indicated that the hypo (Pol II-A) and hyper (Pol II-O) phosphorylated RPB1 levels were comparable in both mutant and control conditions (Figure 5E). We then specifically analyzed the presence of the phosphorylation of the serine 5 (RPB1^pSer5^, pSer5) and serine 2 (RPB1^pSer2^, pSer2) at day 3 and day 4 by immunofluorescence (IF). RPB1^pSer5^ and RPB1^pSer2^ are associated with the initiation and the elongation status of Pol II, respectively (reviewed in ^49^). Similarly, no obvious difference in RPB1^pSer2^ and in RPB1^pSer5^ phosphorylation signal could be observed between control and mutant *RT7* and *RT10* cells (Sup Figure 6A, B) indicating that Pol II transcription initiation and elongation is active and is not obviously affected after TAF7 and TAF10 depletion.

**Figure 5:**
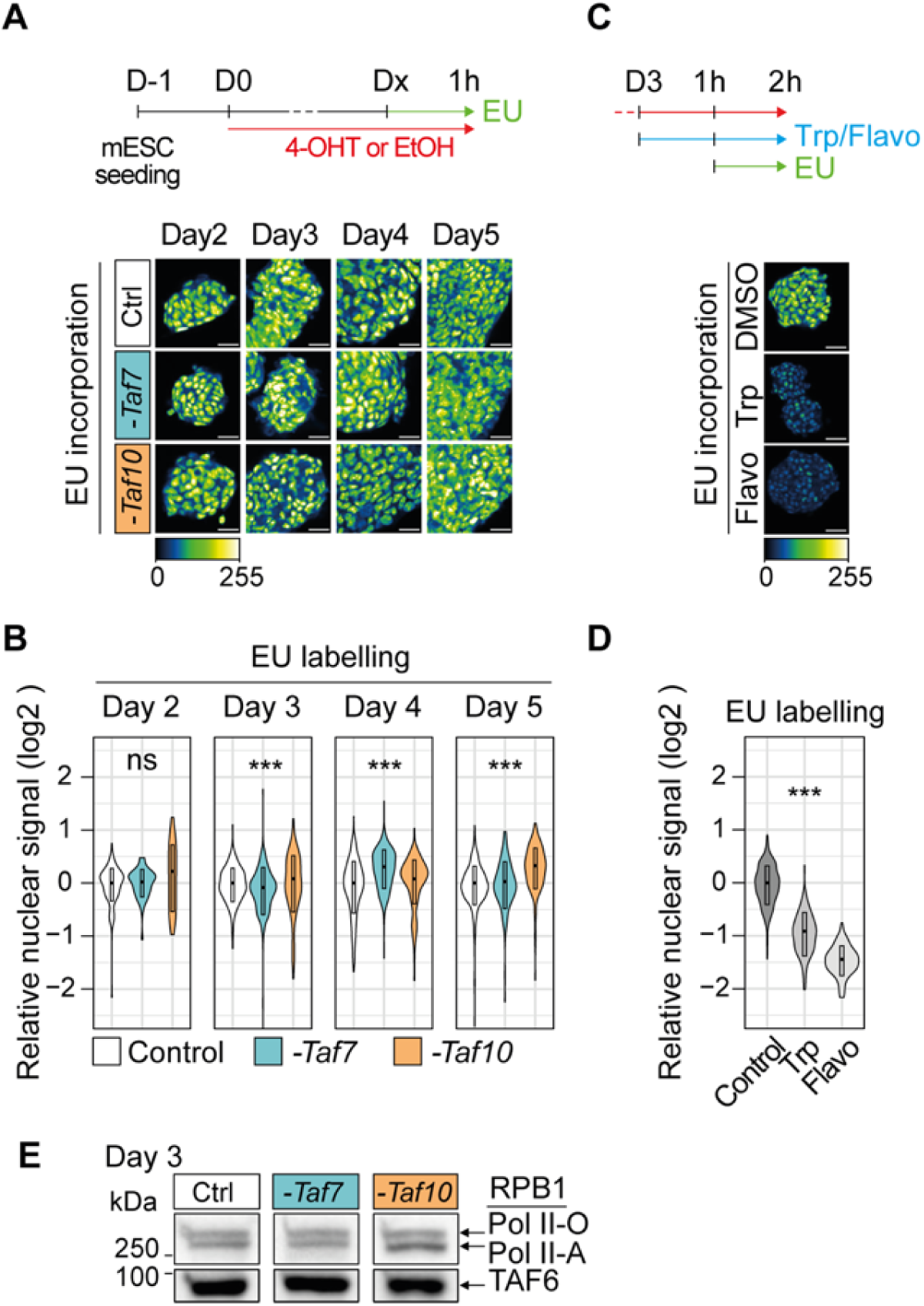
Depletion of holo-TFIID in absence of TAF7 or TAF10 has only limited defects on Pol II global transcription. **(A,B)** Analysis of nascent transcription by one hour incubation with 1 mM of 5-ethynyluridine (EU) after 2 to 5 days of 4-OHT (-*Taf7* or -*Taf10*) or EtOH (Control) treatment labeling in *RT7* and *RT10* cells after 2 to 5 days of 4-OHT (- *Taf7* or -*Taf10*) or EtOH (Control) treatment. The time course of the experiments is indicated at the top. Representative pictures and quantification presented as violin plots are shown in (A) and (B), respectively. D2; two biological replicates (n=2), D3; n=3, D4; n=3, D5; n=2. **(C,D)** EU incorporation in control cells after 2 hours of treatment with Pol II transcription inhibitors (500 µm of triptolide (Trp) or 300 µM of flavopiridol (Flavo)). Representative pictures and quantification presented as violin plots are displayed in (C) and (D), respectively (n=1). Hundreds of measurements for individual nuclei were performed for the quantification, scale bar: 50 µm. Kruskal Wallis test: ns; not significant, *<0.05; **<0.01; ***<0.001. **(E)** Western blot analysis of the phosphorylated forms of RBP1 using an anti-RBP1 antibody from chromatin extracts of EtOH-treated *RT7* mESCs (Ctrl) and 4-OHT treated *RT7* (*-Taf7*) and *RT10* (*-Taf10*) mESCs after 3 days of treatment. The hyper-phosphorylated RPB1 (Pol II-O, active) and hypo-phosphorylated (Pol II-A) are indicated on the right. At the bottom anti-TAF6 western blot analysis as loading control.

Altogether, these data suggest that despite a dramatic decrease in the amount of holo-TFIID, the global activity of the Pol II is maintained in TAF7 or TAF10 depleted mESCs.

### TAF7 or TAF10 depletion have a very weak impact on nascent Pol II transcription at day 3

As global Pol II transcription is not obviously impaired in the TAF7 and TAF10 depleted mESCs, we investigated whether the depletion in TAF7 or TAF10 would have a specific impact on Pol II transcription mESCs by analysis of nascent RNA by 4-sU-seq (Figure 6A). To limit any bias associated with the indirect effects of the drastic reduction of holo-TFIID, we carried out our analyses at day 3. As spikes, we added unlabeled *S. pombe* (yeast) as well as *Drosophila* S2 4-sU labelled RNA. Only reads from 4-sU labeling experiments could be mapped on the *Drosophila* and mouse genomes while no reads from purified transcripts could be mapped on the yeast genome, confirming the enrichment of newly synthetized RNAs genome wide (Figure 6B).

**Figure 6:**
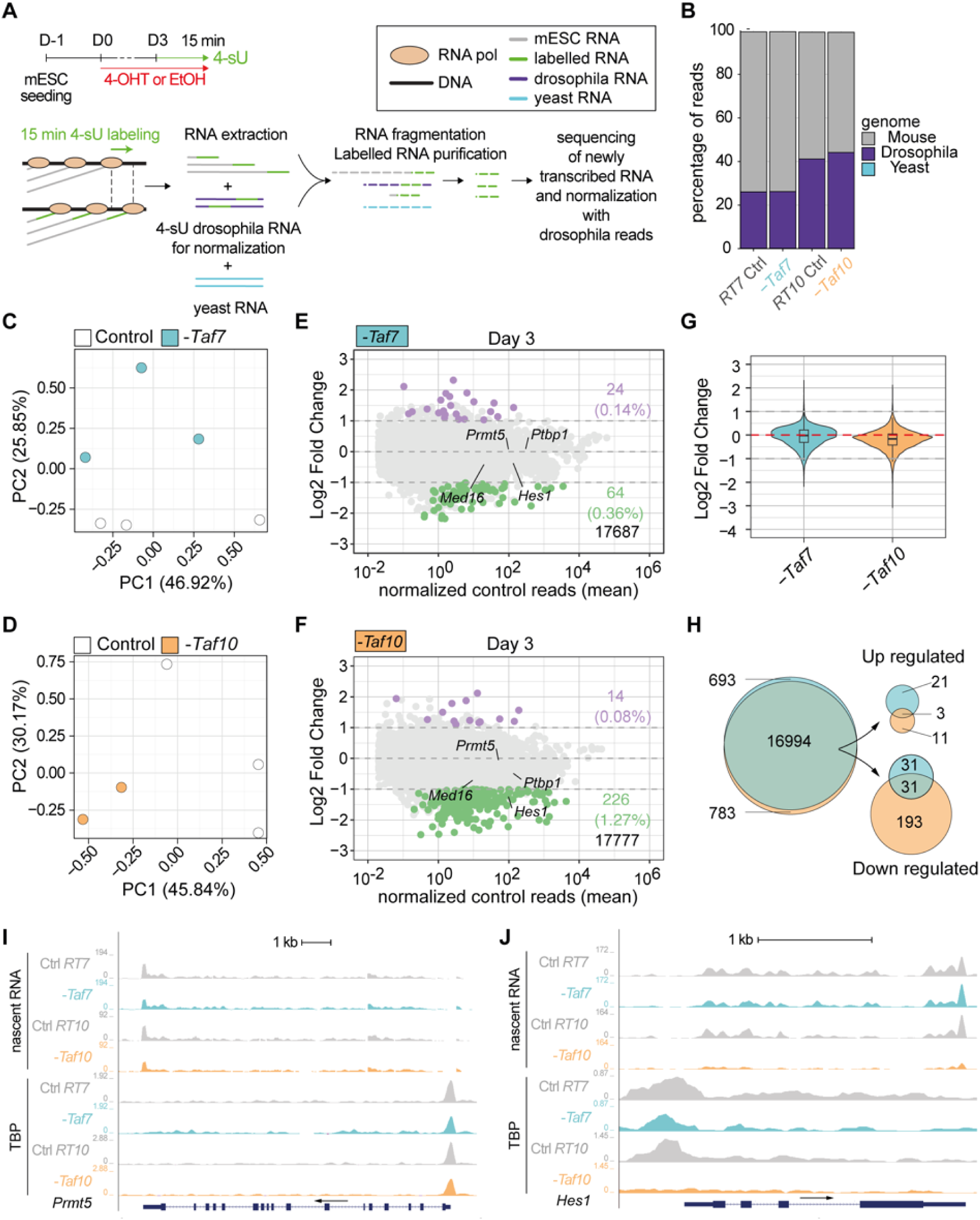
TAF7 loss has a milder impact on RNA pol II gene transcription than TAF10 loss after 3 days of treatment. **(A)** Time course and scheme of the nascent RNA sequencing. After three days of treatment with 4-OHT (*-Taf7* and *-Taf10* for mutant *RT7* and *RT10* mESC, respectively) or EtOH (Control), nascent transcripts were labeled with 4-sU during 15 minutes. 4-sU labelled RNA from *Drosophila* S2 cells were used as spike for normalization and unlabelled yeast RNA as a control for the purification of the nascent transcripts. **(B)** Percentage of mouse (grey), *Drosophila* (purple) and yeast (cyan) reads in the different datasets. **(C-D)** Principal component analysis (PCA) of the nascent transcriptomes of *RT7* (C) and *RT10* (D) control (white dot) and mutant (colored dot) mESCs. **(E,F)** MA plots of 4-sU-seq from D3 mutant versus control RT7 (E) and RT10 (F) with the number and percentage of significantly down-(green) or up-regulated (purple) protein coding transcripts displayed on the right and total transcripts detected at the bottom. A threshold of 20 normalized reads per gene length in kb was used to select actively expressed protein coding genes. Significance transcripts were filtered using an adjusted P value ≤ 0.05 and an absolute Log2 Fold Change ≥ 1. **(G)** Comparisons of global Log2 fold changes in TAF7- and TAF10-depleted mESCs. **(H)** Venn diagrams analysis of all (left), up-regulated (top, right) and down-regulated (bottom, right) protein coding transcripts in TAF7- and TAF10-depleted mESCs. **(I,J)** UCSC genome browser views of nascent RNA (top) and TBP distribution (bottom) at *Prmt5* (I) and *Hes1* (J) loci between control (Ctrl *RT7* and Ctrl *RT10*), TAF7 depleted (-*Taf7*) and TAF10 depleted (- *Taf1*0) mESCs. Y axes indicate the genomic coverage. Arrows indicate direction of transcription.

PCA analyses indicated that in both mESCs lines, depleted and control condition samples are well separated (Figure 6C, D). Interestingly, while in *RT10* cells, the treatment is associated with PC1 (Figure 6D), it is associated with PC2 in *RT7* cells (Figure 6C), suggesting that the nascent transcription is more altered after TAF10 depletion than TAF7 depletion.

Differential expression analyses confirmed that TAF7 and TAF10 depletions are not accompanied by a global reduction in Pol II transcription, as only a small fraction of the nascent transcripts is affected (Figure 6E, F). In *RT7* mESCs, 0.36% of transcripts (64) were significantly downregulated versus 0.14% (24) upregulated (Figure 6E), while in *RT10* mESCs, 1.26% of transcripts (226) were significantly downregulated versus 0.08% (14) upregulated (Figure 6F) in the 4-OHT treated compared to the control conditions, with a threshold of -1 Log2 fold change and an adjusted p-value of 0.05. While there is no obvious difference in the ratio of down-versus up-regulated transcripts in TAF7-depleted mESCs, there are much more down-regulated transcripts in TAF10-depleted cells (Figure 6E, F) suggesting that there might be a very slight decrease in global transcription in *RT10* mutants compared to *RT7* mutants (Figure 6G). Venn diagram analysis showed partial overlap of commonly differentially regulated transcripts (Figure 6H), but overall, no relevant GO categories were significantly enriched due to the limited number of differentially expressed transcripts.

Intersection of our CUT&RUN and 4-sU-seq data analyses suggested that when nascent transcription is not significantly affected by TAF7 or TAF10 depletion, the TBP peaks were not affected as seen at different loci on UCSC genome browser snapshots (see Figure 6I, Sup Figure 6C, D). In contrast, in the case of the *Hes1* gene which is significantly down-regulated in TAF10 depleted mESCs (Figure 6F), TBP was evicted from the *Hes1* promoter (Figure 6J) indicating that genes which are downregulated following TAF10 depletion seem to have lost TBP.

In conclusion, our data indicate that Pol II global activity is not seriously affected in conditions where holo-TFIID is depleted in either TAF7- or TAF10-lacking mESCs. In line with the above biochemical analyses of TFIID and TFIID-like complexes, TAF10 depletion has a somewhat stronger effect than TAF7 depletion for the transcription of a subset of genes. This suggests that the TAF7-less TFIID complex might be able to allow efficient recruitment of Pol II on the promoters and that the partial TAF assemblies detected in the TAF10 depleted cells, while being also globally sufficient to sustain Pol II transcription, fail to do so for a subset of genes.

### Embryonic deletion of either Taf7 or Taf10 induces different embryonic phenotypes with distinct severities

Our experiments performed on mESCs in steady state conditions indicate that Pol II transcription is not majorly affected at day 3 (Figure 5, 6). However, at day 6, it is obvious that the mESCs are affected, especially when TAF10 is depleted (Figure 1) suggesting that while no major defects are detected initially, Pol II transcription is actually affected on a longer term. To analyze the effect of the TAF7 and TAF10 depletion in a more dynamic *in vivo* transcriptional context, we compared the conditional deletion of *Taf10* or *Taf7* in the early mesoderm using the *T-Cre* transgenic line which is active as early as embryonic day (E) 6.5 ^50^. We had previously shown that efficient depletion of TAF10 in the early mesoderm results in a growth arrest, while cyclic transcription in the presomitic mesoderm associated with the periodic production of somites is not affected ^35^. Efficiency of TAF7 depletion was assessed by immunofluorescence (Figure S7A, B). As expected from the recombination pattern of the *T-Cre* line, the structures above the heart are not targeted and therefore normal^50^. Tg(T-Cre/+);*Taf7^f/f^* or Tg(T-Cre/+);*Taf10^f/f^* mutant embryos (hereafter called *TCre;Taf7* and *TCre;Taf10*, respectively) are similar at E 9.5. The only visible difference that we could detect was that while no limb buds are present in *TCre;Taf10* mutant embryos ^35^, *TCre;Taf7* embryos displayed forelimb buds although smaller when compared to the control (Figure 7B, C compared to 7A, white arrowhead). At E10.5, both *TCre;Taf10* and *TCre;Taf7* mutant embryos display a growth arrest in the trunk and tail regions (Figure 7E, F compared to D). After E10.5, in addition to the presence of limb buds in the *TCre;Taf7* embryos, more differences could be observed between the two mutants. First, enlarged pericardia are observed in all the *TCre;Taf7* mutant embryos (Figure 7E, white arrow) suggesting that *Taf7* deletion leads to cardiovascular defects not observed in *TCre;Taf10* mutant or control embryos (Figure 7D, F, white arrow). Second, while *TCre;Taf10* mutants could not be retrieved after E11.5 due to placenta and allantois degeneration ^35^, *TCre;Taf7* mutant embryos could be still detected at E12.5 characterized by hypotrophied limb buds, trunk and tail regions (Figure 7H). Third, presence of blood cells was obvious in the vasculature of controls and *TCre;Taf7* mutant embryos (Figure 7J, K, black arrowhead), but not of *TCre;Taf10* mutant embryos (Figure 7L, black arrow). Altogether, these data indicate that conditional deletion of *Taf7* or *Taf10* in the same genetic context leads to a similar growth arrest phenotype, but different outcomes. *Taf10* conditional deletion leads to a more severe phenotype associated with lack of forelimbs and red blood cells, while *Taf7* conditional deletion leads to a milder phenotype and is associated with specific cardiovascular defects. These embryonic data confirm the difference in phenotype severity observed in TAF7-versus TAF10-depleted mESCs and further support that TAF10-depletion has a stronger impact on TFIID assembly compared to TAF7-depletion, as shown by our IP-MS analyses.

**Figure 7:**
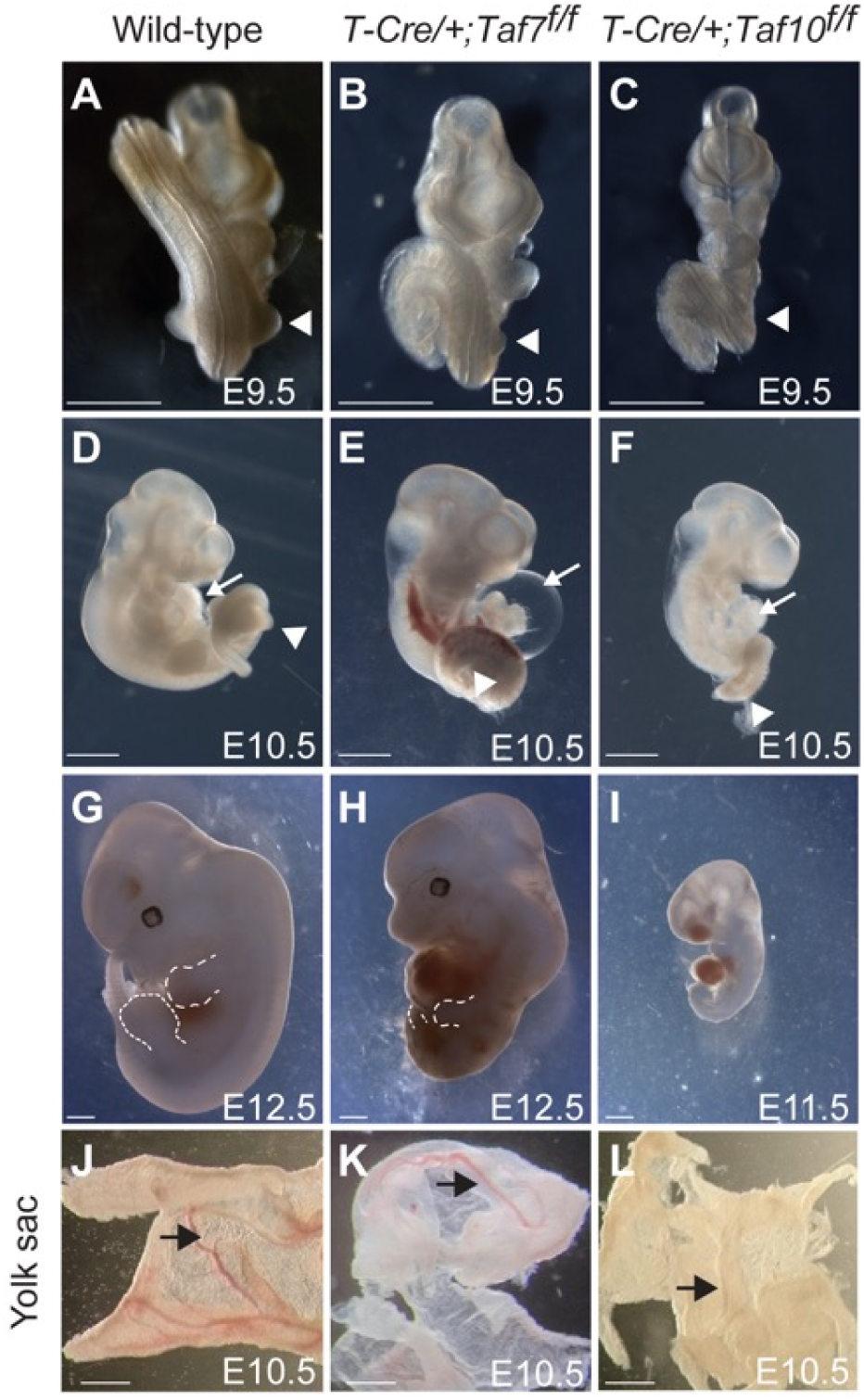
Conditional deletion of *Taf7* or *Taf10* in the early mesoderm results in similar yet different phenotypes. **(A-I)** Wholemount view of wild-type (A, D, G), Tg(T- Cre/+)*;Taf7^f/f^* mutant (B, E, H) and Tg(T-Cre/+)*;Taf10^f/f^* mutant (C, F, I) embryos at E9.5 (A- C, n>4), E10.5 (E-F, n>4), and E12.5 (G, H, n>2). As no Tg(T-Cre/+)*;Taf10^f/f^* mutant embryos could be recovered at E12.5, a E11.5 Tg(T-Cre/+)*;Taf10^f/f^*mutant embryo is shown (I). White arrowheads (A-C) indicate the position of the forelimbs. Dashed lines (G, H) indicate the limb buds. White arrows (D-F) indicate the heart. **(J-L)** Wholemount views of wild-type (J), Tg(T-Cre/+)*;Taf7^f/f^* (K) and Tg(T-Cre/+)*;Taf10^f/f^* (L) yolk sacs embryos are showed (n>3). Black arrows indicate the presence of blood vessels. Scale bars; 1 mm.

In conclusion, our *in vivo* results demonstrate that embryonic deletion of either *Taf7* or *Taf10* induce different embryonic phenotypes with distinct severities. Thus, while our *in vitro* data demonstrated that partial TFIID modules are able to sustain nascent Pol II transcription in mESCs, these partial TFIID assemblies cause embryonic lethality over time as they do not support all active gene regulatory changes during development.

## Discussion

In this study we have analyzed the depletion of two different TFIID subunits; TAF7 and TAF10 in the same cellular context. Using a CRE/*Lox*P conditional approach in mESCs, we showed that depletion of TAF10 leads to a more severe phenotype than TAF7 depletion. The increased severity of the TAF10 depletion phenotype is a direct consequence of a TFIID defect, but not a consequence of a SAGA defect, suggesting that TAF7 and TAF10 depletion have a different impact on TFIID. Analysis of TFIID assembly indicated that TAF10 depletion has a more pronounced effect on TFIID assembly than TAF7 depletion. In the absence of TAF7, an almost fully assembled TFIID complex lacking only TAF7 was detected while in the absence of TAF10, TFIID assembly was altered at the early stages and the core-TFIID, the basic building block of holo-TFIID, was the major TFIID submodule detected. Analyses of the effect of the depletion of TAF7 or TAF10 on the distribution of TBP on the chromatin showed only a modest reduction of TBP recruitment. Interestingly, analysis of the presence of TAF7, TAF10 and TAF12 at these TBP peaks in TAF7 depleted cells confirmed the depletion of TAF7, with only a minor effect on TAF10 and TAF12, in good agreement with the recruitment of a TAF7-less TFIID complex at the active promoters in these cells. Analysis of the presence of the same subunits at the TBP peaks in the TAF10 depleted mESCs confirmed the depletion of TAF10. However, while TAF7 occupancy levels were down regulated, TAF12 levels appeared not impacted, confirming that the major TFIID partial complex assembled in these cells is the core TFIID and also suggesting that this partial submodule can be recruited at the promoters when holo-TFIID is depleted. Surprisingly, despite the different levels of defect in TFIID assembly and promoter binding, we could not detect any strong global effect on Pol II transcriptional activity in both TAF7 and TAF10 depleted cells. Nevertheless, newly transcribed RNA-seq analyses showed that depletion of TAF10 resulted in more downregulated genes than depletion of TAF7, but with only hundreds of transcripts being affected.

### Different requirements for TAF7 and TAF10 during development

Both *Taf7* and *Taf10* null mutant embryos die around implantation ^37,38^, indicating that both TAF7 and TAF10 are similarly important for early mouse development. Nevertheless, we observed a difference in phenotype severity between TAF7 or TAF10 depletion. Our data indicate that TAF10 is important for the survival of the mESCs, as already suggested by the increased apoptosis observed in *Taf10* mutant F9 cells and in the inner cell mass of *Taf10^-/-^*mutant blastocysts ^37,51^. On the contrary, TAF7 is not essential for the survival of the mESCs, in agreement with the observation that *Taf7^-/-^* blastocyst can survive until hatching at E4.5 ^38^. Importantly, as TAF10 is also part of the transcriptional co-activator complex SAGA, we confirmed that the difference of phenotype in the *RT10* mESCs cells is not a consequence of a SAGA defect. We already showed that in absence of TAF10, SAGA complex assembly is disrupted ^35^. However, deletion of *Supt7l* coding for the HFD partner of TAF10 in SAGA did not increase the severity of the TAF7 depletion phenotype. Remarkably, we also observed that TAF10 was required for the stability of SUPT7L, as it is for TAF8 ^35,43^. It is tempting to speculate that TAF10 is also required for the stability of its other HFD partner in lobe A; TAF3. As a consequence, on one hand, destabilization of TAF8 would lead to the loss of TAF2 from lobe B and on the other hand, loss of TAF3 is predicted to impair integration of TAF11/TAF13 in the lobe A thus preventing TBP integration. The severity of this domino effect in absence of TAF10 potentially explains the stronger phenotype observed in TAF10-depleted mESCs and *TCre;Taf10* embryos

We interpret the increased severity in TAF10-depleted cells compared to TAF7-depleted cells by a different impact on TFIID assembly (see below). However, in the *Taf7* and *Taf10* conditional mutant embryos we observed phenotypes which were specific of each TAF. Indeed, while no limb bud could be observed in the *TCre;Taf10* mutant embryos, the *TCre;Taf7* mutant embryos developed forelimbs, although smaller compared to the controls. Limb bud specification is the result of a genetic cascade that is initiated by the activation of *Tbx5* (for the forelimb) in the lateral plate mesoderm which will result in the activation of *Fgf8* in the future apical ectodermal ridge (AER) and the establishment of a feed forward loop between the AER and the underlying mesenchyme that supports limb bud elevation ^52^. The presence of forelimbs in the *TCre;Taf7* embryos indicates that the genetic cascade leading to the limb bud specification and elevation is initiated whereas it is not in the *TCre;Taf10* mutant embryos as shown by the lack of *Fgf8* induction in the ectoderm ^35^.

We observed two striking different phenotypes: enlarged pericardium in the *TCre;Taf7* embryos and a lack of blood cells in the *TCre;Taf10* embryos. This suggests that TAF7 and TAF10 are associated with specific regulatory functions in spite of the fact that TAF7 and TAF10 are both subunits of holo-TFIID. This might suggest that TAF7-less TFIID cannot regulate genes necessary for cardiovascular development, while the production of red blood cells is more dependent on TAF10-regulated pathways as supported by the direct interaction between TAF10 and GATA1 during erythropoiesis ^6^. Importantly the lack of TAF10 seriously affects both holo-TFIID and -SAGA assemblies, but does not affect many gene regulatory pathways important for other developmental processes. This observation is in line with our finding that partial TFIID complexes can still regulate Pol II gene expression from many developmental gene promoters.

### Sequential assembly of holo-TFIID

Holo-TFIID is an about 1 MDa complex composed of 14 subunits, out of which some of them are present twice in the complex. How such a big complex can assemble in the cells is major biological question. Studies using overexpression of a combination of different TAF proteins have proposed that assembly of holo-TFIID is the result of the sequential incorporation of different TFIID submodules: a core-TFIID composed of two copies of TAF5, TAF6 and TAF9, a building block composed of a TAF2, TAF8 and TAF10 called 3-TAF, and a TAF11, TAF13 and TBP submodule ^11,14,18^. Recently, we have sown that TAF1 is a scaffold for the sequential incorporation of different cytoplasmic building blocks leading to the formation of a holo-TFIID complex that is immediately imported into the nucleus ^13^. Our data support the sequential assembly model for holo-TFIID as we observed that TFIID formation is arrested at different levels in our TAF7 or TAF10-depleted cells. Indeed, our IP-MS analyses indicate that in absence of TAF7, a TAF7-less TFIID complex containing TBP, is formed. This is in agreement with the fact that TAF7 is the last subunit to be incorporated on TAF1 ^13^. On the contrary, in TAF10 depleted cells, we could detect only the core-TFIID complex composed of TAF4, TAF5, TAF6, TAF9 and TAF12. In normal conditions, the next step after holo-TFIID assembly is the recruitment of the 3-TAF module ^11,14^. In absence of TAF10, as it is required for the stability of its HFD partners (this study and ^35,43^), there is no 3-TAF complex formed. Altogether, our data confirm that *in vivo*, the holo-TFIID complex is sequentially assembled from different building blocks.

### Active Pol II transcription in TAF7 and TAF10 depleted mESCs

A striking observation from our study is that despite an obvious impact on TFIID when TAF7 or TAF10 are depleted, we did not observe a major impact on nascent Pol II transcription. We already reported that conditional deletion of *Taf10* during somitogenesis did not have an impact on the active transcription of cyclic genes associated with the formation of the somites ^35^. In this precedent study, we had performed a transcriptomic analysis of steady state RNA which showed a very limited effect but we could not exclude a compensation of transcription initiation defects by a buffering of RNA decay as already proposed in yeast ^53,54^. In the present study, we showed the lack of obvious impact on active transcription by analyzing EU run-on experiments, phosphorylated Pol II and nascent 4-sU-RNA-seq. Only a very limited proportion of expressed transcripts were significantly and differentially affected in TAF7 or TAF10-depleted cells after 3 days of 4-OHT treatment when our IP-MS data showed that the assembly of TFIID is impaired, although analysis of the Log2 fold change distributions indicates that there is nevertheless a slight global down-regulation shift when TAF10 is depleted, correlating with the more severe phenotype observed in TAF10 depleted cells. We also showed that while TAF7 or TAF10 depletion did not affect cell viability when passaged at day 2 of treatment, clonal growth was affected when *RT7* and *RT10* cells were passaged at day 4 of treatment (Figure 1I, J). These data indicate that TAF7 and TAF10 depletions are actually damaging for the mESCs but that during a window of time, the cells are able to compensate.

These surprising observations can be explained by several mutually non-exclusive hypotheses. A first hypothesis could be that residual low abundant holo-TFIID complexes in TAF7 or TAF10-depleted cells might be responsible for the maintenance of active transcription. Indeed, although the depletion of TAF7 or TAF10 is very efficient as shown by our western blot analysis, even from chromatin extracts, we still detect some residual proteins in our IP-MS experiments that could be explained by the stabilization of either TAF7 or TAF10 within holo-TFIID as we already observed for TAF10 in E9.5 mouse *TCre;Taf10* embryos ^35^. It is conceivable that chromatin bound holo-TFIID has a longer half-life and is very difficult to destroy in the time frame of our studies. This would suggest that only a very minimal amount of holo-TFIID maybe required to maintain active transcription in a steady-state situation. However, as the mESCs are still dividing, especially in the TAF7-depleted cells which are almost indistinguishable from the control *RT7* cells up to 6 days of treatment, remaining chromatin-bound TFIID complexes would be dislodged during DNA synthesis and diluted out in the cell population. Thus, this hypothesis cannot solely explain our observations.

A second hypothesis could be that holo-TFIID is not required to maintain active transcription as proposed in ^40^. During the very first round of Pol II transcription initiation, TFIID would be required to create a scaffold that would allow some GTFs to remain at the promoter facilitating re-initiation and the recruitment of new Pol II ^55,56^. This model would not necessarily require fully assembled holo-TFIID, maybe only TBP, for re-initiation of transcription. To further test this hypothesis, it would be interesting to differentiate our *RT7* and *RT10* mESCs to evaluate the impact of the depletion of TAF7 or TAF10 on the activation of a specific new gene expression program. It would be important to test differentiation in various cell types as it is already established that the loss of TAF10 has a differential effect depending on the cellular context ^35,37,39,40^.

A third hypothesis could be that mESCs are not using a holo-TFIID complex. It has been proposed that human ES cells (hESCs) have a non-canonical TBP-containing TAF complex as expression of several TAF proteins including TAF10, but not TAF7 were not detected ^57^. However, as shown in the same study and in our present data, this is not the case in mESCs. Differential requirements for TAF4, TAF8 and TAF12 have been reported as *Taf4* deletion is not impacting growing mESCs on the contrary to *Taf8* conditional deletion and TAF12 depletion ^31,58,59^. Interestingly, depletion of TAF8 and TAF12 leads to a strong defect in holo-TFIID assembly ^58,59^. Our data are in favor of a TAF7-less TFIID complex almost as active as the holo-TFIID, as we did not observe major difference in the proportion in TAF proteins, except TAF7, in our anti-TAF12 or anti-TBP IP-MS and in our CUT&RUN analyses. These data are in agreement with the observation that TAF7 is not required for the survival and the differentiation of mature T cells and that in these conditions, an apparent TAF7-less TFIID was observed, although the efficiency of the depletion was only about 50%^38^.

### Transcription initiation by partial TFIID complexes

Recent structural studies have proposed that TFIID is playing a major role in PIC assembly and deposition of TBP on core promoters ^10,12,60^. While TBP has been initially characterized as the TATA-box binding protein ^61^, only a minority of mammalian promoters have a TATA box ^62^ and TFIID is able to deposit TBP on TATA box-containing or TATA-less promoters, leading to a similar bending of the DNA ^10^. In this model, TBP plays a major role as an anchor on the DNA, allowing the positioning of the PIC independently of the sequence of the promoter ^63^. Therefore, TFIID and TBP appear central to the process of Pol II transcription initiation. However, several evidences in the literature point out that Pol II transcription initiation does not always rely on TBP or TFIID. It has been shown that the TBP-free TAF-complex (TFTC), a mixture of SAGA and other TAF10-containing TAF complexes ^64^, is able to initiate transcription *in vitro* on TATA-containing and TATA-less promoters ^65^. Pol II transcription initiation of snRNA coding genes is controlled by the SNAPc complex which recruits TBP on the promoters without TFIID ^66^. In *Drosophila*, *H1* and ribosomal protein coding genes transcription initiation is controlled by a non-TFIID complex containing the TBP paralog TBPL1 ^67–70^ and different *Drosophila* PIC complexes not containing TBP and associated with different classes of promoters have also been described recently ^71^. In the mouse, we recently showed that during oocyte growth, transcription is mediated by an oocyte-specific complex containing the TBP paralog TBPL2, lacking TBP and TAF proteins ^72^. More strikingly, TBP and TBP paralogs independent transcription has been described during Xenopus development ^73^ and two recent studies in mESCs or in human HAP1 cells have shown that acute depletion of TBP has no major impact on Pol II transcription without compensation by a TBP paralog ^74,75^.

Our data are in favor of transcription initiation mediated by partially assembled TFIID complexes in the mESCs. On one hand, our data strongly support the idea that the TAF7-less TFIID complex is able to mediate transcription initiation, at least up to a certain degree as the TAF7-depleted mESCs are sensitive to late replating and as the *TCre;Taf7* embryos have a phenotype. On the other hand, we show that a partially assembled core-TFIID-like complex in absence of TAF10 can maintain the transcription activity of many genes (Figure 6). It has been recently shown that an *in vitro* assembled lobe C-like single TAF (S-TAF) complex composed of TAF1, TAF7, TAF11, TAF13 and TBP is able to promote transcription initiation *in vitro* ^19^. However, we could not detect strong interaction between TBP and TAF1, TAF7, TAF11 or TAF13 in our anti-TBP IP-MS, suggesting that the S-TAFs complex does not exist *in vivo* in TAF10 depleted cells. As shown in the structural data, the core-TFIID does not interact strongly with TBP, as TBP interacts with TAF11/13 and TAF1 ^10,12,15,18,60,76^. Our data confirmed this observation as in our anti-TBP IP-MS in the TAF10 depleted cells, contrary to the TAF7-depleted cells, there is a strong decrease interaction between TBP and the TAF proteins which is confirmed by the relocation of TBP to Pol III and at a lesser extent Pol I machineries (Sup Figure 3). Moreover, it is not clear how the core-TFIID could be imported in the nucleus as it has been proposed that the NLS of TAF8 which is incorporated in the 8-TAF complex is important for this import ^11,14,45^. As an alternative, we recently showed that TAF1 acting as a scaffold is able to assemble holo-TFIID in the cytoplasm prior its fast import in the nucleus ^13^. As TAF1 is able to interact in the structure with TAF6 ^10,13^, a tempting speculation is that the core-TFIID complex is able to be recruited on the nascent TAF1 protein which already interacts with TBP. Such a complex could potentially interact with the DNA via TAF4 and TAF1, as well as TBP as these proteins are making contacts with the DNA in the holo-TFIID/PIC structure ^10^. The apparent size of 670 kDa of the partial TFIID complex in our gel filtration experiment supports the existence of such a complex. However, this interaction might be weak as we did not have enough sensitivity to detect this interaction in our IP-MS experiment, and this could be explained by the absence of TAF8.

In conclusion, our study supports the vision of a much more dynamic and flexible Pol II transcription initiation machinery. Interestingly, this flexibility has some impact on human physiopathology as a growing number of mutations have been identified in *TAF1*, *TAF2*, *TAF4*, *TAF6*, *TAF8* and *TAF13* genes, in patients associated with neurodevelopmental defects and intellectual disabilities ^58,77,78^. Surprisingly, these mutations mainly affect the brain development and function but without any major effect on the other organs, clearly suggesting that the transcription initiation machinery has a certain degree of adaptability but that the cellular context is a main constraint.

### Limitation of the Study

In our study, we tested the functional importance of TAF7 and TAF10 using an inducible CRE/*Lox*P approach. While the genomic deletion is very efficient and fast, this approach does not target the proteins that have already been translated, on the contrary of an acute depletion approach. However, we evaluated the potential persistence of some TAF7 or TAF10 protein at different time points, by IF, western blot and by IP-MS. Moreover, we have analyzed the status of the TFIID complex to confirm the impact of the depletion of TAF7 or TAF10.

As our study is more long term compared to an acute depletion approach, we decided to perform our transcriptomic and genomic analyses at day 3 which corresponds to an efficient depletion of the targeted proteins and to a stage where no important cellular phenotype could be observed, however, as our cells have been depleted of TAF7 or TAF10 for days, we cannot rule out any secondary effects although, as we show, there are no major defects in transcription. This does not rule out the total absence of phenotype as cells become affected on the longer term but our study does not allow the analysis of the longer-term defects.

## Supporting information

figures_plus_legend

supplementart figures_plus_legend

## STAR Methods

See Supplementary data.

## Data availability

Proteomic data have been deposited in the proteomics identification database (ProteomeXchange PRIDE database, accession code: PXD046459). Transcriptomic and genomic data have been deposited on the GEO portal (GSE245196)

## Acknowledgments

We thank all members of the Tora lab for protocols, thoughtful discussions and suggestions, A. Bernardini for thoroughly reading and commenting the manuscript, C. Ebel and M. Philipps for help with FACS sorting, S. Falcone, Michael Gendron and the IGBMC animal facility for animal caretaking and the IGBMC cell culture facility.

## Author’s contributions

VH, PB, DD, EGS, CR, MS, YH, CE, LHA performed experiments. BM and LN performed the mass spectrometry analyses, MJ, SLG, VH, YH and SDV performed the bioinformatic analyses, VH, LT, and SDV designed the experiments with the help of FM. SDV conceived, supervised the project and wrote the paper with support from VH and LT.

## Supplemental Figure legends

**Figure S1:**
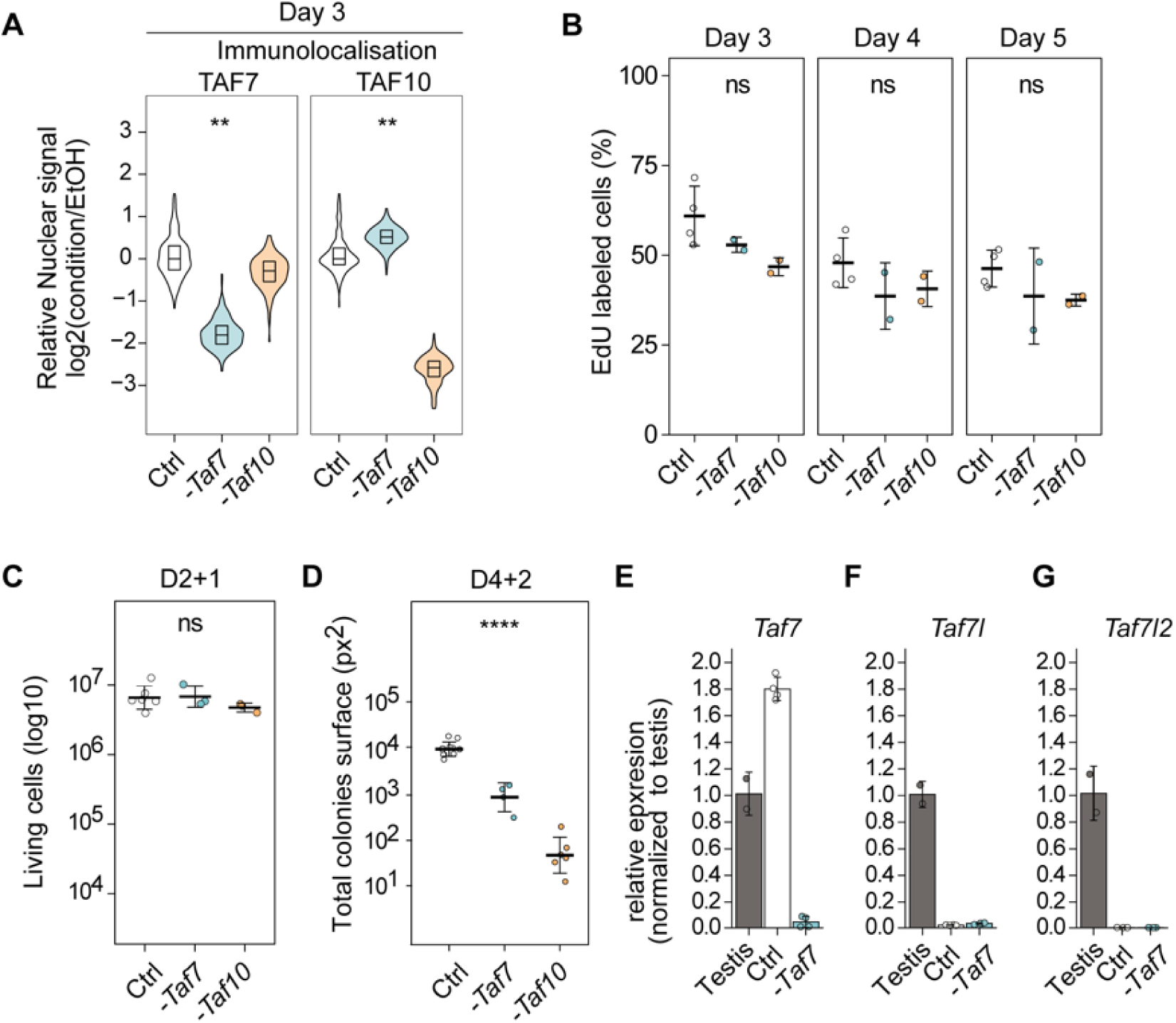
TAF7 depletion in mESCs results in a milder phenotype compared to TAF10 depletion. **(A)** Nuclear quantification of anti-TAF7 and anti-TAF10 immunolocalization signal in control condition (Ctrl) or after *Taf7* (-*Taf7)* or *Taf10* deletion (*-Taf10)* after 3 days of treatment. **(B)** Percentage of cells in S phase assessed by 5-ethynyldeoxyuridine (EdU) staining at 3, 4, and 5 days of treatment. For D3 to D5, Ctrl; n = 4, *-Taf7;* n = 2, *-Taf10*; n = 2 biological replicates. **(C,D)** Quantifications of the ability of the *RT7* or *RT10* depleted cells to growth after cell passage. Living Trypan blue negative cell numbers after cell passage at D2 and analyzed at D3 (D2+1). Ctrl; n = 6, *-Taf7;* n = 3, *-Taf10*; n = 3 biological replicates (C). Total colonies surfaces estimated by Crystal violet staining after cell passage at D4 and analyzed at D6 (D4+2). Ctrl; n = 10, *-Taf7;* n = 4, *-Taf10*; n = 6 biological replicates (D). For all panels, the control condition (Ctrl) corresponds to all *RT7* and *RT10* cells treated with EtOH. **(E-G)** RT-qPCR analysis of *Taf7, Taf7l* and *Taf7l2* expression from control (Ctrl), mutant *(-Taf7) RT7* mESCs at day 4 and wild type mouse testis RNA. The RNA polymerase III transcribed gene *Rn7sk* was used as reference gene and the data were normalized to the testis. n=2 biological x 2 technical replicates, except for testis; n= 2 technical replicates. Means ± standard deviation are shown. (A-C) Kruskal–Wallis test: ns; not significant, * <0.05; **<0.01; *** <0.001; **** <0.0001.

**Figure S2:**
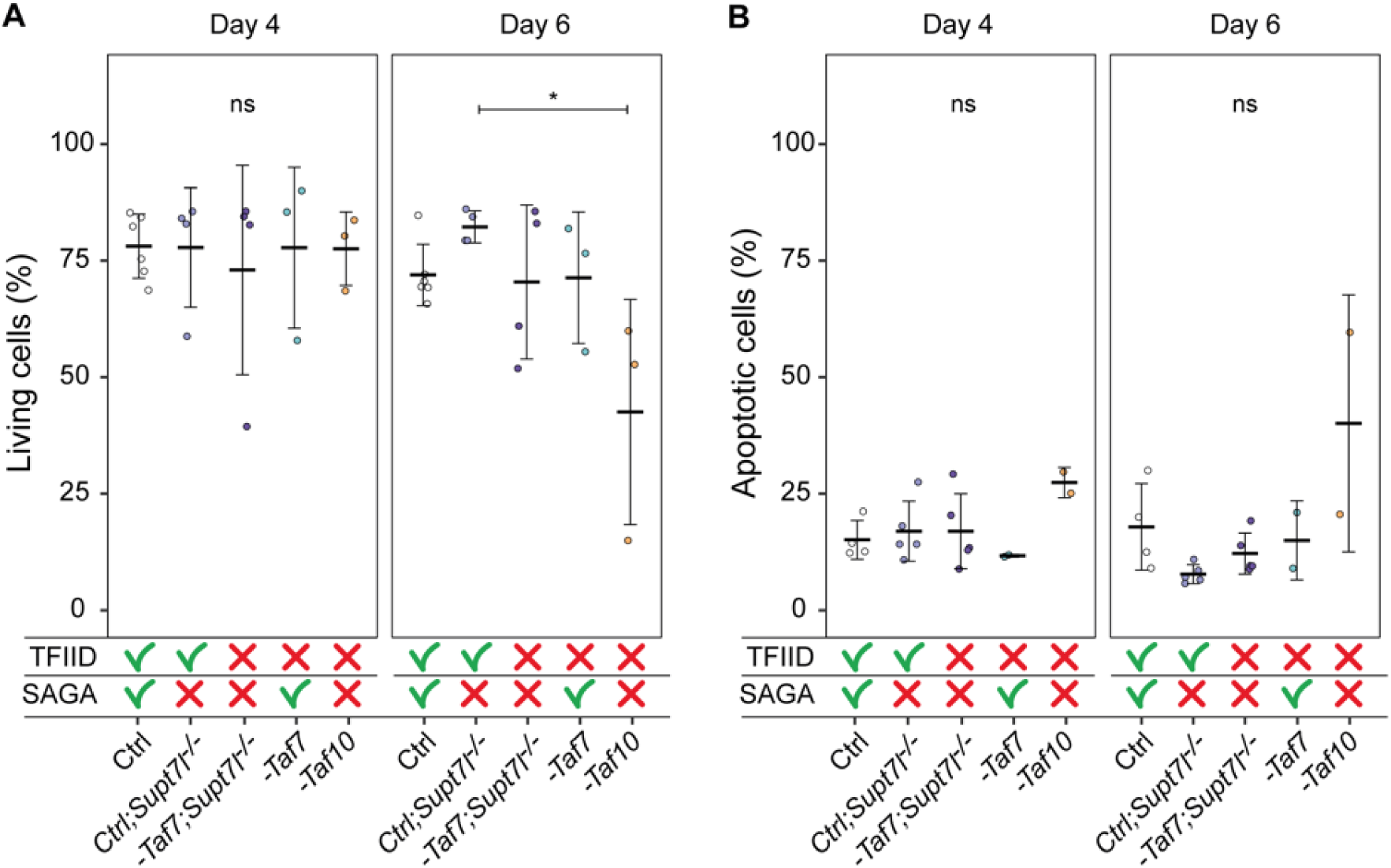
*Supt7l* loss of function in TAF7 depleted mESCs does no impact cell viability. **(A)** Percentage of living mESCs analyzed by Trypan blue staining after 4 and 6 days of treatment. Ctrl; n = 6, Ctrl;*Supt7l^-/-^*; n = 4, *-Taf7;Supt7l^-/-^*; n = 4, *-Taf7;* n = 3, *-Taf10*; n = 3 biological replicates per day. **(B)** Percentage of apoptotic (Annexin V positive and propidium iodide negative) mESCs after 4 and 6 days of treatment. Ctrl; n = 4, Ctrl;*Supt7l^-/-^*; n = 5, *- Taf7;Supt7l^-/-^*; n = 5, *-Taf7;* n = 2, *-Taf10*; n = 2 biological replicates per day. The impact on TFIID and SAGA assembly is indicated at the bottom. Kruskal–Wallis test followed by a post hoc Dunn test if significant: ns; not significant, * <0.05.

**Figure S3:**
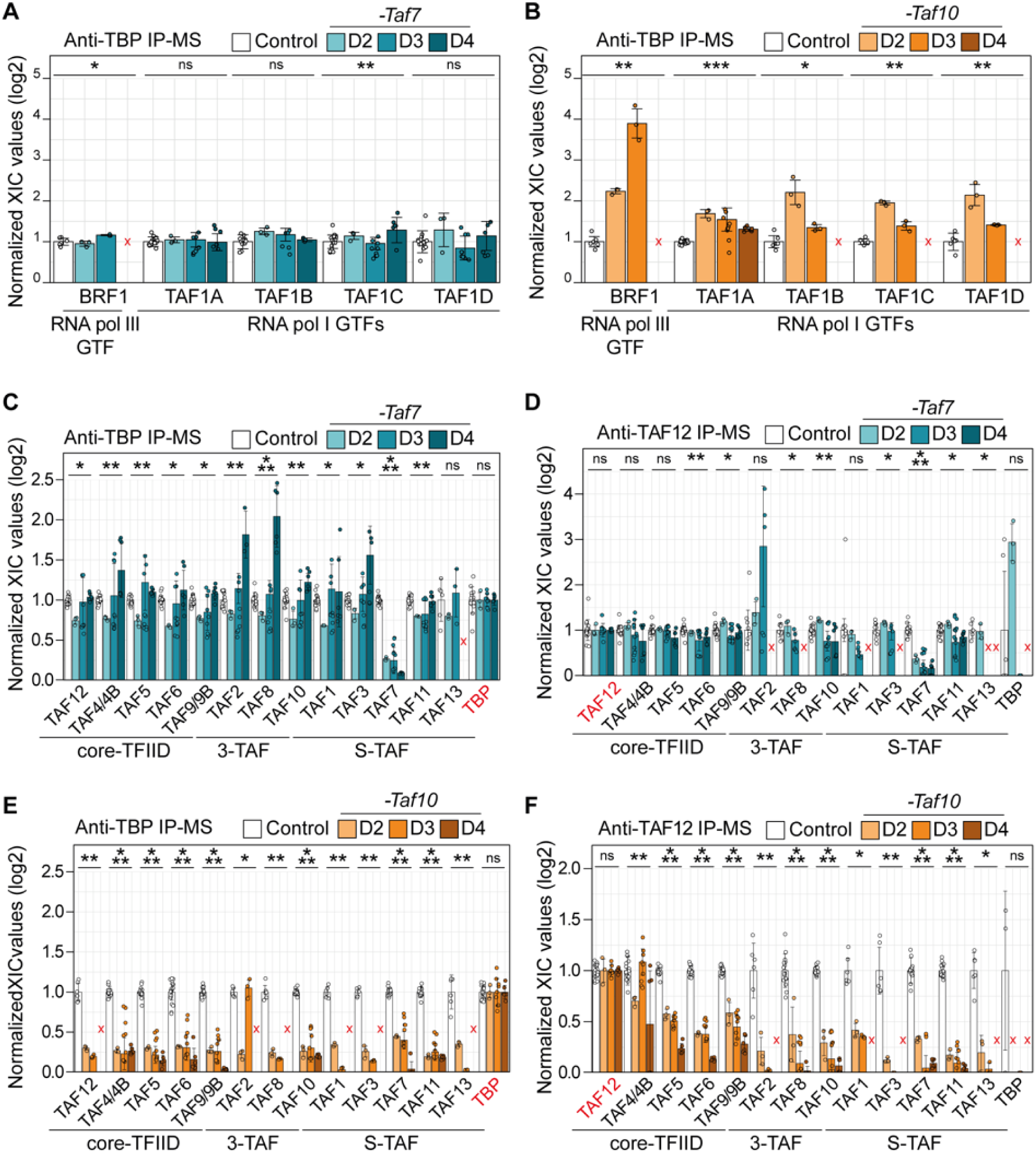
Different effects on holo-TFIID assembly in TAF7- and TAF10-depleted mESCs. **(A-F)** Normalized relative XIC values for RNA polymerase I and III GTFs subunits (A, B) and TFIID subunits (C-F) after anti-TBP (A-C,E) or anti-TAF12 (E,F) IP-MS. For each IP-MS, the XIC values of each subunit were first normalized to the mean XIC of the bait protein and then normalized to the mean of the respective control condition (EtOH treated *RT7* for mutant *-Taf7*, EtOH treated *RT10* for mutant *-Taf10)*. Analyses were performed on day 2, day 3, and day 4 of treatment. Red crosses indicate proteins not detected in the control condition. Day 2; n = 1 biological replicate x 3 measures, D3; 3x3, D4; 2x3. Means ± standard deviation are shown. Kruskal-Wallis test. ns; not significant, * <0.05; **<0.01; *** <0.001.

**Figure S4:**
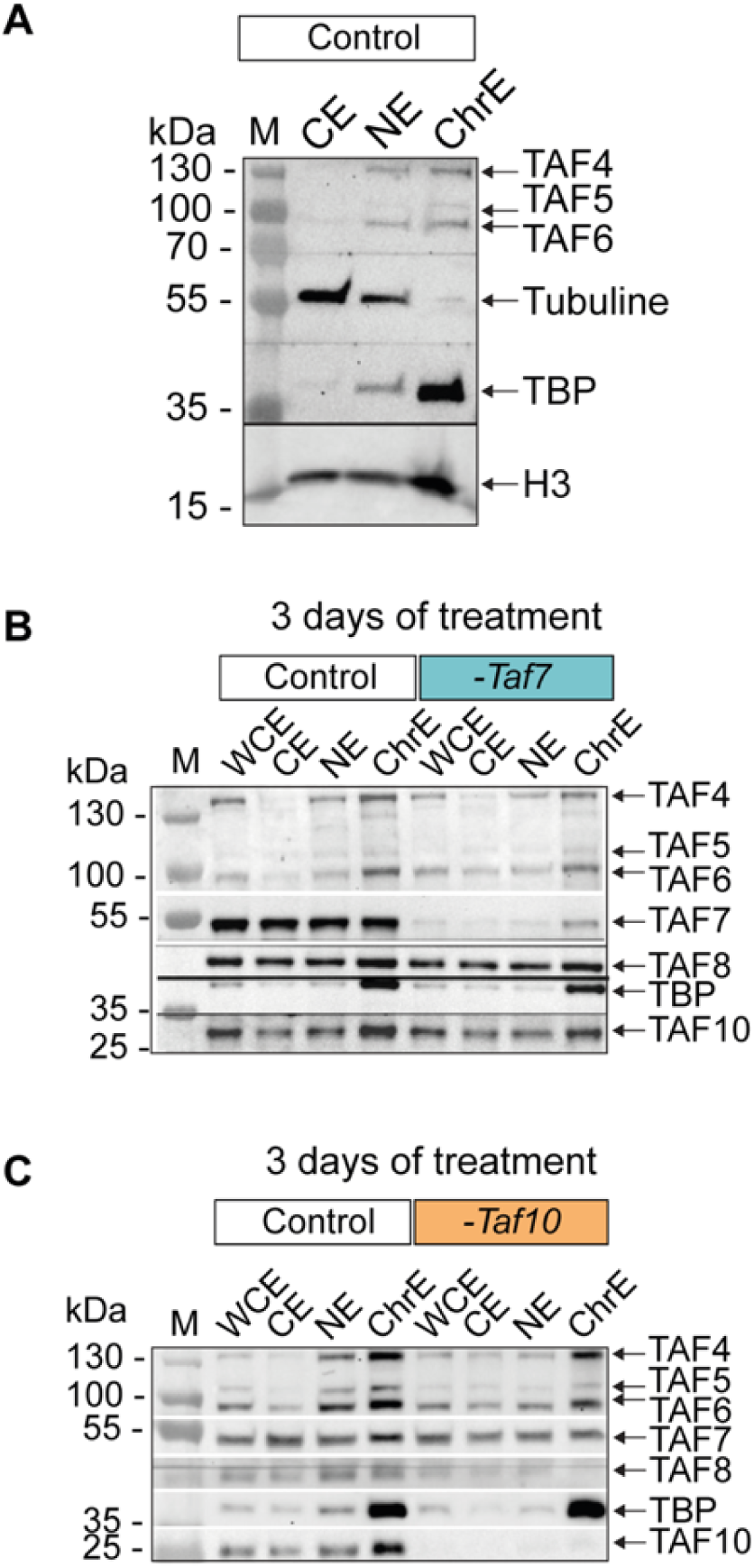
TBP recruitment on the chromatin of TAF7- and TAF10-depleted mESCs. **(A)** Western blot analysis on cytoplasm-enriched extract (CE), nuclear-enriched extract (NE), and chromatin-enriched extract (ChrE) obtained from untreated mESCs. The position of the different proteins probed is indicated on the right and the molecular weights on the left. **(B,C)** Western blot analyses of whole cell extract (WCE), CE, NE and ChrE from 4-OHT treated *RT7* (-*Taf7*, C) and 4-OHT treated *RT10* (-*Taf10*, D) depleted mESCs and their respective controls (Control, EtOH treated cells) at day 3. The position of the different proteins probed is indicated on the right and the molecular weights (M) on the left.

**Figure S5:**
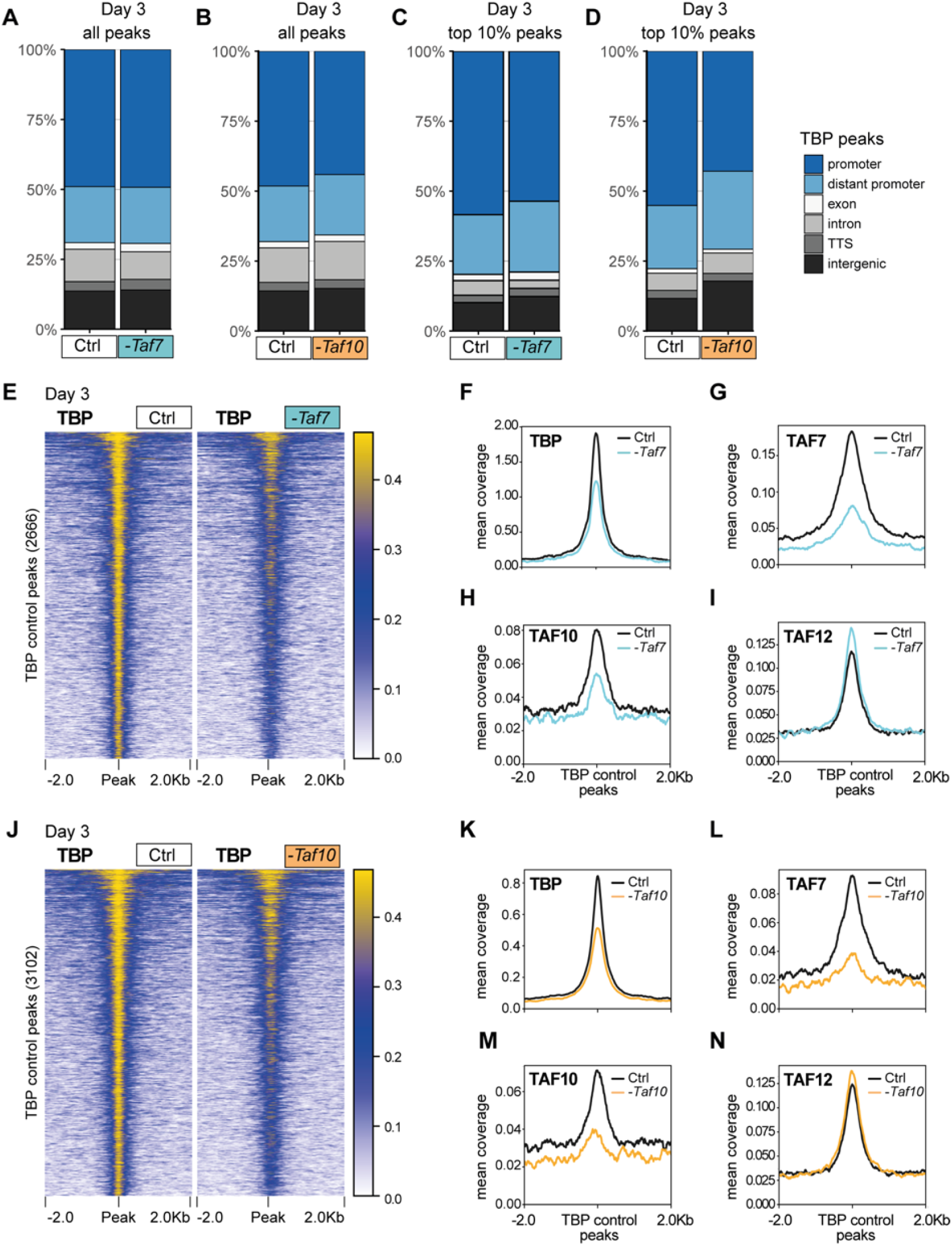
TBP recruitment on the chromatin of TAF7- and TAF10-depleted mESCs. **(A-D)** All (A,B) and top10% (C,D) TBP CUT&RUN peak annotation in *RT7* (A,C) and *RT10* (B,D) control (Ctrl) and depleted (-*Taf7* and -*Taf10*) mESCs. The legend of the annotations is indicated on the right. **(E)** Heatmap showing the TBP distribution at the position of the TBP peaks (± 2 kb) detected in the control condition, in control (Ctrl) and mutant *RT7* (-*Taf7*) mESCs at day 3. **(F-I)** Mean profile of TBP (F), TAF7 (G), TAF10 (H) and TAF12 (I) protein accumulation at the location of the TBP control peaks, in control (black line) and in TAF7- depleted (blue line) mESCs. **(J)** Heatmap showing the TBP distribution at the position of the TBP peaks (± 2 kb) detected in the control condition, in control (Ctrl) and mutant *RT10* (- *Taf10*) mESCs at day3. **(K-N)** Mean profile of TBP (F), TAF7 (G), TAF10 (H) and TAF12 (I) protein accumulation at the location of the TBP control peaks, in control (black line) and in TAF10-depleted (orange line) mESCs.

**Figure S6:**
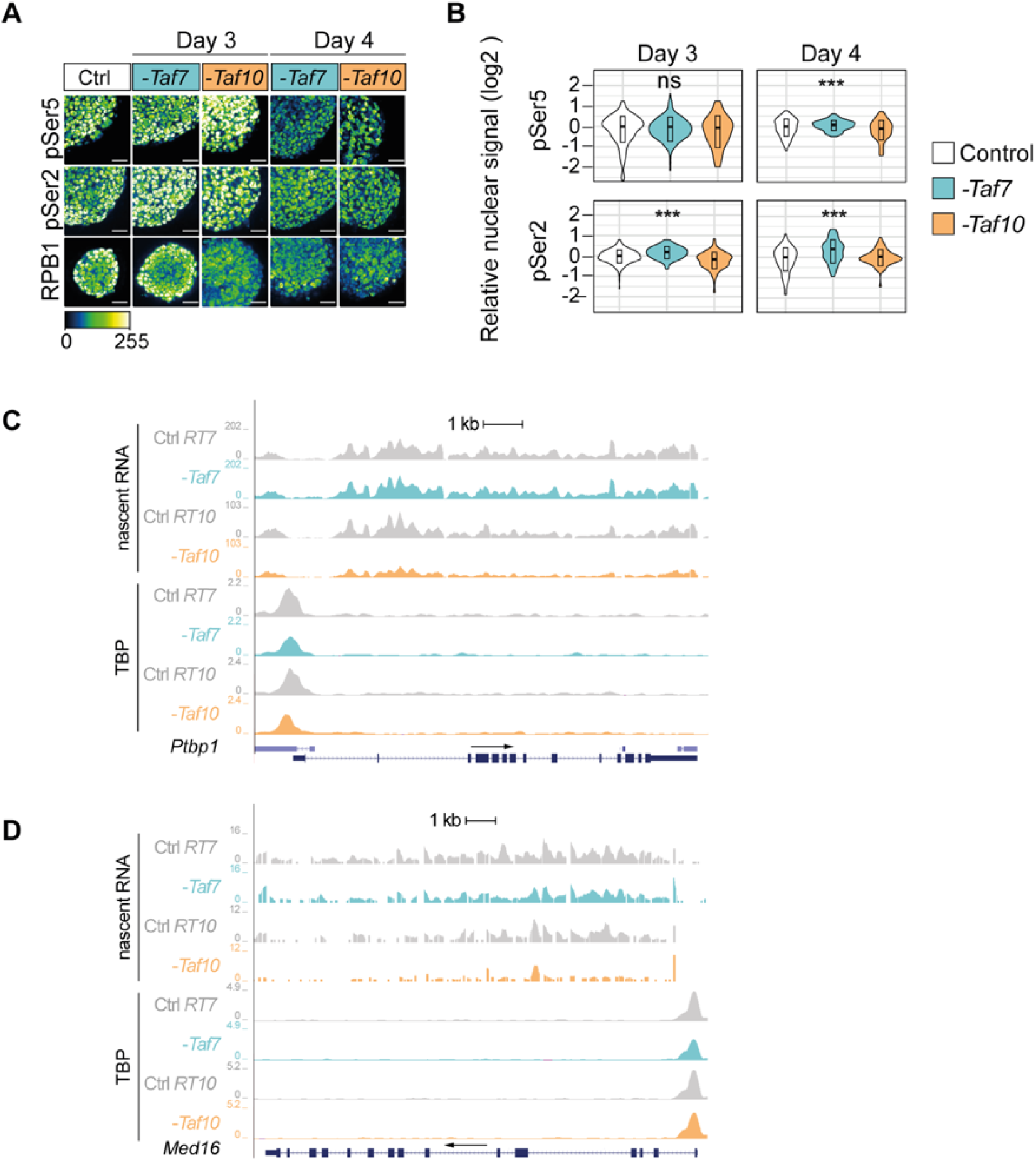
RNA Pol II global activity is not affected in TAF7- and TAF10-depleted mESCs. **(A)** Representative views of immunofluorescence using anti-RPB1 (bottom), anti- RPB1^pSer2^ (middle) and anti-RPB1^pSer5^ (top) antibodies on *RT7* and *RT10* mESCs treated 3 or 4 days with 4-OHT (-*Taf7* or -*Taf10*) and with EtOH (Ctrl). Color scale (Green Fire Blue LUT scale) is indicated at the bottom of the panel, scale bar: 15 µm. **(B)** Quantifications of RPB1^pSer5^ and RPB1^pSer2^ nuclear signal represented as violin plots. D3 and D4; n=1 biological replicate. Hundreds of measurements for individual nuclei were performed for the quantification, scale bar: 50 µm. Kruskal-Wallis test. ns; not significant, * <0.05; **<0.01; *** <0.001. **(C,D)** UCSC genome browser views of nascent RNA (top) and TBP distribution (bottom) at *Ptbp1* (C) and *Med16* (D) loci between control (Ctrl *RT7* and Ctrl *RT10*), TAF7 depleted (-*Taf7*) and TAF10 depleted (-*Taf1*0) mESCs. Y axes indicate the genomic coverage. Arrows indicate direction of transcription.

**Figure S7:**
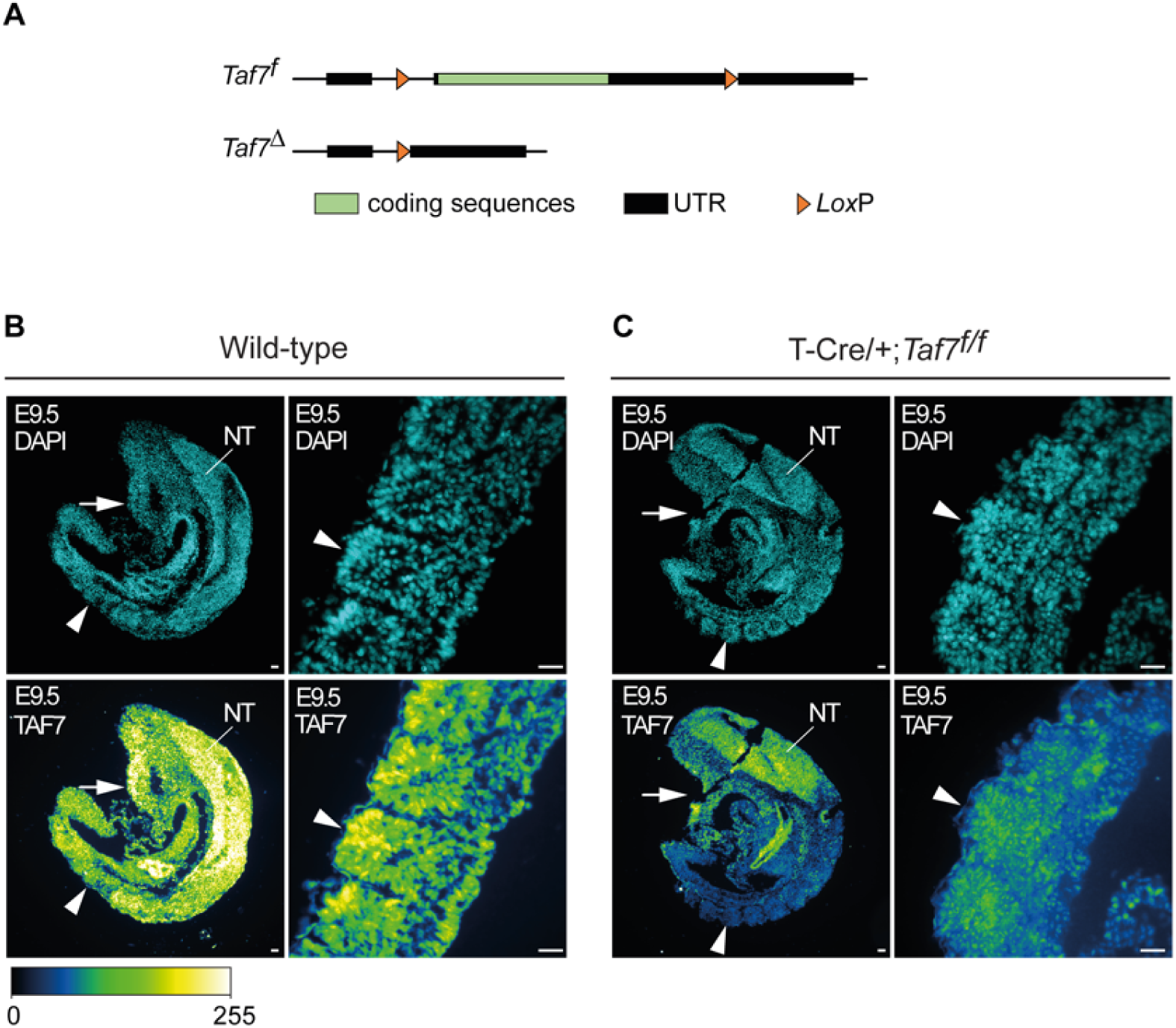
Efficient TAF7 depletion in the paraxial mesoderm of embryonic day (E) 9.5 Tg(T-Cre/+)*;Taf7^f/f^* mutant embryos. **(A)** Map of the *Taf7* conditional (*Taf7^f^*) and deleted (*Taf7*^Δ^) alleles after CRE recombination. UTRs and ORF are indicated in black and green boxes, respectively. *Lox*P sites are indicated by orange arrowheads. **(B-C)** DAPI counterstained anti-TAF7 immunofluorescence on sagittal sections from E9.5 wild-type (B) and Tg(T-Cre/+)*;Taf7^f/f^ mutant* (C) embryos. In Tg(T-Cre/+) embryos, CRE is active in progenitors that contribute to mesoderm cells posterior to the heart (white arrow), including the paraxial mesoderm (unsegmented presomitic mesoderm and somites (arrowhead)) but not in the neural tube (NT). Color scale (Green Fire Blue LUT scale) is indicated at the bottom (n=2), scale bar: 30 µm.

