## Supplementary material for "RNA polymerase II transcription with partially assembled TFIID complexes": material_methods

**STAR * METHODS**

**KEY RESSOURCES TABLE**

**Antibodies**

| REAGENT or RESSOURCE | SOURCE | IDENTIFER |
|---|---|---|
| mouse monoclonal anti-TAF4 (WB) | In house | 32TA-2B9, [1] |
| mouse monoclonal anti-TAF5 (WB) | In house | 1TA-1C2, [2] |
| mouse monoclonal anti-TAF6 (WB) | In house | 25TA-2G7, [3] |
| mouse monoclonal anti-TAF7 | In house | 19TA-2C7, [3] |
| rabbit polyclonal anti-TAF7 (WB, IF, CUT&RUN) | In house | 3407, [4] |
| mouse monoclonal anti-TAF7L (WB) | In house | 46TA-2D5, [5] |
| mouse monoclonal anti-TAF8 (WB) | In house | 3478, [4] |
| mouse monoclonal anti-TAF10 (WB, IF, CUT&RUN) | In house | 6TA-2B11, [1] |
| mouse monoclonal anti-TAF12 (WB, IP, CUT&RUN) | In house | 22TA-2A1, [3] |
| mouse monoclonal anti-TBP (WB, IP) | In house | 3TF1-3G3, [6] |
| rabbit polyclonal anti-TBP (CUT&RUN) | Abcam | ab28175 |
| mouse monoclonal anti-SUPT7L (WB) | Bethyl laboratories | A302-803A |
| rat monoclonal anti- RPB1pSer5 (IF) | GmbH antibody service | CTD4-3E8, [7] |
| tat monoclonal anti- RPB1pSer2 (IF) | GmbH antibody service | CTD4-3E10, [7] |
| mouse monoclonal anti-GST (IP) | In house | 15TF2-1D10, [8] |
| Alexa Fluor® 488-labelled goat anti-rabbit IgG | Life Technologies | A-11008, RRID: AB_143165 |
| Alexa Fluor® 546-labelled goat anti-mouse IgG | Life Technologies | A-11003 RRID: AB_2534071 |
| Alexa Fluor® 488-labelled goat anti-rat IgG | Life Technologies | A-11006, RRID: AB_2534074 |
| Peroxydase AffiniPure goat anti-rabbit IgG(H+L) | Jackson ImmunoResearch | 111-035- 144, RRID: AB_2307391 |
| Peroxydase AffiniPure F(ab')$_2$ Fragment goat anti-mouse IgG, Fcγ specific | Jackson ImmunoResearch | 111-036- 071, RRID: AB_2338524 |

WB; western blot, IP; immunoprecipitation, IF; immunofluorescence

**Chemicals, Peptides, and recombinant proteins**

| REAGENT or RESSOURCE | SOURCE | IDENTIFER |
|---|---|---|
| 4′,6-diamidino-2-phenylindole dihydrochloride (DAPI) | Molecular Probes | D1306 |
| 4-Hydroxytamoxifen (4-OHT) | Sigma Aldrich | H7904 |
| 4-Thiouridine (4-sU) | Glentham Life Science | GN6085 |
| 5-Ethynyl Uridine (5-EU) | | |
| BlueTrypan Staining 0.4% | Invitrogen | T10288 |
| CHIR99021 | Axon Medchem | 1386 |
| Crystal Violet | Sigma Aldrich | C3886-25G |
| Complete Protease Inhibitor Cocktail (cOmplete), EDTA free | Roche | 11873580001 |

| | | |
|---|---|---|
| EZ-link™ HPDP-Biotin | Thermo Scientific | 21341 |
| Flavopiridol | Sigma Aldrich | F3055-1mg |
| Leukemia inhibitory factor (LIF) | In house | NA |
| Pierce™ ECL Western Blotting Substrate | ThermoFisher Scientific | 32109 |
| Protein G Sepharose | GE healthcare | 17-0618-05 |
| Random hexamer primer | Thermo Scientific | SO142 |
| RNasin (40U/µL) | Promega | N2111 |
| RiboPure – Yeast Kit | Invitrogen | AM1926 |
| TRI® Reagent (Trizol) | Molecular Research Center Inc. | TR188 |
| Triptolide | Sigma Aldrich | T3652-1mg |
| VECTASHIELD® Mounting Media without DAPI | VectorLabs | H-1000 |

**Critical Commercial Assays**

| REAGENT or RESSOURCE | SOURCE | IDENTIFER |
|---|---|---|
| AMPure XP beads | Beckman-Coulter | A63882 |
| Click-it RNA Imaging Kits | Invitrogen | C10329 |
| EdU Staining Proliferation Kit | Abcam | ab222421 |
| LightCycler® 480 SYBR® Green 2x PCR Master Mix I | Roche | 4887352001 |
| MicroPlex library preparation kit v3 | Diagenode | C05010001 |
| NucleoSpin RNA XS, RNA extraction kit | Machery-Nagel | 740902.50 |
| Pierce™ ECL Western Blotting Substrate | ThermoFisher Scientific | 32109 |
| QuantiTect Reverse Transcription Kit | Qiagen | 205311 |
| Protein Assay Dye Reagent Concentrate | Bio-Rad | 5000006 |
| RiboPure – Yeast Kit | Invitrogen | AM1926 |
| SPRIselect beads | Beckman-Coulter | B23319 |
| SuperScript IV Reverse Transcriptase | Invitrogen | 18090050 |
| TRI® Reagent (Trizol) | Molecular Research Center Inc. | TR188 |
| TruSeq Stranded Total RNA LT Sample Prep Kit with Ribo-Zero Gold | Illumina | RS-122-2301 |
| µMACS Streptavidin Kit | Miltenyi Biotec | 130-074-101 |

**Deposited Data**

| REAGENT or RESSOURCE | SOURCE | IDENTIFER |
|---|---|---|
| Mass-spectrometry proteomics TFIID complex anti-TAF12 IPs from WT and TAF7 or TAF10 depleted mESCs | This study | PRIDE PXD046459 |
| Mass-spectrometry proteomics TFIID complex anti-TBP IPs from WT and TAF7 or TAF10 depleted mESCs | This study | PRIDE PXD046459 |
| Nascent transcriptomic data from WT and TAF7 or TAF10 depleted mESCs | This study | GEO GSE245196 |
| CUT&RUN from WT and TAF7 or TAF10 depleted mESCs | This study | GEO GSE245196 |

**Experimental Models: Mouse Lines**

| REAGENT or RESSOURCE | SOURCE | IDENTIFER |
|---|---|---|
| Tg(*T-Cre*) | (Perantoni et al., 2005) | N/A |

| | | |
|---|---|---|
| $R26^{CreERT2}$ | [9] | MGI:3699244 |
| $Taf10^{f/f}$ | [1] | MGI:3606185 |
| $Taf7^{f/f}$ | [10] | MGI:5430373 |
| Tg(*T-Cre/+*);$Taf7^{f/f}$ | This study | N/A |
| Tg(*T-Cre/+*);$Taf7^{f/f}$ | [4] | N/A |

**Experimental Models: Cell Lines**

| REAGENT or RESSOURCE | SOURCE | IDENTIFER |
|---|---|---|
| $R26^{CreERT2/+}$;$Taf7^{f/f}$ | This study | N/A |
| $R26^{CreERT2/R}$;$Taf10^{f/f}$ | (Bardot et al., 2017) | N/A |
| $R26^{CreERT2/+}$;$Taf7^{f/f}$;$Taf10^{f/f}$ | This study | N/A |
| $R26^{CreERT2/+}$;$Taf7^{f/f}$;$Sup7l^{-/-}$ | This study | N/A |
| CD1 WT fibroblasts | IGBMC cell culture platform | N/A |

**Oligonucleotides**

| REAGENT or RESSOURCE | SOURCE | IDENTIFER |
|---|---|---|
| For primer sequences, see Sup. Table S1 | This study | N/A |
| For sgRNA Sequences, see Sup. Table S2 | This study | N/A |

**Software and Algorithms**

| REAGENT or RESSOURCE | SOURCE | IDENTIFER |
|---|---|---|
| Adobe Illustrator | Adobe | N/A |
| Bowtie2 | [11] | http://bowtie-bio.sourceforge.net/bowtie2/index.shtml |
| cutadapt | [12] | https://cutadapt.readthedocs.io/en/v1.10/ |
| deepTools | [13] | https://github.com/deeptools/deepTools |
| DESeq2 | [14] | https://bioconductor.org/packages/ |
| ImageJ | [15] | https://imagej.net/software/imagej/ |
| Image Lab Software | Bio Rad | https://www.bio-rad.com/fr-fr/product/image-lab-software?ID=KRE6P5E8Z |
| Homer | [16] | http://homer.ucsd.edu/homer/ |
| htseq-count | [17] | https://htseq.readthedocs.io/en/master/ |
| MACS2 | [18] | https://hbctraining.github.io/ |
| Proteome Discoverer 2.2 | ThermoFisher Scientific | https://www.thermofisher.com/fr/fr/home/industrial/mass-spectrometry/liquid-chromatography-mass-spectrometry-lc-ms/lc-ms-software/multi-omics-data-analysis/proteome-discoverer-software.html |
| R | R-project | https://CRAN.R-project.org/ |
| RStudio | RStudio | https://www.rstudio.com/categories/rstudio-ide/ |
| STAR | [19] | https://github.com/alexdobin/STAR |

### RESSOURCE AVAILABILITY

#### Lead contact

**Materials availability**

Cell lines generated in this study are available upon request.

**Data and code availability**

Nascent RNA-seq, CUT&RUN haven been deposited at GEO [20] and are publicly available as of the date of publication. The mass spectrometry proteomics data have been deposited to the ProteomeXchange Consortium via the PRIDE [21] partner repository and are publicly available as of the date of publication. Accession numbers are listed in the key resources table.

This paper does not report original code.

Any additional information required to reanalyze the data reported in this study is available from the lead contact upon request.

### EXPERIMENTAL MODEL AND STUDY PARTICIPANT DETAILS

#### Mice

Animal experimentation was carried out according to animal welfare regulation and guidelines of the French Ministry of Agriculture and Ministry of Higher Education, Research, and Innovation. The original mouse lines ($Taf7^{f}$, $Taf10^{f}$, $R26^{CreERT2}$, Tg(*T-Cre*)) were already described (Supplementary Table 3).

#### Generation and maintenance of mESCs

E3.5 blastocysts were collected from $R26^{CreERT2/+};Taf7^{f/f}$ X $Taf7^{f/f}$ or $R26^{CreERT2/+};Taf10^{f/f}$ X $Taf10^{f/f}$ mating. Uteruses were collected, embryos were flushed out with M2 medium (37°C) and placed in 96 well plates coated with mouse embryonic fibroblast (MEF) feeders in 2i+lif medium (DMEM medium supplemented with 15% fetal calf serum ES-tested, 2 mM L-glutamine, 0.1% ß-mercaptoethanol, 100 UI/ ml penicillin and 100 mg/ml streptomycin, 0.1 mM non-essential amino acids, 100 µL/50 mL of leukemia inhibitory factor (LIF), 3 µM CHIR99021 and 1 mM PD0325901) at 37°C under 5% $CO_2$. For initial amplification, mESCs were maintained on feeders until frozen in DMEM medium supplemented with 30% fetal calf serum and 20% DMSO. For experiment, mESCs were grow on gelatin. All the clones established were mycoplasma-free and were used for experiments before passage 35.

$R26^{CreERT2/+};Taf7^{f/f};Supt7l^{-/-}$ mESCs were generated from the clone *RT7#13*. These cells were transfected at 70% confluence with plasmid constructs containing *Cas9-EGFP* and gRNA (Supplementary Table

4) using Lipofectamine 2000 kit. Two days after, single EGFP$^{+}$ cells were isolated in 96 well plates using the BD FACS Aria TM II (BD Biosciences), amplified and frozen.

### METHOD DETAILS

#### Collection of mouse embryos

Embryos were collected in PBS, fixed in 4% PFA/PBS for 1 hour at 4°C under agitation and rinsed three times in PBS. The embryos were imaged using a Leica MZ16 macroscope coupled to a CoolSnap-Pro color camera (RS Photometrics).

#### Embryo sectioning and immunolocalization

Fixed embryos were equilibrated in 30% sucrose/PBS (3 h, 4°C), embedded in Cryomatrix (Thermo Fischer) and stored at -80°C. Fifteen micrometers-sections were cut on a Leica cryostat.

Sections were rehydrated in PBS, permeabilized in 0.5% Triton X-100/PBS (Sigma-Aldrich) for 40 min at RT (room temperature), blocked in AB buffer (3% Bovine Serum Albumin (BSA), 1% goat serum, 0.1% Tween 20 in PBS) for 40 min at RT and rewashed in 0.1% Triton X-100/PBS. Primary antibodies (Supplementary Table 2) were diluted 1/1000 in AB buffer and incubated overnight at 4°C. Sections were then washed three times in 0.1% Triton X-100/PBS for 40 min each. Secondary antibody (Supplementary Table 2) was diluted 1/1000 in 1 µg/mL DAPI (4′,6-diamidino-2-phenylindole dihydrochloride)/AB buffer and incubated for 1 h at RT. The sections were then washed in 0.1%Triton X-100/PBS several times. The slides were mounted in Vectashield® and imaged with a LSM 510 laser-scanning microscope (20x Plan APO objective). The pictures are shown with the LUT “Green Fire Blue” scale.

#### Genotyping and screening

Mouse tail tips were digested in 300 µL of 200 µg/mL of proteinase K in tail digestion buffer (10 mM Tris pH 7, 200 mM NaCl, 5 µM EDTA, 0.2% SDS). Embryonic yolk sacs and mESCs were digested in 100 µL of 200 µg/mL of proteinase K in 1X PCR buffer. 0.6 µL of lysate was used in 25 µL of PCR reaction with Taq DNA polymerase kit. Primers are listed in Supplementary Table 1.

#### 4-hydroxytamoxifen (4-OHT) treatment

$5x10^4$ or $8x10^5$ cells were seeded in 6 well plates or in one P100 petri dish, respectively, at day -1 (D-1). At D0, cells were treated with 100 nM of 4-OHT in a final volume of 2 mL or 12 mL, respectively. Control cells were treated with ethanol (0.1% EtOH final) in the same final volume. The experiments were performed at D2, D4 or D6. The 6 well plates were used for phenotypic analyses, and the P100 petri dishes for proteomic and transcriptomic analyses.

**mESC phenotypic analysis**

Cell counting

Trypsinized cells were resuspended in a dozen to hundreds of µL of PBS according to the size of the cell pellet. Cells were stained with Trypan Blue and counted on a Countess II Automated Cell Counter (Thermo Fischer Scientific). Different clones were used: *RT7*: 5 clones (#5, #7, #8, #13, #15), *RT10*: 5 clones (#3, #6, #9, #15, #41), *RT7; Supt7l*$^{-/-}$: 3 clones (#45, #71, #72).

Apoptosis assay

Dead floating and attached cells were collected and stained with the FITC Annexin V Apoptosis Detection Kit and counterstained with propidium iodide PI (Thermo Fischer Scientific) and analyzed using BD FACS Celesta (BD Bioscience). A minimum of 10,000 events were recorded. Different independent clones were used: *RT7*: 4 clones (#5, #7, #8, #13), *RT10*: 6 clones (#3, #6, #9, #15, #19, #41), *RT7; Supt7l*$^{-/-}$: 3 clones (#3, #71, #72).

Measuring of surface area of colonies

Cells were rinsed with PBS, fixed with 4% PFA for 30 min at 4°C, washed with PBS, stained for 30 min with 0.1% crystal violet dye and rewashed with PBS. Pictures were taken with a macroscope M420 (Leica) coupled with a CoolSNAP camera (RS Photometrics). Different independent clones were used: *RT7*: 2 clones (#5, #13), *RT10*: 3 clones (#3, #6, #9).

Cell cycle analysis by EdU

Cells were plated on gelatinized round glass slides. They were incubated with 10 µM of 5-ethynyl-2'-deoxyuridine (EdU, ThermoFischer Scientific) for 3 h and then fixed with 4% PFA for 15 min at RT. They were then stained with the EdU Staining Proliferation Kit (Abcam, ab222421). Slides were mounted, cells were imaged and EdU signal quantified as described in "Embryo sectioning and immunolocalization" and "Software and data analysis" sections.

**Depletion analysis**

Whole cell extract

Cells were pelleted, rinsed, resuspended in one equivalent volume of WCE buffer (50 mM Tris HCl pH7.9, 25% Glycerol, 0.2mM EDTA, 5 mM $MgCl_2$, 600 mM KCl, 0.5% NP40, 1 mM DTT, 1X cOmplete) and incubated for 30 min on ice. Then, 3 volumes of IP0 (25mM Tris HCl pH7.9, 5% Glycerol, 0.1% NP40, 1 mM DTT, 1X PIC) were added and incubated for 30 min on ice. Proteins presented in the supernatant were recovered after high-speed centrifugation.

Cytoplasmic, nuclear, chromatin extract

This method was previously published in [22]. One volume of cell pellet is lysed in 2 volumes of ice-cold E1 buffer (50 mM HEPES-KOH, pH7.5, 140 mM NaCl, 1 mM EDTA, pH8, 10% glycerol, 0.5% NP40, 0.25% Triton-X-100, 1 mM DTT, 1X cOmplete protease inhibitor (PIC)) by up-and-down pipetting. The suspension is centrifugated at 1100g at 4°C for 2min. The supernatant is collected as the cytoplasmic extract (CE). The pellet is resuspended and incubated 10 min on 5 volumes of ice-cold E1 buffer. After 2min of centrifugation at 1100g, the pellet is resuspended in 1 volume of ice-cold E2 buffer (10 mM Tris-HCl pH8.0, 200 mM NaCl, 1 mM EDTA pH8.0, 1 mM EGTA pH8.0, 1xPIC). The suspension is centrifugated at 1100g at 4°C for 2min. The supernatant is collected as the nuclear extract (NE). The pellet is resuspended and incubated 10 min on 2 volumes of ice-cold E2 buffer. After 2min of centrifugation at 1100g, the pellet is resuspended in 1 volume of ice-cold E3 buffer (50 mM Tris-HCl pH6.8, 20 mM NaCl, 1 mM MgCl2, 1% NP-40, 1xPIC) and transferred in a new clean tube. 1/1000 of benzonase (Sigma, E1014) is added and chromatin digestion is achieved during 30min at RT. The suspension is centrifugated at 16000g at 4°C for 10min. The supernatant is collected as the chromatin extract (ChrE).

Western blot (WB)

Protein concentrations were measured by Bradford method. 20 µg of WCE or 15µg of CE, NE or ChrE were boiled for 5 min in 100 mM Tris HCl pH 6.8, 30% glycerol, 4% SDS, 0.2% Bromophenol Blue, 100 mM DTT, resolved in 10% SDS-polyacrylamide gel and transferred to a nitrocellulose membrane (Protran, Amersham). After blocking in 3% milk/PBS, primary antibodies diluted 1/1000 in 0.3% milk/PBS (Supplementary Table 2) were incubated overnight at 4°C. After 3 washes in 0.05% Tween 20/PBS, HRP-coupled secondary antibodies (Supplementary Table 2) diluted 1/10000 in 0.3% milk/PBS were incubated 2 h at RT followed by ECL detection (ThermoFisher Scientific) in a ChemiDoc Touch Imaging System.

Immunofluorescence (IF) on mESC

Cells plated on gelatinized round glass slides were rinsed, fixed with 4% PFA/PBS for 10 min at RT, rinsed again and then permeabilized in 0.1% Triton X-100/PBS for 20 min at RT. After PBS washes, primary antibodies (Supplementary Table 2) diluted in 10% fetal calf serum (FCS)/PBS were incubated for 1 h at RT. Cells were rinsed twice with 0.02% Triton X-100/PBS for 5 min. Secondary antibodies (Supplementary Table 2) were diluted 1/1000 in 1 µg/mL DAPI/10% FCS/PBS and incubated for 1 h at RT. Cells were then washed twice for 5 min in 0.02% Triton X-100/PBS. Slides were mounted, cells were imaged and EdU signal quantified as described in “Embryo sectioning and immunolocalization” and “Software and data analysis” sections.

RT-qPCR

RNAs were extracted using TRI® Reagent (Molecular Research Center Inc), precipitated in isopropanol, washed with 75% EtOH, resuspended in RNase-free water and quantified with a Nanodrop (ThermoFisher Scientific). For RNA extraction from testis, tissues were stocked with a B pestle in a glass dounce grinder (Kimble) followed by high-speed centrifugation.

Reverse Transcription (RT) was performed using with 1 µg of total RNA using QuantiTect Reverse Transcription Kit (Qiagen) in T100 Bio-Rad machine. For qPCR, cDNAs were diluted 5 times and amplified using LightCycler 480 SYBR Green 2x PCR Master Mix I with 0.6 mM of forward and reverse primers (Sigma Aldrich, Sup. Table S1) in 8 µL of reaction volume. qPCR reaction was realized using a LightCycler 480 machine (Roche). Normalized values correspond to ΔCT.

**TFIID complex composition analysis**

Whole cell extracts

Cells (RT7#13 or RT10#41) were pelleted, rinsed, and resuspended in one volume of ice-cold Hypotonic Buffer (1 mL for 1 g of cells) (10 mM Tris pH8, 1.5 mM $MgCl_2$, 10 mM KCl supplemented with cOmplete protease inhibitor mix 1x (Roche). Cells were then lysed by 10 gentle strokes with a B pestle in a glass Dounce grinder (Kimble). After 10 min of centrifugation at 9000g, the pellet, which contains nuclei, was resuspended in one volume of High Salt Buffer (20 mM Tris pH8, 1.5 mM $MgCl_2$, 450 mM NaCl, 0.2 mM EDTA, 25% glycerol, 0.5% NP40, supplemented with cOmplete protease inhibitor mix 1x). Nuclei were lysed by 10 gentle strokes and incubated on ice for 30 min. After 10 min of centrifugation at 9000 g, the supernatant was recovered as nuclear enriched whole cell extract (NWCE).

IP-MS analysis

*Immunoprecipitation (IP)*

One mg of NWCE was first incubated for 1 h with 120 µL of Protein-G Sepharose beads (GE healthcare) in 1 mL of IP100 buffer (25 mM Tris HCl pH 7.9, 10% Glycerol, 0.1% NP40, 5 mM $MgCl_2$, 100 mM KCl, 1X cOmplete) at 4°C under gentle agitation. NWCE was isolated and incubated with 10 to 30 µL of antibodies (Supplementary Table 2) during 2 hours at 4°C under gentle agitation and then incubated with fresh 120µL of Protein-G Sepharose beads overnight. Beads were then washed at 4°C twice with IP500 Buffer (25 mM Tris HCl pH7.9, 10% Glycerol, 0.1% NP40, 5 mM $MgCl_2$, 500 mM KCl, 1X cOmplete) under gentle agitation and then three times with IP100 buffer, each time for 10min. Immunoprecipitated proteins were eluted with 50 µL of acid Glycine buffer (0.1 M glycine pH2.8) directly buffered with 1 µL of 10 mM Tris HCl pH8. Each immunoprecipitation was verified by western blot by loading 20 µL of input and of supernatant and 15 µL of eluted proteins.

*Liquid digestion*

Eluted proteins were TCA-precipitated overnight at 4°C. Samples were then centrifuged at 14000 rpm for 30 minutes at 4°C. Pellets were washed twice with 500 µL cold acetone, centrifuged at 14000 rpm for 10 minutes at 4°C, denatured with 8 M urea in Tris-HCl 0.1 mM, reduced with 5 mM TCEP for 30 minutes, then alkylated with 10 mM iodoacetamide for 30 minutes in the dark. Both reduction and alkylation were performed at room temperature and under agitation (850 rpm). Double digestion was performed with endoproteinase Lys-C (Wako) at a ratio of 1/100 (enzyme/proteins) in 8 M urea for 4h, followed by an overnight modified trypsin digestion (Promega) at a ratio of 1/100 (enzyme/proteins) in 2 M urea. Both LysC and Trypsin digestions were performed at 37°C. Peptide mixtures were then desalted on C18 spin-columns and dried on Speed-Vacuum before LC-MS/MS analysis.

*LC-MS/MS analysis*

Each sample was analyzed in triplicate (experimental triplicate) using an Ultimate 3000 nano-RSLC (Thermo Scientific, San Jose California) coupled in line with a LTQ-Orbitrap ELITE mass spectrometer via a nano-electrospray ionization source (Thermo Scientific, San Jose California).

Peptide mixtures were loaded on a C18 Acclaim PepMap100 trap-column (75 µm ID x 2 cm, 3 µm, 100Å, Thermo Fisher Scientific) for 3.5 minutes at 5 µL/min with 2% ACN, 0.1% FA in $H_2O$ and then separated on a C18 Accucore nano-column (75 µm ID x 50 cm, 2.6 µm, 150Å, Thermo Fisher Scientific) with a 100 minutes linear gradient from 5% to 50% buffer B (A: 0.1% FA in $H_2O$ / B: 99% ACN, 0.1% FA in $H_2O$), then a 20 minutes linear gradient from 50% to 70% buffer B, followed with 10 min at 99% B and 10 minutes of regeneration at 5% B. The total duration was set to 140 minutes at a flow rate of 200nL/min. The oven temperature was kept constant at 40°C.

The mass spectrometer was operated in positive ionization mode, in data-dependent mode with survey scans from m/z 300-1600 acquired in the Orbitrap at a resolution of 240,000 at m/z 400. The 20 most intense peaks (TOP20) from survey scans were selected for further fragmentation in the Linear Ion Trap with an isolation window of 2.0 Da and were fragmented by CID with normalized collision energy of 35%. Unassigned and single charged states were rejected.

The Ion Target Value for the survey scans (in the Orbitrap) and the MS2 mode (in the Linear Ion Trap) were set to 1E6 and 5E3 respectively and the maximum injection time was set to 100 ms for both scan modes. Dynamic exclusion was used. Exclusion duration was set to 30 s, repeat count was set to 1 and exclusion mass width was ± 10 ppm.

*Data Analysis*

Proteins were identified by database searching using SequestHT (Thermo Fisher Scientific) with Proteome Discoverer 2.2 software (Thermo Fisher Scientific) on *Mus musculus* database (Swissprot, non-reviewed, release 2019_08_07, 55121 entries). Precursor and fragment mass tolerances were set at 7 ppm and 0.5 Da respectively, and up to 2 missed cleavages were allowed. Oxidation (M) was set as variable modification, and Carbamidomethylation (C) as fixed modification. Peptides were filtered with a false discovery rate (FDR) at 1%, rank 1 and proteins were identified with 1 unique peptide.

Each experiment was analyzed separately using Extracted Ion Chromatogram (XIC) values [23]. First, only peptides whose mean XIC values from triplicate measurements of the control condition (EtOH) were greater than the mean XIC values from triplicate measurements of the mock IP were kept for further analysis. Note that TAF4 and TAF4B were analyzed together in a virtual TAF4.4B protein and TAF9 and TAF9B in a TAF9.9B protein. Second, protein XIC values (PXV) were calculated by averaging XIC values of peptides kept belonging to the same proteins (1). Third, PXV were normalized by the mean PXV of the mock IP ($\Delta PXV_x$) (2) and then by the normalized mean PXV of the bait ($\Delta PXV_{bait(x)}$) (3). Last, fold change (FC) was calculated from one condition (EtOH or 4-OHT) and the control condition (EtOH) (4).

(1) $PXV_x = \frac{\sum_{i=1}^{n} XIC_{peptide_{(x)}}}{n}$

(2) $\Delta PXV_x = PXV_{x(IP)} - \frac{\sum_{j=1}^{n} PXV_{x_{(IPmock)}}}{n}$

(3) $\Delta PXV_{bait(x)} = \Delta PXV_x \;/\; \frac{\sum_{j=1}^{n} \Delta PXV_{bait}}{n}$

(4) $\mathrm{FC} = \frac{\Delta PXV_{bait(x)}}{\Delta PXV_{EtOH(bait(x))}}$

where x is the protein of interest, i is the number of peptides belonging to the same protein and j is the number of measurements performed on the same IP under the same conditions.

In the graph, each point corresponds to one FC value and the bar plot is the average of the FC values from different IP experiments, corresponding to the same condition, for a protein or group of proteins belonging to the same sub-complex.

<u>Gel Filtration (GF)</u>

One mg of NWCE was diluted twice in the GF buffer (20 mM Tris pH8, 1.5 mM $MgCl_2$, 450 mM NaCl, 0.2 mM EDTA,10% glycerol, 0.5% NP40), centrifuged for 10min at 16 000 rpm then passed at 0.4 mL/min through a Superose 6 GL 10/300 column (Sigma Aldrich) previously equilibrated with GF buffer. About 60 fractions of 250 µL were collected and 27µL of each fraction were analyzed by western blot.

**Transcription analysis**

<u>Analysis of newly synthesized RNA by EU labelling</u>

Cells (RT7#13 or RT10#41) plated on gelatinized round glass slides were incubated with 1 mM of 5-ethynyl-uridine (EU, ThermoFisher Scientific, E10345) for 1 h, fixed with 4% PFA for 15 min at RT and rinsed. They were then stained using the Click-it RNA Imaging Kits (Invitrogen, C10329) according to the manufacturer's guidelines. As a control, cells were incubated 1 h either with 500 µM of triptolide (Sigma Aldrich, T-3652) or 300 µM of flavopiridol (Sigma Aldrich, F-3055) followed by another hour incubation with 1 mM EU. Cells were imaged and EU signal quantified as described in IF section.

Newly synthesized 4-sU RNA sequencing

*4-sU RNA labeling and purification*

The protocol for newly synthesized RNA sequencing is based on published protocols [24]. Briefly, mESCs were seeded at D-1, treated with either EtOH (control) or 4-OHT (mutant) from D0 to D3 with a reseeding/amplification step at D2. On D3, cells were labelled with 500 µM 4-sU for 15 min. In parallel, drosophila S2 cells were also labelled with 500 µM 4-sU for 15 min. RNAs were extracted with Trizol, precipitated with isopropanol, washed with 75% ethanol, and resuspended in DEPC-treated water. DNAse treatment was performed using the TURBO DNA-free kit. Non labelled *S.cerevisiae* RNAs were isolated using the RiboPure - Yeast kit (Invitrogen). RNAs were measured with the Qbit machine using the Quant-it RNA Broad Range kit. 200 µg of mESC RNAs were mixed with 25 µg of S2 RNAs and 25 µg of yeast RNAs. The mixture was precipitated again, resuspended in 130 µL of DEPC-treated water and then fragmented with Covaris E220 sonicator. 1 µL of RNA were collected before and after sonication to check fragmentation on a 1% agarose gel and on bioanalyzer. The fragmented RNA is expected to have an average size of 1.5 kb. To isolate newly synthesized 4-sU labelled RNA fragments, RNAs were first biotinylated with Biotin-HPDP molecules, then combined with streptavidin magnetic beads and finally isolated on column (µMACS streptavidin beads and kit, miltenyi). RNAs from flowthrough and elution are then precipitated in absolute EtOH with 0.1 mg/mL glycogen and 300mM NaOAc (pH 5.2) overnight at -20°C, washed with 75% EtOH and resuspended in respectively 150µL and 15µL of DEPC-treated water.

The quality of 4-sU labelled RNA fragment purification was checked by RT-qPCR prior sequencing. 2 µL of eluted RNAs and 7.5 µL of RNAs from the flowthrough were collected into respectively 12 µL and 2.5 µL of DEPC-treated water. RT was carried out using Superscript IV kit and qPCR by 480 SYBR green I Master kit. Primers are listed in Supplementary Table 3. Good purification is characterized by a lower amount of yeast RNA and a higher amount of mouse intronic sequence in the purified sample than in the flowthrough sample after normalization to Drosophila RNAs (ΔCt) and RNA from EtOH flowthrough sample (-ΔΔCt).

*Library preparation*

15 to 50 ng of eluted RNA was used for the library preparation using TruSeq Stranded Total RNA LT Sample Prep Gold Kit (Illumina, RS-122-230) according to the Illumina protocol with the following modifications. Four-thiouridine-labelled RNA was cleaned up using 1.8× RNA Clean AMPure XP beads (A63882, Beckman-Coulter) and fragmented using divalent cations at 94ºC for 1 min without depletion of rRNA. While double stranded cDNA synthesis and adapter ligation were performed according to manufacturer instructions, the number of PCR cycles for library amplification was reduced to 10 cycles. After purification using SPRIselect beads (B23319, Beckman-Coulter), libraries were sequenced with 1×50 bp on a HiSeq4000 System (Illumina).

*Data analysis*

Reads were preprocessed to remove adapter, polyA, low-quality sequences (Phred quality score below 20) and reads shorter than 40 bases. These preprocessing steps were performed using cutadapt (version 1.10, [12]). Reads were mapped to rRNA sequences using bowtie (version 2.2.8, [11]) and reads mapping to rRNA sequences were then removed for further analysis. Reads were mapped onto the mm10 and BDGP6 assembly of *Mus musculus* and *Drosophilia melanogaster* genome using STAR (version 2.5.3a, [19]). Gene expression quantification was performed from uniquely aligned reads using htseq-count (version 0.6.1p1, [17]), with annotations from Ensembl version 93 and "union" mode. Only non-ambiguously assigned reads to a gene have been retained for further analyses. Data were normalized using size factors computed with the median-of-ratios method, proposed in [25], on *Drosophilia melanogaster* counts. Principal Component Analysis was computed on variance stabilizing transformed data calculated with the method proposed in [14], using size factors computed from *Drosophilia melanogaster* counts. Comparisons of interest were performed using Wald statistic test for differential expression and implemented in the Bioconductor package DESeq2 (version 1.16.1, [14]). Eulerr plots were generated using eulerr package (version 4.2.2). In genome browser view, normalized mESC read counts were used.

**Cleavage under targets and release using nuclease (CUT&RUN) and library preparation.**

CUT&RUN experiments were performed in biological duplicate as described in [26]. Briefly, mESC (RT7#13 or RT10#41) were treated with EtOH or 4-OHT for 3 days and 250,000 cells were used per CUT&RUN sample. Cells were washed and resuspended in wash buffer and incubated for 10 min at room temperature with 10 µl of concanavalin A-coated beads (Bangs Laboratories, BP531). Cells bound to beads were permeabilized using 0.05% digitonin and incubated with the appropriate antibody at 4°C overnight. Protein A-MNAse (pA-MN) was added to a final concentration of 700 ng/ml and incubated at 4°C for 1 h on a tube rotator. After washing, pA-MN was activated with 2 µl of 100 mM CaCl2 and digestion was performed for 30 min at 0°C. The reaction was stopped with 100 µl of stop buffer containing 2 pg/ml of heterologous spike-in DNA from yeast. Release of the DNA fragments was achieved by incubating samples at 37°C during 15 min. DNA was extracted using NucleoSpin columns (Macherey-Nagel) and eluted in 30 µl of NE buffer.

CUT&RUN-seq libraries were generated using the MicroPlex library preparation kit v3 (Diagenode), following the manufacturer's instructions, except that the stage 4 of the library amplification PCR was performed with a combined annealing-extension step for 10 s at 60°C, and stage 5 with an extension step for 10 s at 60°C for 7 cycles. Yield and size distribution were quantified on a 2100 Bioanalyzer instrument (Agilent). Sequencing was performed by the GenomEast platform (IGBMC) on an Illumina NextSeq 2000 (PE-50, 20 million reads).

**CUT&RUN data analysis**

Data were preprocessed with cutadapt v4.0 [12] to trim adapter sequences (Nextera Transposase Sequence) from 3' end of reads. Cutadapt was used with the following parameters '-a AGATCGGAAGAG -A AGATCGGAAGAG -m 25:25'. Reads were mapped to Mus musculus genome (assembly mm10) using Bowtie2 v2.4.4 [11] with default parameters except for "–end-to-end –very-sensitive –no-mixed –no-discordant -I 10 -X 700". The following table shows the number of reads aligned to the Mus musculus genome. BigWig files were generated using deeptools bamCoverage v3.5 [13] with the following parameters "-bs 10 –normalizeUsing CPM –effectiveGenomeSize 2652783500 –skipNonCoveredRegions –extendReads". Bigwig files of mean signal per condition were generated using deeptools bamCompare v3.5 [13] with the following parameters "-of bigwig –operation mean –effectiveGenomeSize 2652783500 –normalizeUsing CPM –scaleFactorsMethod None –extendReads". The peak calling was done with Macs2 v2.2.7.1 [18] with default parameters except "-f BAMPE -q 0.1". Peaks were annotated relative to genomic features using Homer v4.11 [16] (annotations got extracted from gtf file downloaded from Ensembl 102)). The tool deeptools computeMatrix v3.55 [13] was used to generate a count matrix at the positions of interest (union of peaks of all datasets to compare) and finally the tool deeptools plotProfile v3.5 was used to generate mean profile plots and deeptools plotHeatmap v3.5 was used to generate heatmaps. Top 10% peaks were filtered using an in-house R scripts, using the fold enrichment score as input.

**Software and data analysis**

Analyses were performed using custom scripts available on request (ImageJ version 1.53q, R software version 4.0.2, [15]). R analyses were achieved using ggplot2 version 3.4.0, tibble 3.1.8, tidyr 1.2.1, readr 2.1.3, purr 0.3.5, dplyr 1.1.10, stringr 1.4.1, forcats 0.5.2) and rstatix (version 4.2.2, https://rpkgs.datanovia.com/rstatix/) for statistical analysis. For specific analyses, such as RNA sequencing, the libraries used are indicated in the text of these specific sections.

Genome browser view of reads from 4-sU RNA sequencing and CUT & RUN experiment were created using UCSC genome browser (https://genome.ucsc.edu/).

Western blot images were processed on Bio Rad Image Lab Software (version 5.2.1).

**Supplementary Table 1: List of PCR and qPCR primers**

| Species | Gene | Forward | Reverse | Usage |
|---|---|---|---|---|
| Mouse | *Taf7* | GTATGAAAACCTGTGTCCTGG TCTG | GAAGGCAAGTTCTCAATGAA AGGG | PCR |
| Mouse | *Taf10* | GTAGTGTCCAGCACACCTCT | CAGTCTAACCTGCTCCGAGT | PCR |
| Mouse | *Cre* | TGATGGACATGTTCAGGGATC | CAGCCACCAGCTTGCATGA | PCR |
| Mouse | *Rosa-Wt* | AAAGTCGCTCTGAGTTGTTAT | CCTGATCCTGGCAATTTCG | PCR |
| Mouse | *Rosa-Cre* | GGAGCGGGAGAAATGGATATG | CCTGATCCTGGCAATTTCG | PCR |
| Mouse | *Supt7l-exon1-2* | ATTGTGGCGACTGCTTGATAG | ACCCAGAGAGTGACTTTTACCG | PCR |
| Mouse | *Supt7l-5'UTR* | GCAGTTCCCACATAAGAAGCA | AGCCGCGTATACCACTCCT | PCR |
| Mouse | *Rn7sk* | CCATTGTAGGAGAACGTAGGG TAGTCAAGC | CCACATGCAGCGCCTCATT | qPCR |
| Mouse | *Taf7* | AATATGCCGCTACGGTGAGG | TCAGGTTGACATGCCCAGAC | qPCR |
| Mouse | *Taf7L* | CCTGAGAAACATCCGCGGTC | AGACAACGTCTCACTGCCTG | qPCR |
| Mouse | *Taf7L2* | GCAACGGAACGTGTGAAGTG | CCGATGGGAAGTCGTTGTTG | qPCR |
| Mouse | *Taf10* | GAGGGGGCAATGTCTAACGG | CGCGGTTCAGGTAGTAACCA | qPCR |
| Mouse | *Tpt1 (intron)* | TTAAGCACATCCTTGCTAATTTCA | TGTACGAGACAGCAAACAGACTTT | qPCR |
| Mouse | *Clf1 (intron)* | TATGAGACCAAGGAGAGCAAGAA | GTTAAGCTCTGAGAAAGGGAACC | qPCR |
| Drosophila | *Rpl12* | AAGGGAACCTGCAAGGAAGT | CCCTCGTTCAGTTCGTCAATA | qPCR |
| Drosophile | *α-Tubuline* | GCTTCCTCATCTTCCACTCG | GCTTGGACTTCTTGCCGTAG | qPCR |
| *S.cerevisiae* | *PMA1* | CTCATCAGCCAACTCAAGAAA | CGTCATCGTCAGAAGATTCA | qPCR |
| *S.cerevisiae* | *PRPS3* | ATTGTTGAACGGTTTGGC | CCCTTAGCACCAGATTCCATA | qPCR |

**Supplementary Table 2: List of sgRNAs used to generate the *R26$^{CreERT2/+}$;Taf7$^{f/f}$;Sup7l$^{-/-}$* mouse ES cell lines (*RT7;Sup7l$^{-/-}$*)**

| Name | Sequence | PAM | Reference |
|---|---|---|---|
| *mSupt7l-3* | ACCATCTCCCTCGCCCCG | AGG | (Fischer et al., 2021) |
| *mSupt7l-4* | ACCAGTACGTATTCAGAG | TGG | |

1. Mohan, W.S., Scheer, E., Wendling, O., Metzger, D., and Tora, L. (2003). TAF10 (TAF(II)30) is necessary for TFIID stability and early embryogenesis in mice. Mol Cell Biol *23*, 4307–4318. 10.1128/mcb.23.12.4307-4318.2003.

2. Jacq, X., Brou, C., Lutz, Y., Davidson, I., Chambon, P., and Tora, L. (1994). Human TAFII30 is present in a distinct TFIID complex and is required for transcriptional activation by the estrogen receptor. Cell *79*, 107–117.

3. Langer, D., Martianov, I., Alpern, D., Rhinn, M., Keime, C., Dollé, P., Mengus, G., and Davidson, I. (2016). Essential role of the TFIID subunit TAF4 in murine embryogenesis and embryonic stem cell differentiation. Nature communications *7*, 11063–16. 10.1038/ncomms11063.

4. Bardot, P., Vincent, S.D., Fournier, M., Hubaud, A., Joint, M., Tora, L., and Pourquié, O. (2017). The TAF10-containing TFIID and SAGA transcriptional complexes are dispensable for early somitogenesis in the mouse embryo. Development *144*, 3808–3818. 10.1242/dev.146902.

5. Martianov, I., Velt, A., Davidson, G., Choukrallah, M.-A., and Davidson, I. (2016). TRF2 is recruited to the pre-initiation complex as a testis-specific subunit of TFIIA/ALF to promote haploid cell gene expression. Scientific reports *6*, 32069. 10.1038/srep32069.

13. Ramírez, F., Ryan, D.P., Grüning, B., Bhardwaj, V., Kilpert, F., Richter, A.S., Heyne, S., Dündar, F., and Manke, T. (2016). deepTools2: a next generation web server for deep-sequencing data analysis. Nucleic Acids Res. *44*, W160–W165. 10.1093/nar/gkw257.

22. Gillotin, S. (2018). Isolation of Chromatin-bound Proteins from Subcellular Fractions for Biochemical Analysis. BIO-Protoc. *8*, e3035. 10.21769/bioprotoc.3035.

23. Smith, R., and Tostengard, A.R. (2020). Quantitative Evaluation of Ion Chromatogram Extraction Algorithms. J. Proteome Res. *19*, 1953–1964. 10.1021/acs.jproteome.9b00768.

24. Fischer, V., Plassard, D., Ye, T., Reina-San-Martin, B., Stierle, M., Tora, L., and Devys, D. (2021). The related coactivator complexes SAGA and ATAC control embryonic stem cell self-renewal through acetyltransferase-independent mechanisms. Cell Reports *36*, 109598. 10.1016/j.celrep.2021.109598.

25. Anders, S., and Huber, W. (2010). Differential expression analysis for sequence count data. Genome biology *11*, R106. 10.1186/gb-2010-11-10-r106.

26. Meers, M.P., Bryson, T., and Henikoff, S. (2019). A streamlined protocol and analysis pipeline for CUT&RUN chromatin profiling. bioRxiv, 569129. 10.1101/569129.
