## Supplementary material for "RNA polymerase II transcription with partially assembled TFIID complexes": figures_plus_legend

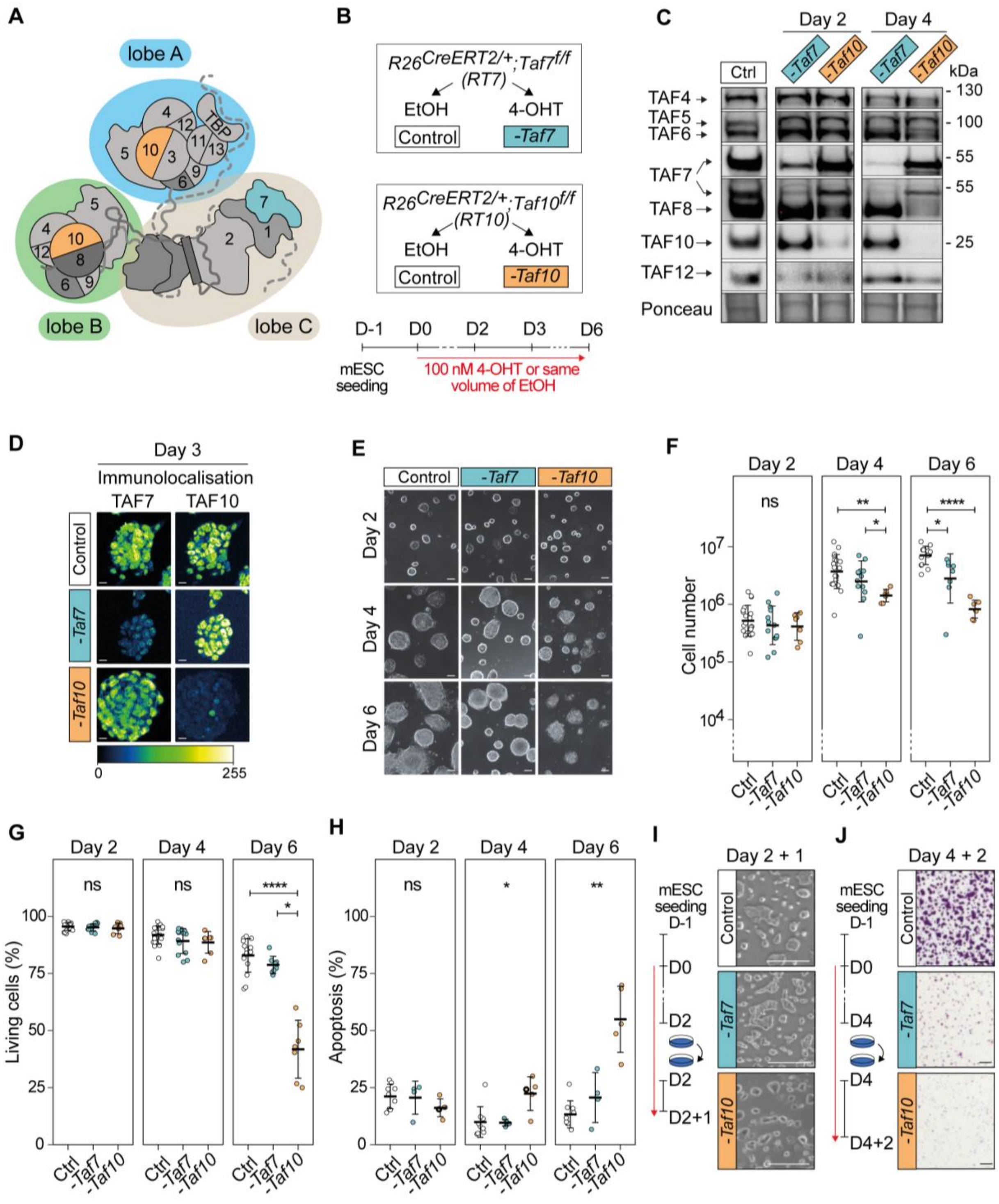


**Fig 1: Phenotypic analysis of the conditional depletion of TAF7 or TAF10 in mESCs.**

(A) Schematic structure of TFIID. Lobe A is indicated in blue, lobe B in green and lobe C in beige. (B) Strategy of the induction of the deletion of *Taf7* (-*Taf7*) in *R26*$^{CreERT2/+}$;*Taf7*$^{f/f}$ (*RT7*) and *Taf10* (-*Taf10*) in *R26*$^{CreERT2/+}$;*Taf10*$^{f/f}$ (*RT10*) mESCs. Cells are plated at day (D) -1, 100 nM 4-hydroxytamoxifen (4-OHT, depleted) or the same of volume of ethanol (control) were added at D0 and maintained until the day of the analysis. (C) Western blot analyses of TAF4, TAF5, TAF6, TAF7, TAF8, TAF10 and TAF12 proteins expression after *Taf7* (-*Taf7*) or *Taf10* (-*Taf10*) deletions after 2 and 4 days of 4-OHT treatment. As a control (Ctrl), *RT7* cells were treated 2 days with EtOH. The Ponceau staining is displayed at the bottom of the panel. (D) Immunolocalization of TAF7 and TAF10 in *RT7* and *RT10* cells treated for 3 days with 4-OHT. As a control, *RT10* cells were treated with EtOH for 3 days. Color scale (Green Fire Blue LUT scale) is indicated at the bottom. Scale bar; 15µm. (E) Monitoring of cell growth over time. *RT7* and *RT10* cells were treated over 6 days, and pictures were taken at D2, D4 and D6. Scale bar; 50 µm. (F) Log10 of the total number of cells at D2, D4 and D6 of treatment. (G) Percentage of living cells after 2, 4, and 6 days of treatment determined by Trypan blue staining. For (E, F), Ctrl: D2; N = 20, D4; n = 20, D6; n = 15, -*Taf7*: D2; n = 13, D4; n = 13, D6; n = 8, -*Taf10*: D2; n = 7, D4; n = 7, D6; n = 7 biological replicates. (H) Percentage of apoptotic cells after 2, 4 and 6 days of treatment determined by Annexin V and propidium iodide (PI) staining. For D2, D4 and D6, Ctrl; n = 9, -*Taf7*; n = 4, -*Taf10*; n = 5 biological replicates. The bars correspond to the mean ± standard deviation. Kruskal-Wallis test followed by Dunn post hoc test if significant: ns; not significant, * <0.05; **<0.01; *** <0.001, **** <0.0001. (I) Cell density after passage at D2 and 1 day of extra culture. (J) Cell density evaluated by Crystal violet staining after passage at D4 and 2 days of culture. Scale bar; 150 µm. For D, F and G, the control conditions correspond to *RT7* cells treated with EtOH.

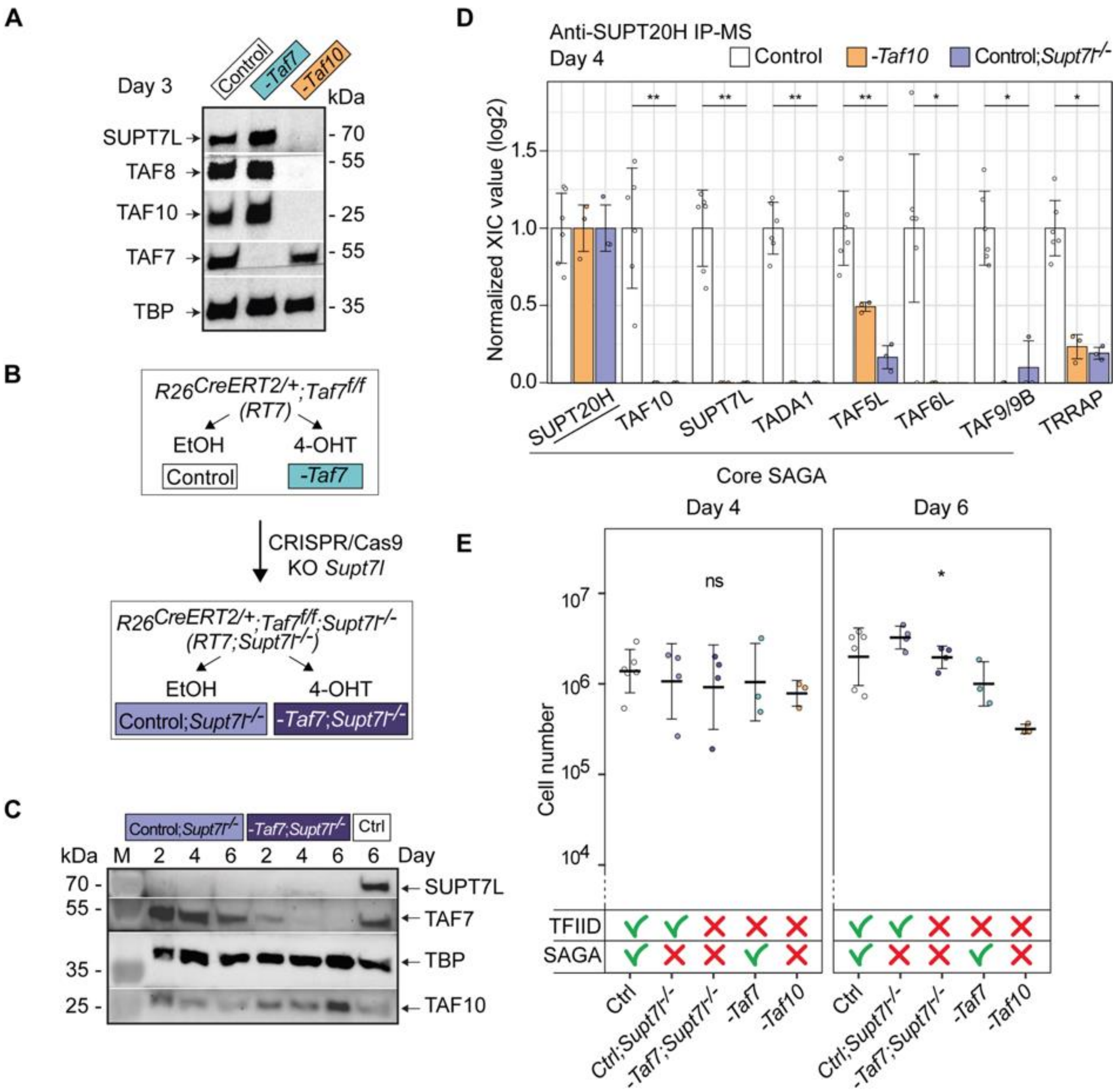


**Fig 2: The more severe phenotype in TAF10 depleted mESCs is not due to the SAGA assembly defect.** **(A)** Western blot analyses of SUPT7L, TAF8, TAF10, TAF7 and TBP expression after deletion of *Taf7* (-*Taf7*) or of *Taf10* (-*Taf10*). *RT7* and *RT10* cells were treated 3 days with 4-OHT and as control, *RT7* cells were treated 3 days with EtOH. **(B)** CRISPR/Cas9 strategy to generate the $R26^{CreERT2/+}$;$Taf7^{f/f}$;$Supt7l^{-/-}$ (*RT7*;$Supt7l^{-/-}$) mESCs. The second coding exon (exon 3) of *Supt7l* was deleted in *RT7* mESCs. Deletion of *Taf7* in the $Supt7l^{-/-}$ genetic background is induced by 4-OHT treatment. **(C)** Western blot analyses of SUPT7L, TAF7, TBP and TAF10 expression in *RT7*;$Supt7l^{-/-}$ cells after 2, 4 and 6 days of treatment with EtOH (Control; $Supt7l^{-/-}$) or 4-OHT (-*Taf7*; $Supt7l^{-/-}$). As a wild type control, *RT7* cells were treated 6 days with EtOH. M; molecular weight marker. **(D)** Anti-SUPT20H IP-MS analyses on nuclear enriched lysates from *RT10* and *RT10* cells treated 4 days with EtOH (Control, *RT7* and *RT10* data merged), *RT10* mESCs treated 4 days with 4-OHT (-*Taf10*) and *RT7*;$Supt7l^{-/-}$ cells treated 4 days with EtOH (Control;$Supt7l^{-/-}$). For each of the proteins of interest, the XIC values of Control;$Supt7l^{-/-}$ and -*Taf10* lysates were normalized to those of *RT7* and *RT10* control cells treated with EtOH, respectively. Control; n = 2 biological replicates x 3 technical replicates, -*Taf10*; n = 1 x 3, Control;$Supt7l^{-/-}$; n = 1 x 3. Means ± standard deviation are shown. Kruskal-Wallis test: * <0.05; **<0.01. **(E)** Total number of cells after 4 and 6 days of treatment. The control condition corresponds to the merging of *RT7* and *RT10* cells treated with EtOH. The impact on TFIID and SAGA assembly is indicated at the bottom. D4, D6: Ctrl; n = 6, Ctrl;$Supt7l^{-/-}$; n = 4, -*Taf7*;$Supt7l^{-/-}$; n = 4, -*Taf7;* n = 3, -*Taf10*; n = 3 biological replicates. Means ± standard deviation are shown. Kruskal-Wallis test followed by a Dunn post hoc test: ns; not significant, * <0.05.

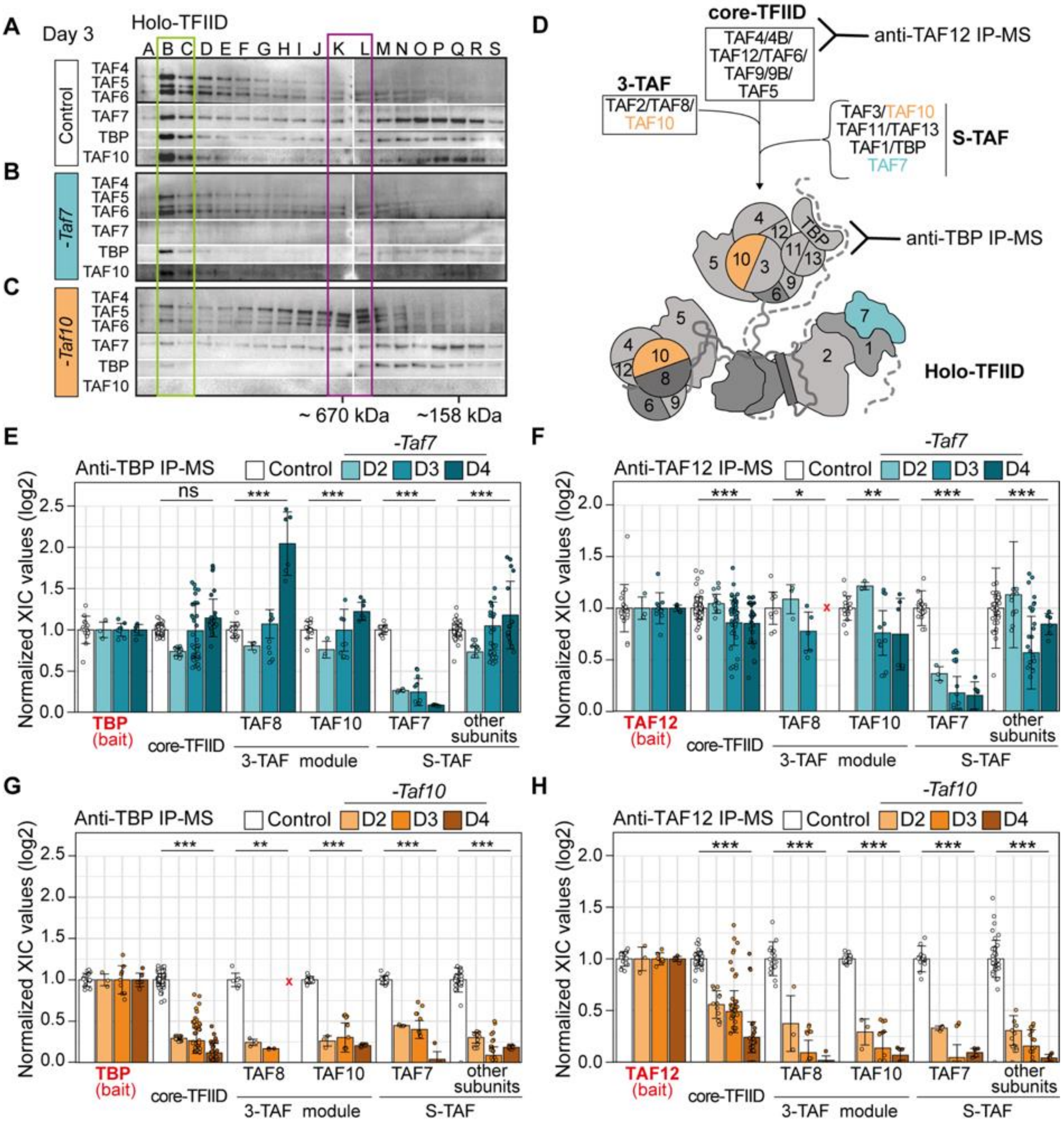

**Fig 3: Depletion of TAF7 or TAF10 differentially affects TFIID assembly. (A-C)** Gel filtration coupled to western blot analysis of TAF4, TAF5, TAF6, TAF7, TAF10 and TBP expression in control *RT10* mESCs treated with EtOH (Control) (A), *RT7* (*-Taf7*) and *RT10* (*-Taf10*) depleted mESCs treated with 4-OHT (B,C) for 3 days. The letters on the top correspond of the different fractions analyzed. Molecular weight positions are indicated at the bottom. Positions of the different complexes are indicated by boxes (n=1). **(D)** Assembly of the holo-TFIID complex representation. Anti-TBP antibodies will detect the presence of the holo-TFIID only while anti-TAF12 antibodies will detect all intermediates from core-TFIID to holo-TFIID. **(E-H)** Relative normalized XIC values of TFIID subunits from anti-TBP IP-MS (E,F) and anti-TAF12 IP-MS (G,H) from *RT7* (E,G) and *RT10* mESCs (F,H) at day 2, 3 and 4 of EtOH (Control) or 4-OHT (*-Taf7* and *-Taf10*) treatment. Subunits of the same submodules (see D) were merged together except for the bait proteins TBP (E,F) and TAF12 (G,H), and TAF7, TAF8 and TAF10. Note that TAF2 data were not taken into account because TAF2 was poorly detected in the controls in the 2 IP-MS. Core-TFIID corresponds to TAF4, TAF5, TAF6, TAF9/9B and TAF12, S-TAF corresponds to TAF1, TAF3, TAF11, TAF13 and TBP for. D2; n = 1 biological replicate x 3 technical replicates, D3; n = 3 biological replicates x 3 technical replicates, D4; n = 2 biological replicates x 3 technical replicates. Red crosses indicate proteins not detected in the control condition. Each dot corresponds to one measure of one subunit. Means ± standard deviation are shown, Kruskal-Wallis test: ns; not significant, * <0.05; **<0.01; *** <0.001.

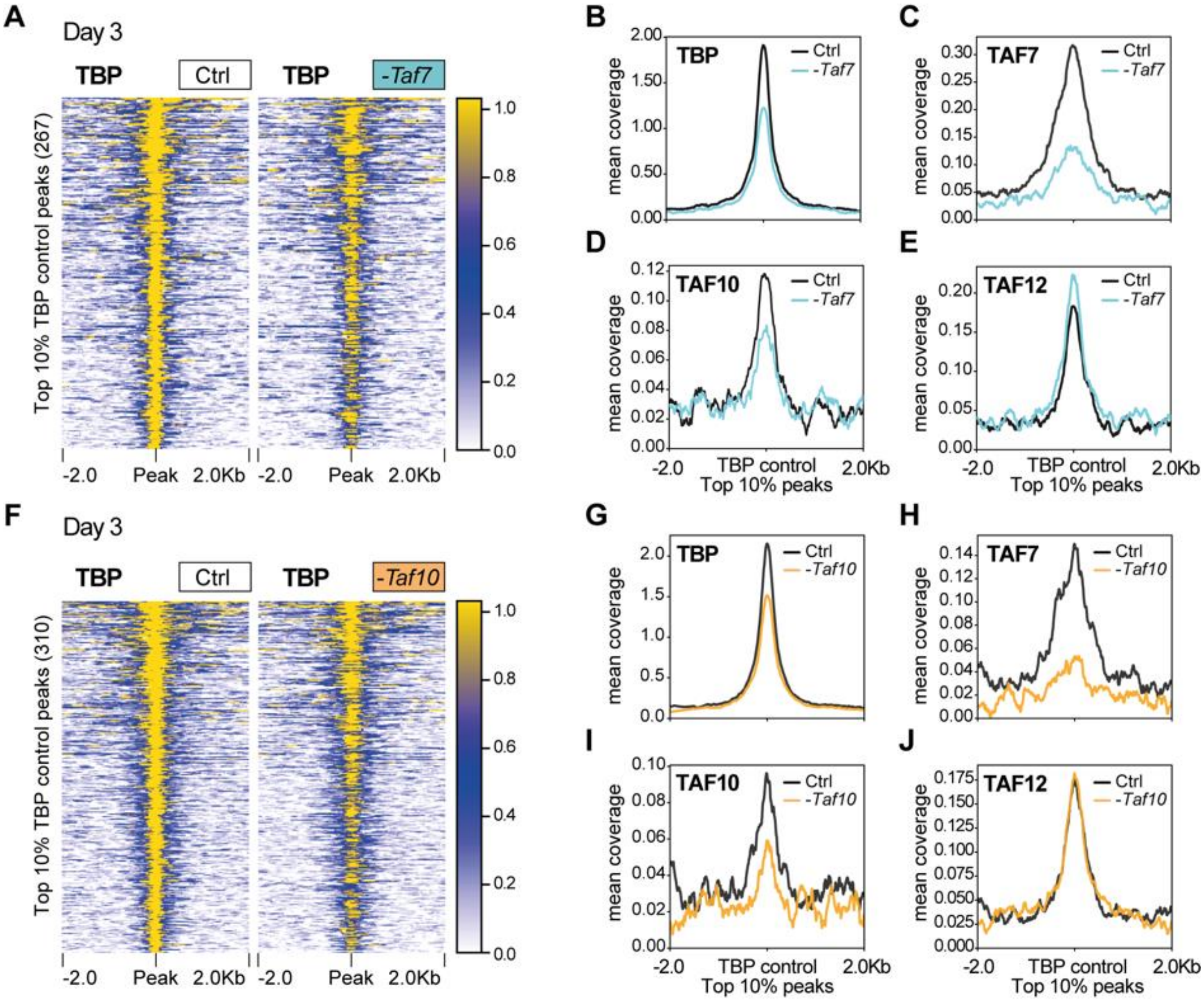


**Figure 4: TBP is still recruited at promoters in *TAF7* or *TAF10*-depleted mESCs. (A)** Heatmap showing the TBP distribution at the position of the top 10% (fold enrichment)TBP CUT&RUN peaks (± 2 kb) detected in the control condition, in *RT7* control (Ctrl) and TAF7-depleted (-*Taf7*) mESCs after 3 days of treatment. **(B-E)** Mean profile of TBP (B), TAF7 (C), TAF10 (D) and TAF12 (E) protein accumulation at the location of the TBP control peaks, in control (black line) and in TAF7 depleted (blue line) mESCs. **(F)** Heatmap showing the TBP distribution at the position of the top 10% TBP CUT&RUN peaks (± 2 kb) detected in the control condition, in *RT10* control (Ctrl) and TAF10-depleted (-*Taf10*) mESCs after 3 days of treatment. **(G-J)** Mean profile of TBP (G), TAF7 (H), TAF10 (I) and TAF12 (J) protein accumulation at the location of the TBP control peaks, in control (black line) and in TAF10 depleted (orange line) mESCs.

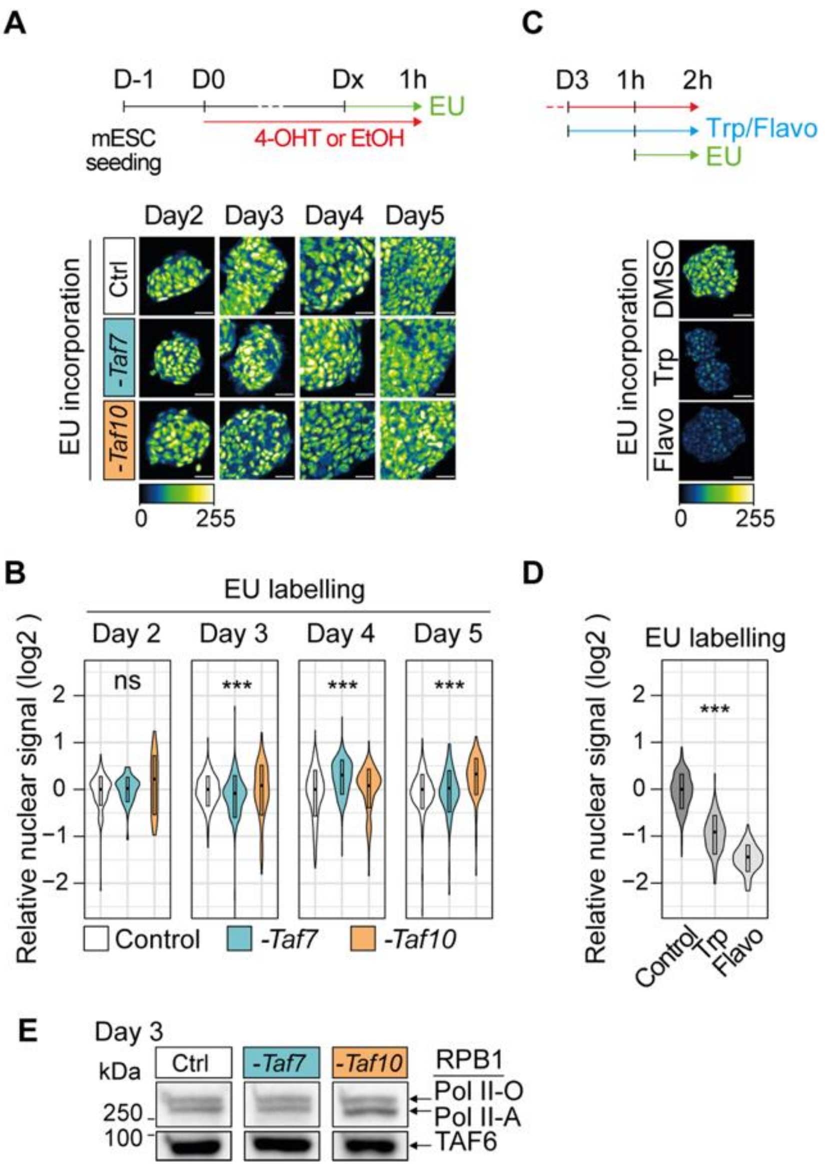


**Figure 5: Depletion of holo-TFIID in absence of TAF7 or TAF10 has only limited defects on Pol II global transcription. (A,B)** Analysis of nascent transcription by one hour incubation with 1 mM of EU after 2 to 5 days of 4-OHT (-*Taf7* or -*Taf10*) or EtOH (Control) treatment (5-ethynyluridine (EU) labeling in *RT7* and *RT10* cells after 2 to 5 days of 4-OHT (-*Taf7* or -*Taf10*) or EtOH (Control) treatment. The time course of the experiments is indicated at the top. Representative pictures and quantification presented as violin plots are shown in (A) and (B), respectively. D2; two biological replicates (n=2), D3; n=3, D4; n=3, D5; n=2. **(C,D)** EU incorporation in control cells after 2 hours of treatment with Pol II transcription inhibitors (500 µm of triptolide (Trp) or 300 µM of flavopiridol (Flavo)). Representative pictures and quantification presented as violin plots are displayed in (C) and (D), respectively (n=1). Hundreds of measurements for individual nuclei were performed for the quantification, scale bar: 50 µm. Kruskal Wallis test: ns; not significant, *<0.05; **<0.01; ***<0.001. **(E)** Western blot analysis of the phosphorylated forms of RBP1 using an anti-RBP1 antibody from chromatin extracts of EtOH-treated *RT7* mESCs (Ctrl) and 4-OHT treated *RT7* (-*Taf7*) and *RT10* (-*Taf10*) mESCs after 3 days of treatment. The hyper-phosphorylated RPB1 (Pol II-O, active) and hypo-phosphorylated (Pol II-A) are indicated on the right. At the bottom anti-TAF6 western blot analysis as loading control.

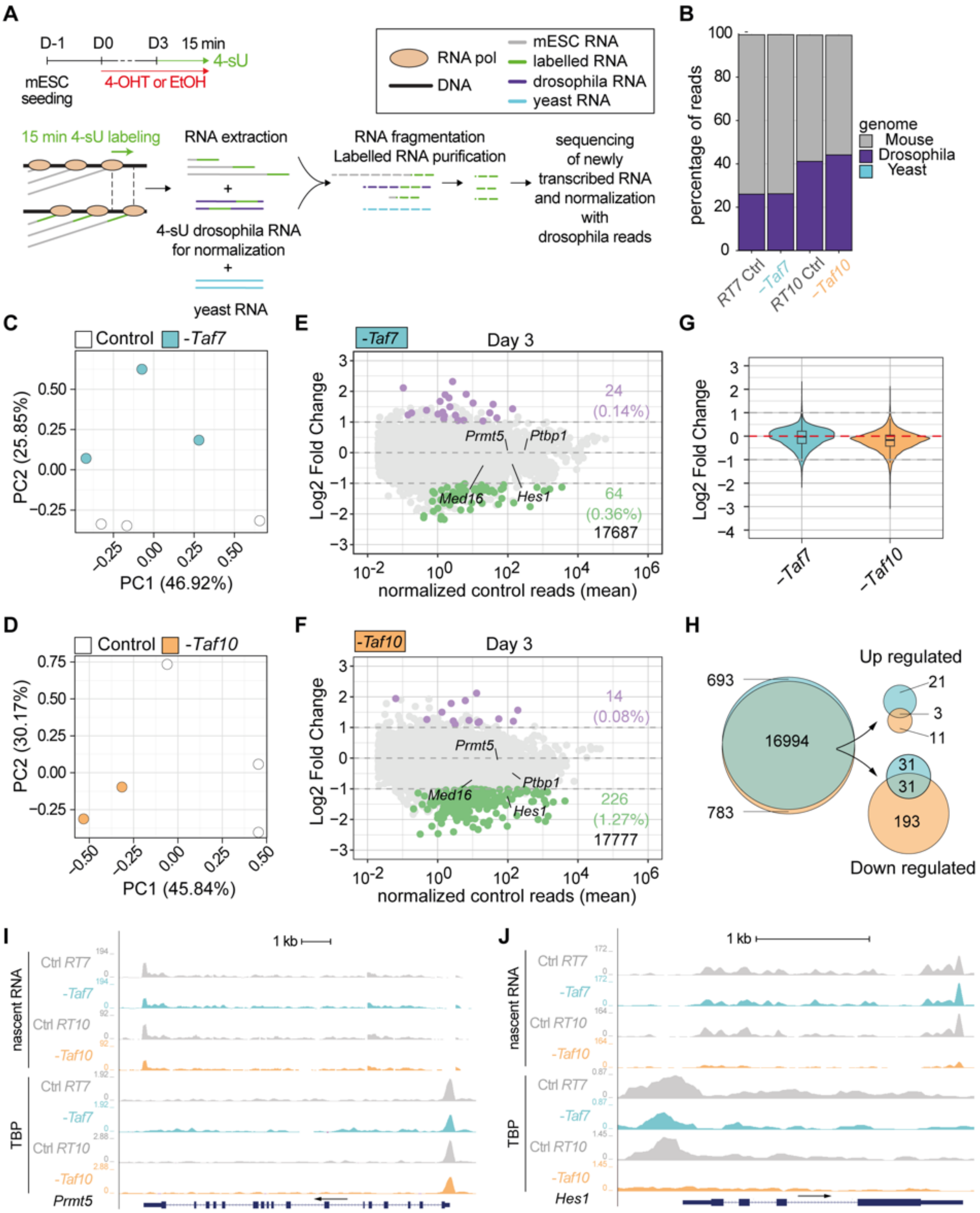


**Figure 6: TAF7 loss has a milder impact on RNA pol II gene transcription than TAF10 loss after 3 days of treatment. (A)** Time course and scheme of the nascent RNA sequencing. After three days of treatment with 4-OHT (-*Taf7* and -*Taf10* for mutant *RT7* and *RT10* mESC, respectively) or EtOH (Control), nascent transcripts were labeled with 4-sU during 15 minutes. 4-sU labelled RNA from *Drosophila* S2 cells were used as spike for normalization and unlabelled yeast RNA as a control for the purification of the nascent transcripts. **(B)** Percentage of mouse (grey), *Drosophila* (purple) and yeast (cyan) reads in the different datasets. **(C-D)** Principal component analysis (PCA) of the nascent transcriptomes of *RT7* (C) and *RT10* (D) control (white dot) and mutant (colored dot) mESCs. **(E,F)** MA plots of 4-sU-seq from D3 mutant versus control RT7 (E) and RT10 (F) with the number and percentage of significantly down- (green) or up-regulated (purple) protein coding transcripts displayed on the right and total transcripts detected at the bottom. A threshold of 20 normalized reads per gene length in kb was used to select actively expressed protein coding genes. Significance transcripts were filtered using an adjusted P value $\leq 0.05$ and an absolute Log2 Fold Change $\geq 1$. **(G)** Comparisons of global Log2 fold changes in TAF7- and TAF10-depleted mESCs. **(H)** Venn diagrams

analysis of all (left), up-regulated (top, right) and down-regulated (bottom, right) protein coding transcripts in TAF7- and TAF10-depleted mESCs. **(I,J)** UCSC genome browser views of nascent RNA (top) and TBP distribution (bottom) at *Prmt5* (I) and *Hes1* (J)loci between control (Ctrl *RT7* and Ctrl *RT10*), TAF7 depleted (-*Taf7*) and TAF10 depleted (-*Taf1*0) mESCs. Y axes indicate the genomic coverage. Arrows indicate direction of transcription.

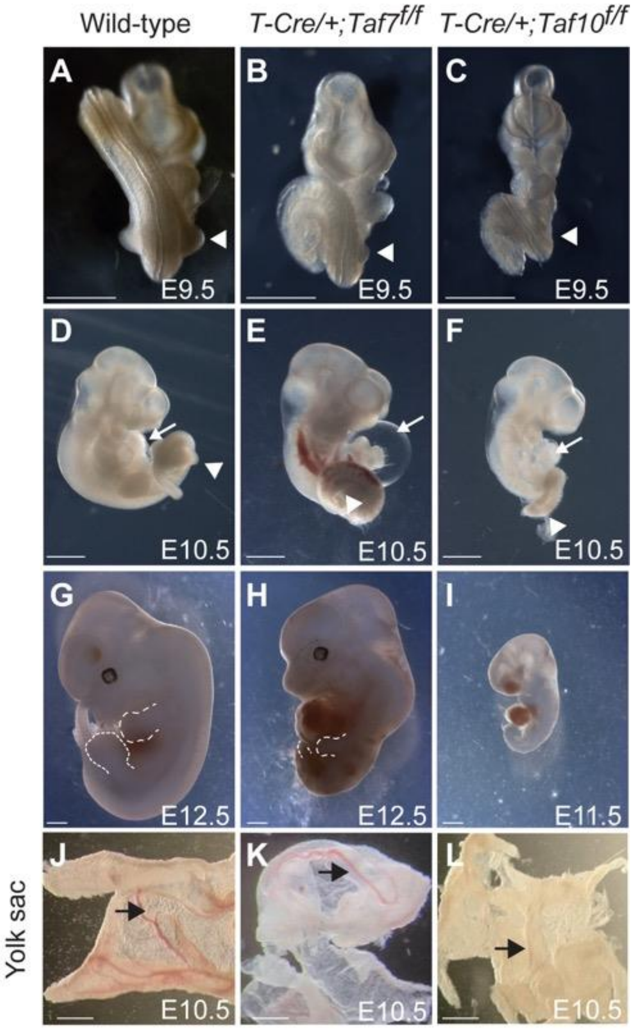


**Figure 7: Conditional deletion of *Taf7* or *Taf10* in the early mesoderm results in similar yet different phenotypes. (A-I)** Wholemount view of wild-type (A, D, G), Tg(T-Cre/+)*;Taf7$^{f/f}$* mutant (B, E, H) and Tg(T-Cre/+)*;Taf10$^{f/f}$* mutant (C, F, I) embryos at E9.5 (A-C, n>4), E10.5 (E-F, n>4), and E12.5 (G, H, n>2). As no Tg(T-Cre/+)*;Taf10$^{f/f}$* mutant embryos could be recovered at E12.5, a E11.5 Tg(T-Cre/+)*;Taf10$^{f/f}$* mutant embryo is shown (I). White arrowheads (A-C) indicate the position of the forelimbs. Dashed lines (G, H) indicate the limb buds. White arrows (D-F) indicate the heart. **(J-L)** Wholemount views of wild-type (J), Tg(T-Cre/+)*;Taf7$^{f/f}$* (K) and Tg(T-Cre/+)*;Taf10$^{f/f}$* (L) yolk sacs embryos are showed (n>3). Black arrows indicate the presence of blood vessels. Scale bars; 1 mm.
