## supplementart figures_plus_legend for "RNA polymerase II transcription with partially assembled TFIID complexes"

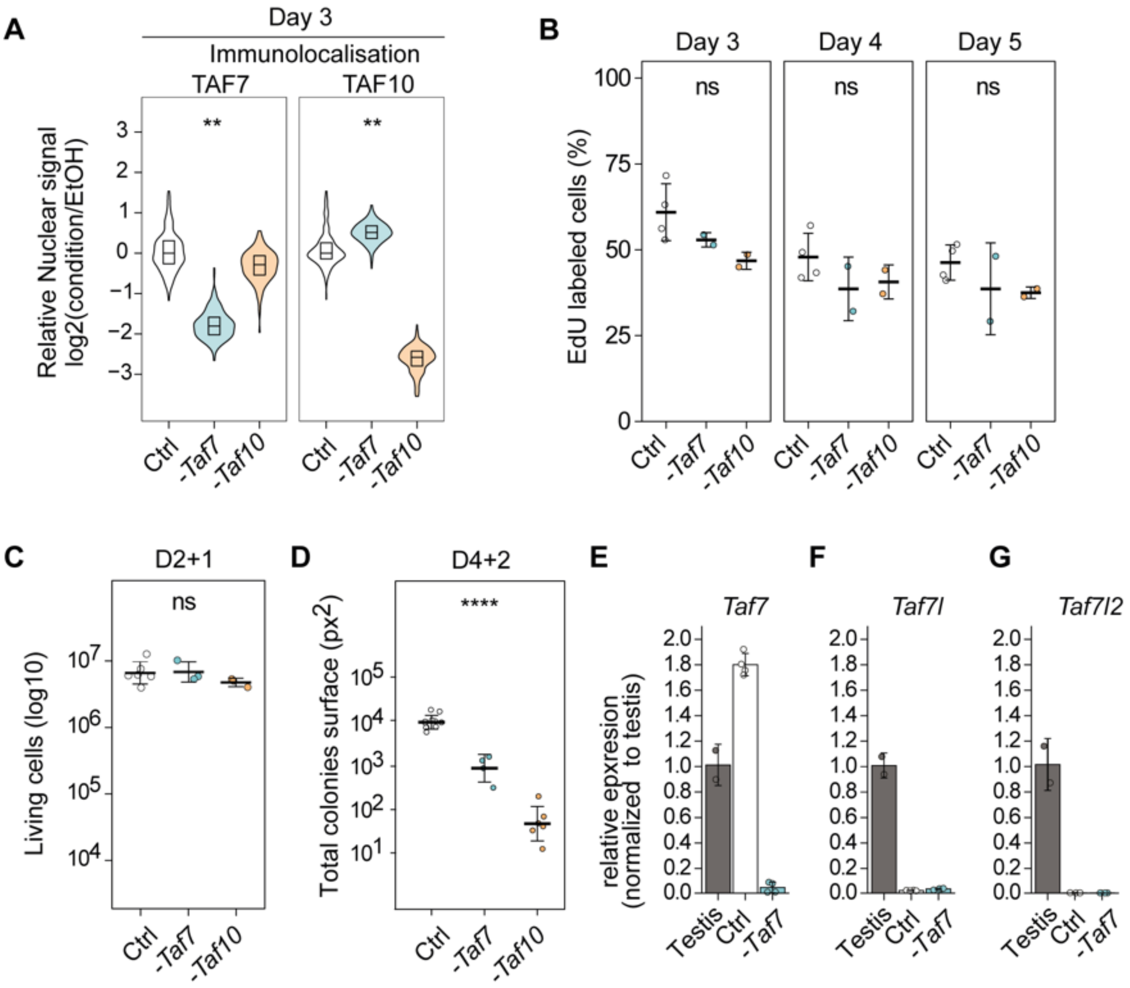


**Sup Figure 1: TAF7 depletion in mESCs results in a milder phenotype compared to TAF10 depletion. (A)** Nuclear quantification of anti-TAF7 and anti-TAF10 immunolocalization signal in control condition (Ctrl) or after *Taf7* (*-Taf7)* or *Taf10* deletion (*-Taf10)* after 3 days of treatment. **(B)** Percentage of cells in S phase assessed by 5-ethynyldeoxyuridine (EdU) staining at 3, 4, and 5 days of treatment. For D3 to D5, Ctrl; n = 4, -*Taf7;* n = 2, -*Taf10*; n = 2 biological replicates. **(C,D)** Quantifications of the ability of the *RT7* or *RT10* depleted cells to growth after cell passage. Living Trypan blue negative cell numbers after cell passage at D2 and analyzed at D3 (D2+1). Ctrl; n = 6, -*Taf7;* n = 3, -*Taf10*; n = 3 biological replicates (C). Total colonies surfaces estimated by Crystal violet staining after cell passage at D4 and analyzed at D6 (D4+2). Ctrl; n = 10, -*Taf7;* n = 4, -*Taf10*; n = 6 biological replicates (D). For all panels, the control condition (Ctrl) corresponds to all *RT7* and *RT10* cells treated with EtOH. **(E-G)** RT-qPCR analysis of *Taf7, Taf7l* and *Taf7l2* expression from control (Ctrl), mutant *(-Taf7) RT7* mESCs at day 4 and wild type mouse testis RNA. The RNA polymerase III transcribed gene *Rn7sk* was used as reference gene and the data were normalized to the testis. n=2 biological x 2 technical replicates, except for testis; n= 2 technical replicates. Means ± standard deviation are shown. (A-C) Kruskal–Wallis test: ns; not significant, * <0.05; **<0.01; *** <0.001; **** <0.0001.

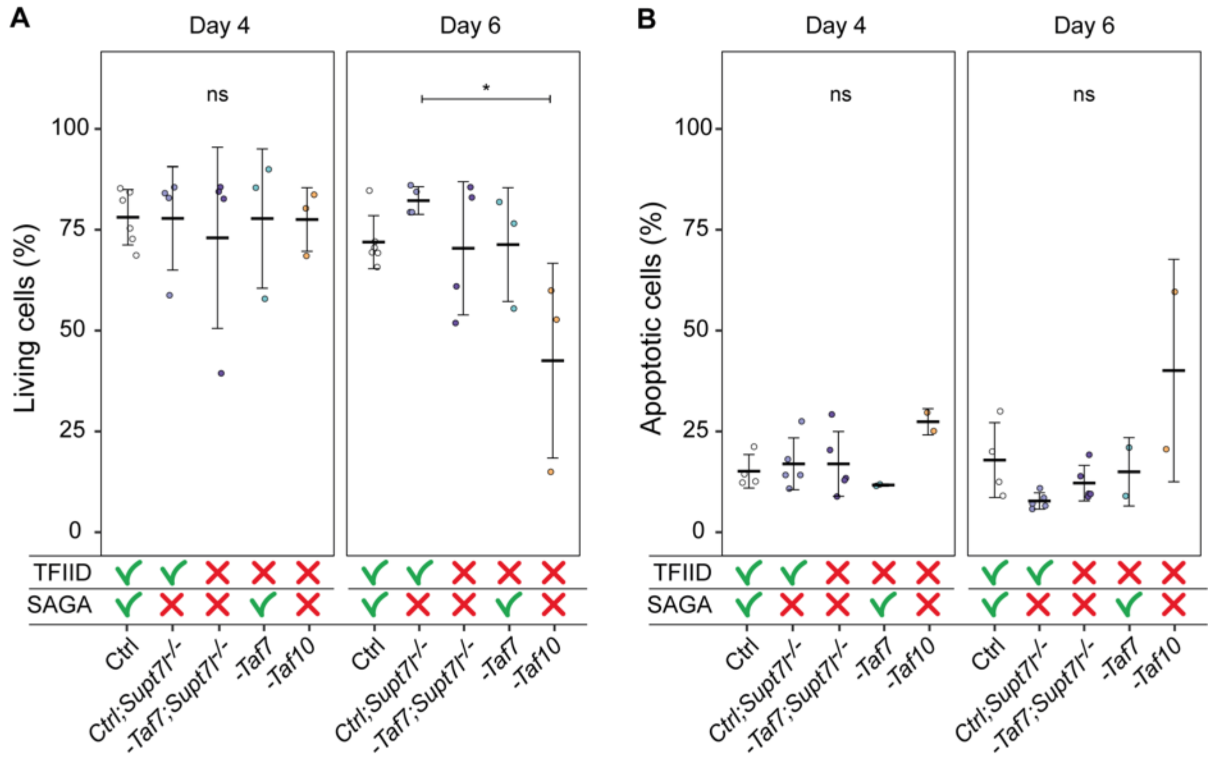


**Figure S2: *Supt7l* loss of function in TAF7 depleted mESCs does no impact cell viability. (A)** Percentage of living mESCs analyzed by Trypan blue staining after 4 and 6 days of treatment. Ctrl; n = 6, Ctrl;*Supt7l*$^{-/-}$; n = 4, -*Taf7;Supt7l*$^{-/-}$; n = 4, -*Taf7;* n = 3, -*Taf10*; n = 3 biological replicates per day. **(B)** Percentage of apoptotic (Annexin V positive and propidium iodide negative) mESCs after 4 and 6 days of treatment. Ctrl; n = 4, Ctrl;*Supt7l*$^{-/-}$; n = 5, -*Taf7;Supt7l*$^{-/-}$; n = 5, -*Taf7;* n = 2, -*Taf10*; n = 2 biological replicates per day. The impact on TFIID and SAGA assembly is indicated at the bottom. Kruskal–Wallis test followed by a post hoc Dunn test if significant: ns; not significant, * <0.05.

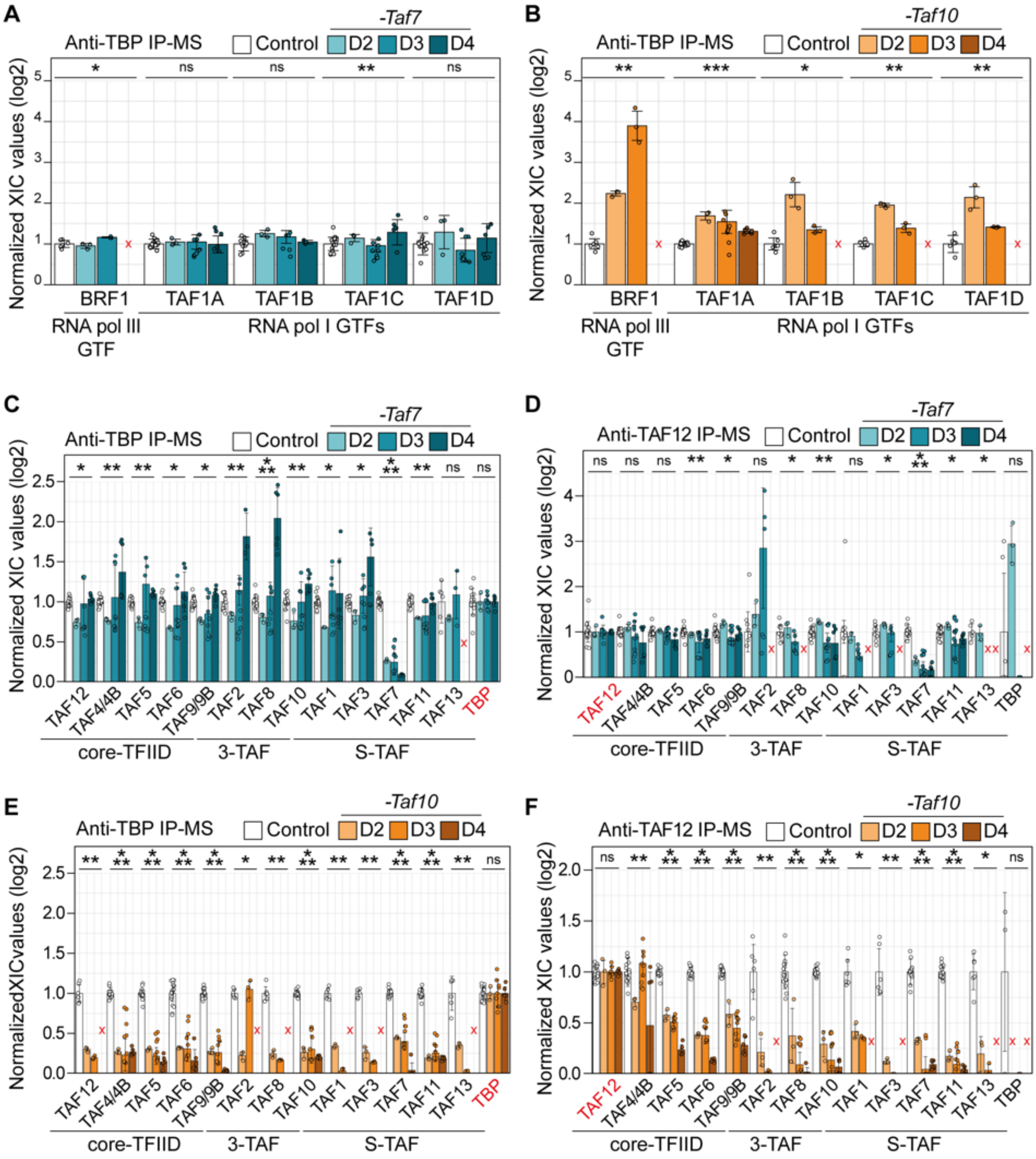


**Figure S3: Different effects on holo-TFIID assembly in TAF7- and TAF10-depleted mESCs. (A-F)** Normalized relative XIC values for RNA polymerase I and III GTFs subunits (A, B) and TFIID subunits (C-F) after anti-TBP (A-C,E) or anti-TAF12 (E,F) IP-MS. For each IP-MS, the XIC values of each subunit were first normalized to the mean XIC of the bait protein and then normalized to the mean of the respective control condition (EtOH treated *RT7* for mutant -*Taf7*, EtOH treated *RT10* for mutant -*Taf10)*. Analyses were performed on day 2, day 3, and day 4 of treatment. Red crosses indicate proteins not detected in the control condition. Day 2; n = 1 biological replicate x 3 measures, D3; 3x3, D4; 2x3. Means ± standard deviation are shown. Kruskal-Wallis test. ns; not significant, * <0.05; **<0.01; *** <0.001.

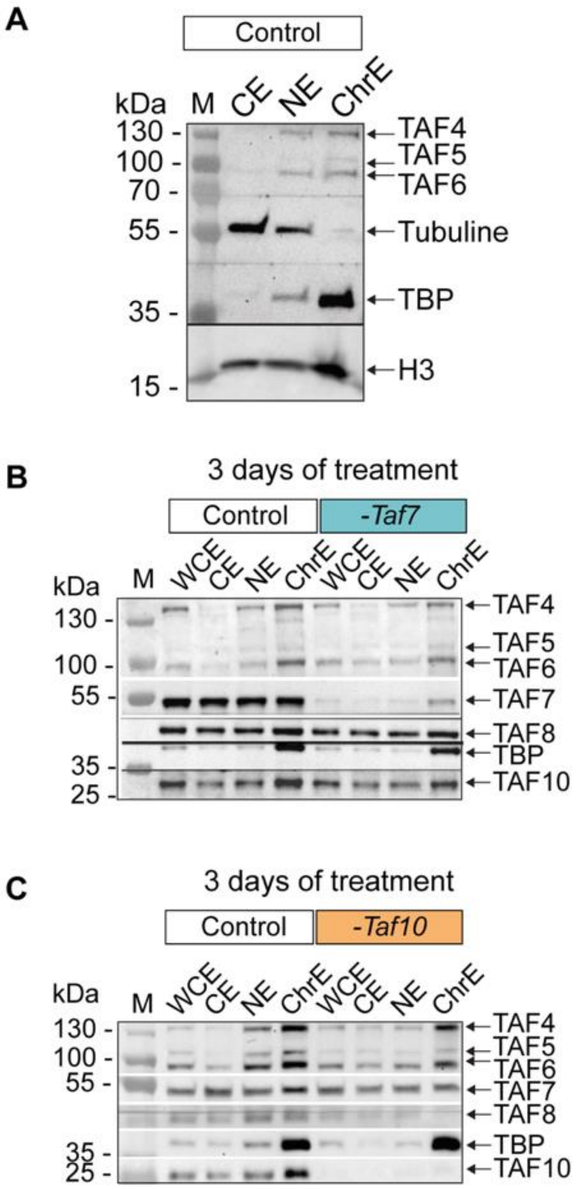


**Figure S4: TBP recruitment on the chromatin of TAF7- and TAF10-depleted mESCs**. **(A)** Western blot analysis on cytoplasm-enriched extract (CE), nuclear-enriched extract (NE), and chromatin-enriched extract (ChrE) obtained from untreated mESCs. The position of the different proteins probed is indicated on the right and the molecular weights on the left. **(B,C)** Western blot analyses of whole cell extract (WCE), CE, NE and ChrE from 4-OHT treated *RT7* (*-Taf7*, C) and 4-OHT treated *RT10* (*-Taf10*, D) depleted mESCs and their respective controls (Control, EtOH treated cells) at day 3. The position of the different proteins probed is indicated on the right and the molecular weights (M) on the left.

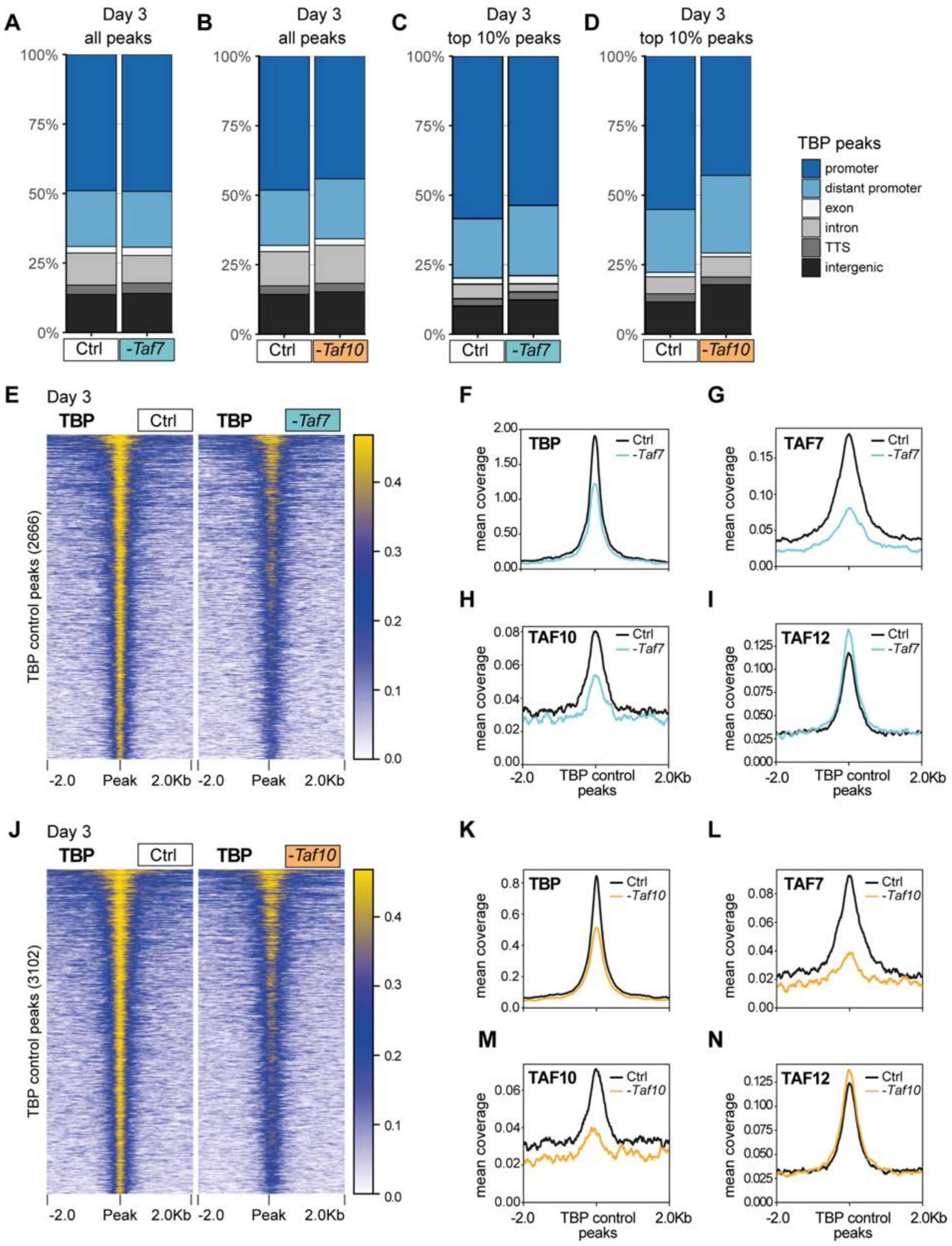


**Figure S5: TBP recruitment on the chromatin of TAF7- and TAF10-depleted mESCs. (A-D)** All (A,B) and top10% (C,D) TBP CUT&RUN peak annotation in *RT7* (A,C) and *RT10* (B,D) control (Ctrl) and depleted (-*Taf7* and -*Taf10*) mESCs. The legend of the annotations is indicated on the right. **(E)** Heatmap showing the TBP distribution at the position of the TBP peaks (± 2 kb) detected in the control condition, in control (Ctrl) and mutant *RT7* (-*Taf7*) mESCs at day 3. **(F-I)** Mean profile of TBP (F), TAF7 (G), TAF10 (H) and TAF12 (I) protein accumulation at the location of the TBP control peaks, in control (black line) and in TAF7-depleted (blue line) mESCs. **(J)** Heatmap showing the TBP distribution at the position of the TBP peaks (± 2 kb) detected in the control condition, in control (Ctrl) and mutant *RT10* (-*Taf10*) mESCs at day3. **(K-N)** Mean profile of TBP (F), TAF7 (G), TAF10 (H) and TAF12 (I) protein accumulation at the location of the TBP control peaks, in control (black line) and in TAF10-depleted (orange line) mESCs.

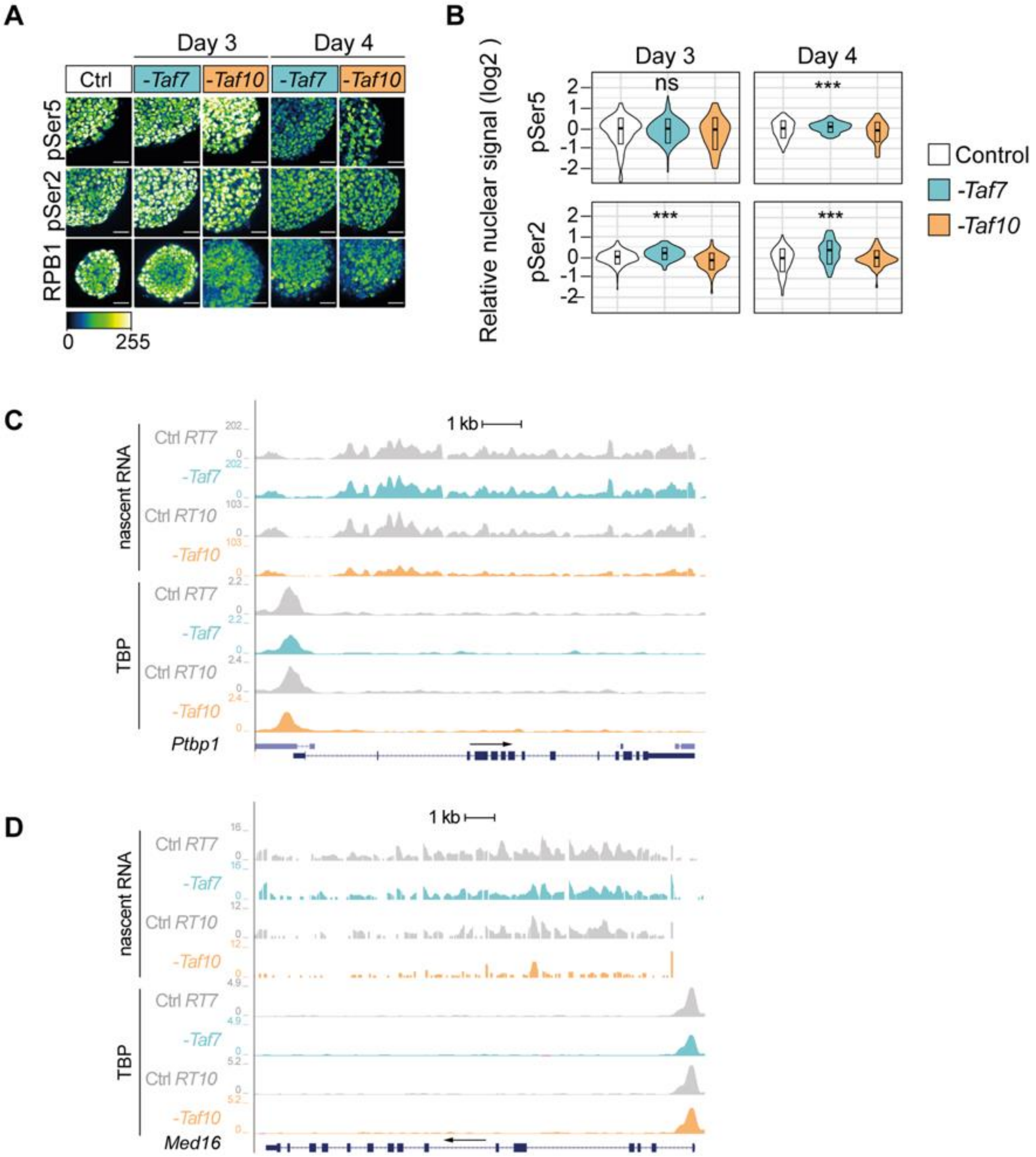


**Figure S6: RNA Pol II global activity is not affected in TAF7- and TAF10-depleted mESCs. (A)** Representative views of immunofluorescence using anti-RPB1 (bottom), anti-RPB1$^{pSer2}$ (middle) and anti-RPB1$^{pSer5}$ (top) antibodies on *RT7* and *RT10* mESCs treated 3 or 4 days with 4-OHT (-*Taf7* or -*Taf10*) and with EtOH (Ctrl). Color scale (Green Fire Blue LUT scale) is indicated at the bottom of the panel, scale bar: 15 µm. **(B)** Quantifications of RPB1$^{pSer5}$ and RPB1$^{pSer2}$ nuclear signal represented as violin plots. D3 and D4; n=1 biological replicate. Hundreds of measurements for individual nuclei were performed for the quantification, scale bar: 50 µm. Kruskal-Wallis test. ns; not significant, * <0.05; **<0.01; *** <0.001. **(C,D)** UCSC genome browser views of nascent RNA (top) and TBP distribution (bottom) at *Ptbp1* (C) and *Med16* (D) loci between control (Ctrl *RT7* and Ctrl *RT10*), TAF7 depleted (-*Taf7*) and TAF10 depleted (-*Taf1*0) mESCs. Y axes indicate the genomic coverage. Arrows indicate direction of transcription.

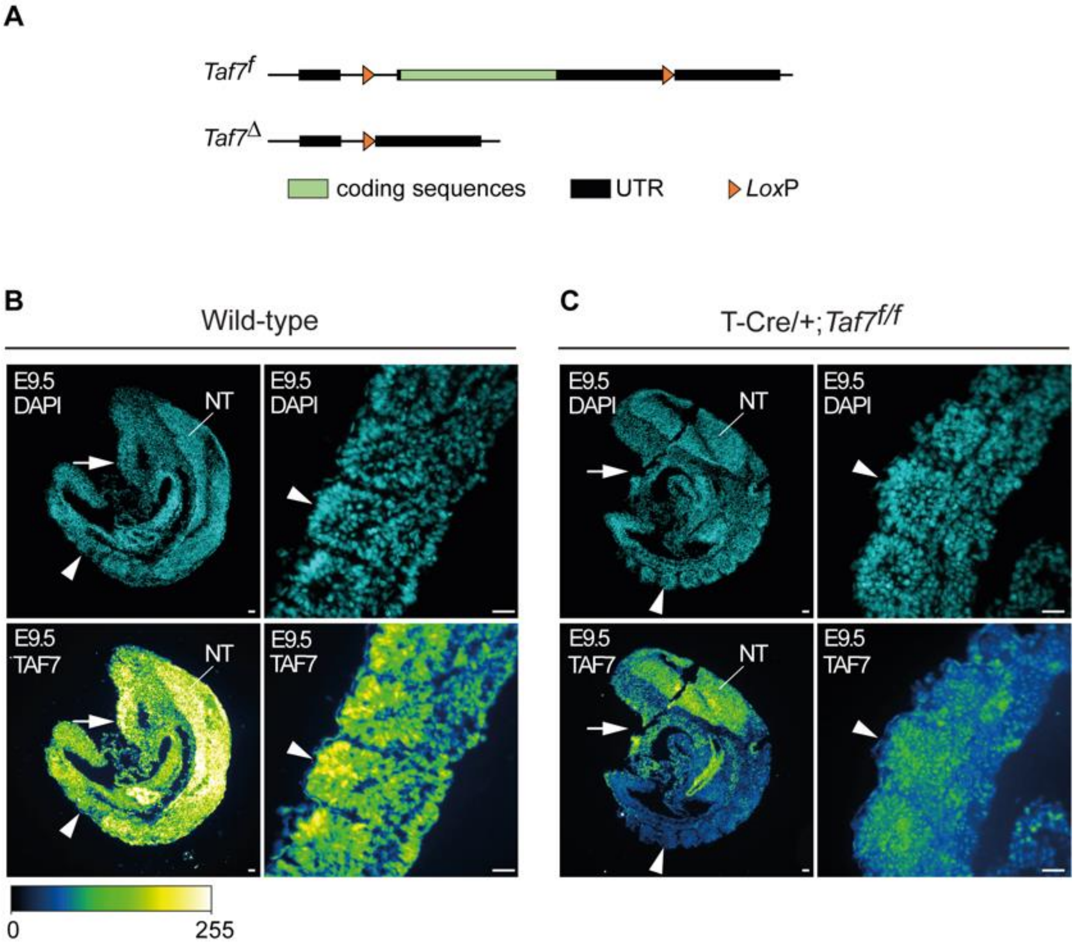


**Sup Figure 7: Efficient TAF7 depletion in the paraxial mesoderm of embryonic day (E) 9.5 Tg(T-Cre/+)*;Taf7$^{f/f}$* mutant embryos. (A)** Map of the *Taf7* conditional (*Taf7$^{f}$*) and deleted (*Taf7$^{\Delta}$*) alleles after CRE recombination. UTRs and ORF are indicated in black and green boxes, respectively. *Lox*P sites are indicated by orange arrowheads. **(B-C)** DAPI counterstained anti-TAF7 immunofluorescence on sagittal sections from E9.5 wild-type (B) and Tg(T-Cre/+)*;Taf7$^{f/f}$ mutant* (C) embryos. In Tg(T-Cre/+) embryos, CRE is active in progenitors that contribute to mesoderm cells posterior to the heart (white arrow), including the paraxial mesoderm (unsegmented presomitic mesoderm and somites (arrowhead)) but not in the neural tube (NT). Color scale (Green Fire Blue LUT scale) is indicated at the bottom (n=2), scale bar: 30 µm.
